# Neural circuit-selective, multiplexed pharmacological targeting of prefrontal cortex-projecting locus coeruleus neurons drives antinociception

**DOI:** 10.1101/2024.06.08.598059

**Authors:** Chao-Cheng Kuo, Jordan G. McCall

**Affiliations:** Department of Anesthesiology, Washington University in St. Louis, St. Louis, MO, USA; Center for Clinical Pharmacology, University of Health Sciences and Pharmacy in St. Louis and Washington University School of Medicine, St. Louis, MO, USA; Washington University Pain Center, Washington University in St. Louis, St. Louis, MO, USA

## Abstract

Selective manipulation of neural circuits using optogenetics and chemogenetics holds great translational potential but requires genetic access to neurons. Here, we demonstrate a general framework for identifying genetic tool-independent, pharmacological strategies for neural circuit-selective modulation. We developed an economically accessible calcium imaging-based approach for large-scale pharmacological scans of endogenous receptor-mediated neural activity. As a testbed for this approach, we used the mouse locus coeruleus due to the combination of its widespread, modular efferent neural circuitry and its wide variety of endogenously expressed GPCRs. Using machine learning-based action potential deconvolution and retrograde tracing, we identified an agonist cocktail that selectively inhibits medial prefrontal cortex-projecting locus coeruleus neurons. *In vivo*, this cocktail produces synergistic antinociception, consistent with selective pharmacological blunting of this neural circuit. This framework has broad utility for selective targeting of other neural circuits under different physiological and pathological states, facilitating non-genetic translational applications arising from cell type-selective discoveries.

## Introduction

The development of cell type-selective neuroscience tools such as optogenetics and chemogenetics has enabled unprecedented dissection of neural circuitry^1–4^. These advanced tools have dramatically improved our understanding of neural circuit function while also providing translational insight for strategies to alleviate neurological and neuropsychiatric disorders. However, these approaches rely exclusively on genetic targeting to particular cell populations^5–8^, largely hindering immediate clinical translation^9–12^. To overcome this limitation, we developed a novel, genetic tool-independent approach for neural circuit-selective manipulation through multiple pharmacological strategies.

To do so, we leveraged the endogenous properties of the mouse locus coeruleus noradrenergic (LC-NE) system. LC-NE neurons, located in the dorsal pons, are the main source of NE for the mammalian central nervous system and have nearly ubiquitous efferent circuitry throughout the forebrain and spinal cord. Consequently, these neurons participate in a variety of brain functions including cognitive control, salience detection, memory, arousal, stress processing, and pain regulation^13–18^. Furthermore, LC-NE neurons express a diverse population of G-protein coupled receptors (GPCRs), leading to complex physiological regulation by endogenous GPCR ligands^19–22^. For example, optogenetic excitation of corticotropin-releasing factor (CRF) secreting axons in LC drive anxiety-related avoidance behaviors, while pharmacological activation of orexin receptors (OXARs) and mu-opioid receptor (MOR) in LC facilitate synaptic plasticity in hippocampus and interfere the performance in cognitive function, respectively^23–25^. These various LC GPCRs are ideal for pharmacological targeting. Despite historically being considered a homogenous structure, recent work has emerged demonstrating anatomically defined modular organization of efferent LC projections during distinct behaviors^26–30^. This heterogeneity is common among central neuromodulatory systems^31–33^ and makes the LC an excellent testbed for a neural circuit-selective control by additive, synergistic, or competitive pharmacology across its efferent modules. However, challenges arise from the lack of understanding about how LC-NE GPCRs help rapidly reallocate of neural activity across LC modules in response to environmental stimuli. The consistent response of LC-NE neurons to noxious nociceptive stimuli has been used as a criterion in single unit recordings^34,35^, the augmented expression of neural excitation markers such as cfos and pERK in LC were found during acute inflammatory pain and during the development of chronic pain^36–38^. Furthermore, pharmacological LC inhibition increases paw withdrawal threshold to noixious stimuli^39–41^. In a separate and contemporaneous study, we show that the genetic knockout of mu opioid receptor (MOR) in the LC increased baseline mechanical and thermal sensitivity^42^. LC modularity is clear and apparent in pain regulation^28–30,42–52^. Notably, the LC-NE system bidirectionally modulates nociception. Efferent spinal cord projections mediate descending analgesia via spinal adrenergic receptors, while activation of LC efferents targeting medial prefrontal cortex (mPFC) is pronociceptive and hyperalgesic after injury^28,42,43^. This LC-mPFC efferent projection is also involved in many cognitive functions such as attention, decision making, fear recall and stress regulation through a dense noradrenergic innervation^15,23,27,43,53,54^. Given that LC-mPFC projections are crucial for regulating cognition and nociception, it is reasonable that differential modulation from distinct GPCRs across efferent-defined LC modules may help achieve rapid, flexible shifts in LC-NE neural activity upon detection of nociceptive stimuli.

Together, the functional separation of the LC-mPFC module from LC-NE neurons that do not innervate the mPFC could be an ideal target for circuit-selective antinociception through multiple GPCR-mediated strategies. To identify circuit-selective pharmacological strategies for targeting the LC-mPFC projection, we developed an accessible *ex vivo* calcium imaging-based approach for large-scale pharmacological scans of endogenous receptor-mediated neural activity. Our method takes advantage of the newly developed ultra-fast kinetics and high sensitivity of the latest generation of genetically-encoded calcium indicators (i.e., GCaMP8f^55^). This assay enabled efficient functional screening of ligands for eighteen different GPCRs expressed in the LC^19,22^. Importantly, this approach can be combined with anatomical tracing to identify pharmacological action at discrete neural circuits, enabling quick identification of circuit-selective receptor-mediated responses. We demonstrate this efficiency using retrograde labeling of LC -mPFC neurons. Cell type-selective genetic approaches have shown this LC-mPFC neural circuit to promote hypersensitivity during nociception^28^. Our method identified that both mu opioid receptor (MOR) and 5HT1a receptor (5HTR1a) agonists more robustly inhibit LC-mPFC neurons compared to other LC neurons. Likewise, mACh1 receptor agonism leads to greater excitation of non-mPFC-projecting LC neurons. To translate these findings *in vivo*, we found local infusion of a cocktail of these slice imaging-identified agonists provides synergistic thermal antinociception – a finding consistent with shifting neural activity away from the LC-mPFC projection through circuit-selective pharmacology. Moreover, genetic deletion of MOR in the LC-mPFC module demonstrates that interaction between LC modules is required for the pharmacological synergy. Altogether, we introduce a new, easily accessible pipeline of techniques to identify non-genetic, circuit-selective pharmacological approaches for neural modulation.

## Results

### Deconvolution of spontaneous firing of LC-NE neurons

While most evidence of modular LC function relies on differential cell type-selective control of efferent circuity with optogenetics or chemogenetics^27–30,54,56^, endogenous control of these modules may be mediated by neuromodulators acting at the diverse array of GPCRs expressed on these neurons. To test whether GPCR-mediated regulation of LC-NE neurons can give rise to this modular function, we selectively expressed GCaMP8f in LC-NE neurons by injecting AAV1-DIO-GCaMP8f into the LC of dopamine beta hydroxylase-Cre (*Dbh*-Cre) mice (**Fig. 1A, B**). Consistent with our previous findings, *ex vivo* cell-attached recordings under normal slice conditions showed most LC-NE neurons fire spontaneous action potentials in the 1-3 Hz range^42,57^ (**Fig. 1C**). Interestingly, we found robust rhythmic fluctuations in the GCaMP8f calcium signal from LC-NE neurons. We then extracted the spatiotemporal information of individual regions of interest (ROI) from calcium imaging recordings taken on our typical epifluorescent slice electrophysiology microscope (**Fig. 1B, S1A;** See Methods). Simultaneous cell-attached recordings demonstrate a strong relationship between spontaneous action potential firing and calcium fluctuations (**Fig. 1D**), suggesting the possibility of extracting LC action potentials from the calcium signal alone^58^. To do so, we adopted CASCADE, a machine learning-based algorithm, to precisely deconvolute individual action potential spikes from the calcium waveforms^59,60^. Using approximately 6 hours of simultaneous cell-attached and calcium imaging recordings we were able to fit a network capable of predicting the temporal location of individual action potentials. We held back 15% of total learning materials for model evaluation (**Fig. 1E**). We then converted calcium signals to the likelihood of spike occurrence, and peaks representing a higher probability of action potential generation were detected and aligned with the electrophysiological ground truth showing clearly identified action potential firing from simultaneous cell-attached recordings (**Fig. 1F, S1B**). Our model displays an excellent predictive accuracy (99.2%) for LC-NE neurons with spontaneous firing rates <4 Hz and this value only dropped to 82.1% when cells fired over 5 Hz. Together, these results provide sufficient deconvolution of action potential firing from LC-NE neurons under normal slice conditions (**Fig. 1C, F**). Further analysis shows most of the decreased predictive accuracy came from missing existing spikes rather than erroneous predictions. This issue was exaggerated in ROIs with higher firing rates due to limits on temporal resolution from the imaging acquisition and smaller calcium fluctuations that occur during higher firing rates in LC-NE neurons (**Fig. S2A-C**). This can also be seen in the temporal difference between the predicted spike and ground truth (**Fig. S2D**). To address the issue of repeated ROIs in both somatic and dendritic components from the same cell, a cross-correlation matrix was made based on the traces of spike likelihood across extracted ROIs (**Fig. S3A**). Interestingly, the distribution of these *ex vivo* correlation coefficients exhibit a high similarity to those from *in vivo* single unit recordings previously reported by Totah and colleagues^61^. These synchronous dynamics could arise from the electrical coupling through dendritic gap junctions between LC-NE neurons, whereas the negative relationship could be driven by alpha2-adrenergic receptor-mediated inhibition from neighboring cells^58,62–65^. Therefore, we used a correlation coefficient cutoff threshold between ROI pairs due to the transient increase in complete synchronicity in ROI pairs with higher coefficients, pairs with a correlation coefficient greater than 0.3 are considered together as a single electrical compartment (**Fig. S3B**). Together, spike deconvolution from *ex vivo* calcium imaging enables efficient recording of individual neuronal activity across the whole LC population.

**Figure 1.**
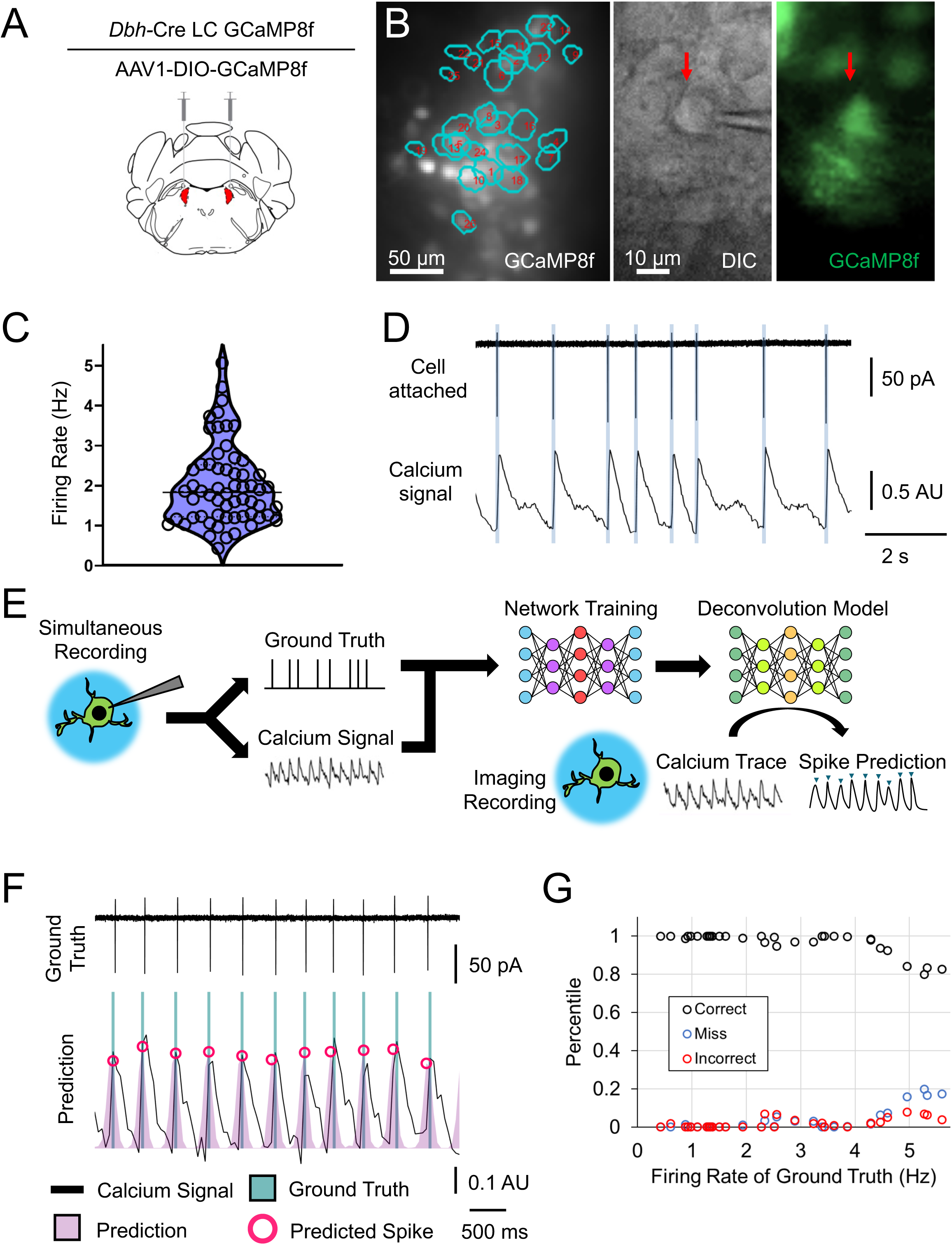
Deconvolution of individual spikes in LC-NE neurons using calcium imaging. (A) A schematic cartoon illustrating the viral strategy for selective expression of GCaMP8f in LC. (B) Left: A DIC image under 40x objective lens showing the spatial information of extracted ROIs; Middle & Right: Live DIC and fluorescent images, respectively, showing the simultaneous electrophysiological and imaging recordings. (C) A scatter plot that displays the distribution of firing rate in LC-NE neurons under normal slice preparation by cell-attached recordings. (D) An example of aligned simultaneous cell-attached and calcium imaging recording, note the complete coincidence between action potentials and peaks of calcium waveform. (E) A schematic cartoon depicting the machine learning-based network training and spike deconvolution using trained model. (F) An example of model evaluation using the simultaneous recording on a LC-NE neuron with around 2 Hz spontaneous firing, note the perfect inference of action potential firing. (G) A plot showing the predictive accuracy for spike deconvolution, excellent accuracy remains with lower firing rate and drops along the firing rate of recorded cells. Proportions of correct and failed predictions are denoted by circle in different colors (correct: black; miss: blue; incorrect: red).

### Multiplexed pharmacological scan of LC-NE neurons

If afferent control through GPCRs underlies some aspect of LC efferent modularity, we must first identify how neurons respond to such receptor activation (**Fig. 2A**). With the ability to monitor LC action potential activity across the majority of the structure established, we next sought to determine the response of individual LC neurons to various GPCR ligands. Following transcriptomic and translatomic results for LC-NE neurons from previous studies^19–22^, we tested forty agonists targeting different LC-expressed GPCRs and selected eighteen of them based on an initial screen for changes in activity. We further divided these agonists into three groups according to their G protein coupling or functional implications (**Table 1**). We then applied agonists from these three groups to brain slices in a mostly randomized order at subsaturating doses with complete functional washouts between ligands (**Fig. 2B**). A few exceptions to the randomization were experimentally necessary. In particular, one agonist in each group caused irreversible effects and, as such, was always applied last (**Table 1**). Additionally, LC-NE neurons co-express muscarinic acetylcholine receptors 1 and 3 (mAChR1/3), but not 2 and 4 (mAChR2/4). However, there is no selective mAChR3 agonist^66^. Therefore, we applied pirenzepine, an antagonist for mAChR1, along with the nonselective muscarinic acetylcholine receptor agonist, muscarine, to obtain selective activation of mAChR3. Therefore, we necessarily always applied this combination after mAChR1 activation. In preliminary cell-attached studies, we identified doses for all agonists that caused detectable, but washable effects (**Fig. S4**). We then performed the slice calcium imaging during baseline and agonist application for each agonist and calculated the pharmacological effect by the change in firing rate from each ligand. In a subset of trials, we bath applied clonidine, a potent alpha2-adrenergic receptor agonist, after the entire pharmacological scan to functionally verify the cellular identity of LC-NE neurons beyond the genetic constraint of GCaMP8f expression (**Fig. 2B**). Only one out of 54 ROI was excluded as not inhibited by clonidine (**Fig. S5A&B**) Cell-attached and calcium imaging recordings showed strikingly similar means and standard deviations in response to the 200nM DAMGO, further demonstrating the reliability of the imaging approach. Our slice imaging approach substantially increased experimental throughput compared to cell-attached recordings (**Fig. 2C&D**, 33 cells from 18 slices vs. 108 ROIs from 5 slices) and enabled identification of LC neurons with distinct DAMGO-mediated responses. The increased sampling from calcium imaging revealed more LC neurons completely silenced by DAMGO as well as small number that increased firing during DAMGO application, consistent with prior literature^67–69^ (**Fig. 2C&D, S5C**). In general, DAMGO inhibited most LC-NE neurons but some variability could be caused by differential expression of functional MOR, intra-cellular signaling molecules, and/or G protein-gated inwardly rectifying K^+^ (GIRK) channel across the LC^70,71^, and showing a weak relationship (R^2^ = 0.2165) with baseline neural activity (**Fig. S5C**). Using this approach we further tested a comprehensive profile of pharmacological responses to agonists targeting different GPCRs native to LC-NE neurons (**Fig. S6**). As with MOR agonism, many agonists drove non-homogeneous effects offering direct evidence for the GPCR-mediated modules in the LC (**Fig. 2E-G**). Although it is possible the pharmacological response could be partially underestimated due to the greater predicted misses in cells with higher firing rate, the variability of agonist-induced responses was still capable of identifying increased firing rates. **Table 2** lists the detailed statistics for results of multiplexed pharmacological scan in LC-NE neurons. Most of these responses appear to be cell-autonomous responses as repeating the scan for Group 1 agonists in the presence of synaptic blockers (5mM kynurenic acid, 1μM strychnine and 100μM picrotoxin) left most results unaltered. We did, however, find that pre-administration of synaptic blockers abolished the excitatory effect from the alpha1-adrenergic receptor agonist phenylephrine. This blunting suggests alpha1-adrenergic receptor-mediated excitation of the LC is largely presynaptic (**Fig. 2H, S7**) which is largely consistent with recent electron microscopy results^72^.

**Figure 2.**
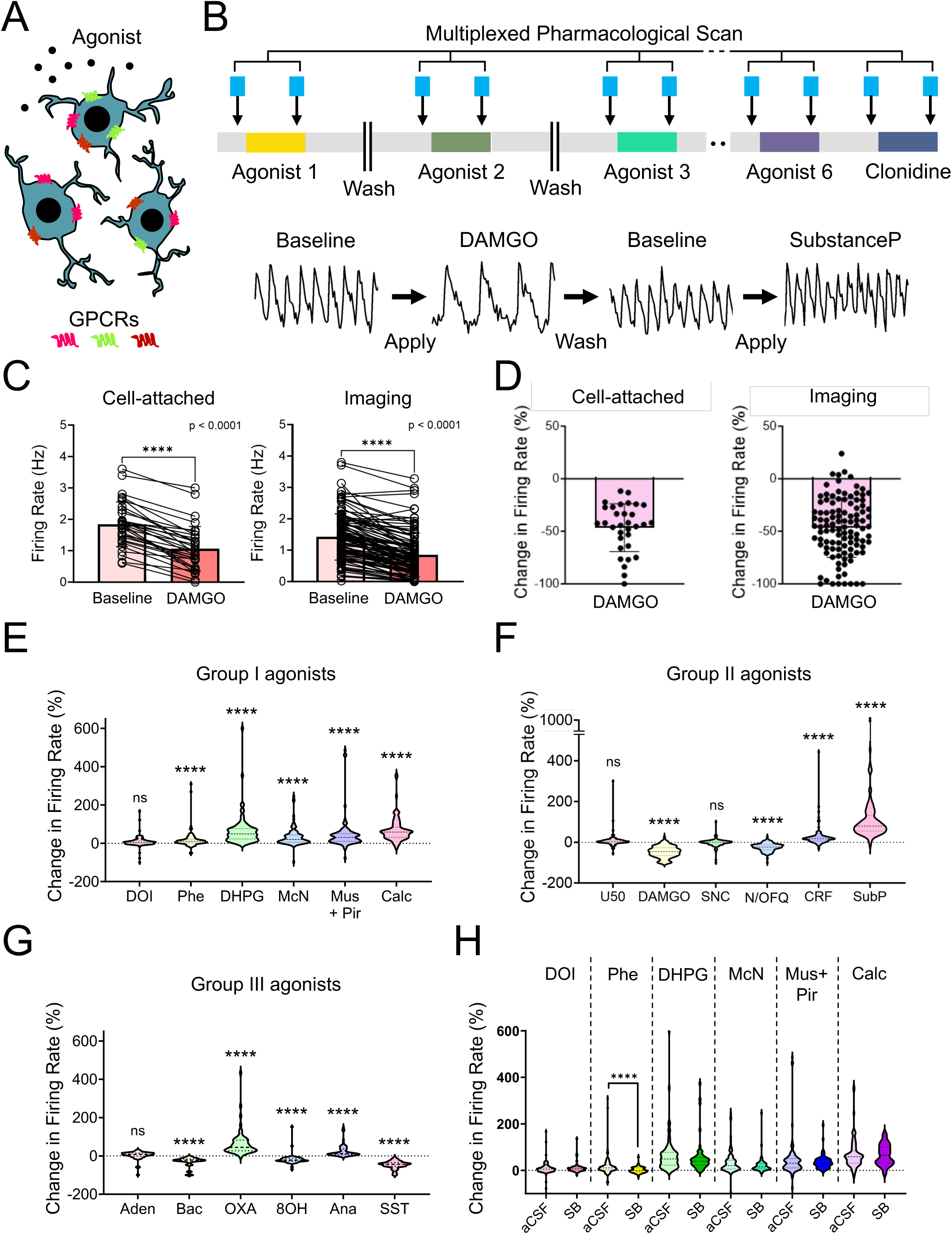
*Ex vivo* multiplexed pharmacological scan of GPCRs in LC. (A) A schematic cartoon displaying differential expression of multiple GPCR in LC-NE neurons. (B) Top, a schematic diagram illustrating the protocol of pharmacological scan. Bottom, representative calcium traces from an ROI showing the pharmacological effects upon application of DAMGO and Substance P, agonists targeting MOR and Neurokinin-1 receptor (NK1R), respectively. Note the stable calcium fluctuations among the baseline traces representing a complete wash between pharmacological effects. (C) Firing rate plots showing the pharmacological effects of DAMGO using cell-attached (left) and imaging (right) recordings. Cell-attached: Paired-t test, t = 11.99, ****p<0.0001. Imaging: Wilcoxon matched-pairs signed rank test, W = -5747, ****p<0.0001. (D) Plots showing the percentile of changes in firing rate from data shown in C (left: cell-attached recording, right: imaging recording). (E-G) Summarized results of changes in firing rate led by pharmacological activations of GPCRs in LC-NE across three groups of GPCR agonists. Repeated measures two-way ANOVA followed by Bonferroni test, ****p<0.0001, ns = not significant, please see **Table 2** for detailed statistics. (H) A plot demonstrating the examination of presynaptic modulation from GPCRs targeted by Group I agonists, only phenylephrine, an agonist targeting alpha1-adrenergic receptor, shows a presynaptic modulation as the significantly different pharmacological effect from the continuous pre-administration of synaptic blockers (5mM kynurenic acid + 1μM strychnine + 100μM picrotoxin) in bath. Repeated measures two-way ANOVA, ****p<0.0001, ns = not significant, please see **Supplementary Datasheet 1** for detailed statistics. All of data are represented as mean ± SD Abbreviations: Phe: phenylephrine, McN: McN-A-343, Mus: muscarine, Pir: pirenzepine, Calc: calcitonin, U50: U50488, SNC: SNC-162, N/OFQ: Nociceptin, SubP: SubstanceP, Aden: adenosine, Bac: baclofen, OXA: orexin A, 8OH: 8OH-DPAT, Ana: anandamide, SST: somatostatin, SB: synaptic blockers.

**Table 1.**

| **Group I (G<sub>q</sub> coupled GPCRs)** |  |  |
| --- | --- | --- |
| Agonist | Target | Dosage |
| DOI | 5HTR2a/c | 10μM |
| Phenylephrine | α1-AR | 20μM |
| DHPG | Type I mGluR | 50μM |
| McN-A-343 | mAChR1 | 10μM |
| Muscarine | mAChRs | 5μM |
| Calcitonin | CalcitoninR | 20nM |

| **Group II (Pain & Stress related)** |  |  |
| --- | --- | --- |
| Agonist | Target | Dosage |
| U50 | Kappa-OR | 10μM |
| DAMGO | Mu-OR | 200nM |
| SNC162 | Delta-OR | 20μM |
| N/O FQ | NOPR | 2.5nM |
| CRF | CRFR | 1μM |
| SubstanceP | NK1R | 200nM |

**Table 1**
| **Group III (Others)** |  |  |
| --- | --- | --- |
| Agonist | Target | Dosage |
| Adenosine | AdenosineRs | 100μM |
| Baclofen | GABABR | 20μM |
| OrexinA | OXR1/2 | 50nM |
| 8OH-DAPT | 5HTR1a | 10μM |
| Anandimide | CBR1/2 | 10μM |
| Somatostatin | SomatostatinR | 20nM |

**Table 2.**

| Group I Agonist | Target Receptor | Repeat measures two-way ANOVA followed by Bonferroni test; Across Pharmacology: F(5,595)=12.27, p<0.0001; Baseline-Drug: F(1,595)=762.4, p<0.0001; Across Pharmacology x Baseline-Drug: F(5,595)=67.40, p<0.0001; Subjects(ROI): F(595,595)=9.494, p<0.0001 |
| --- | --- | --- |
| DOI | 5HTR2a/c | Baseline vs. Drug :1.99 ± 0.89 vs. 2.08 ± 0.89 Hz, p=0.3359, n=105 ROIs |
| Phenylephrine | αAR | Baseline vs. Drug :1.93 ± 0.81 vs. 2.11 ± 0.80 Hz, p=0.0016, n=105 ROIs |
| DHPG | Group I mGluR | Baseline vs. Drug :1.92 ± 0.77 vs. 2.79 ± 0.80 Hz, p<0.0001, n=89 ROIs |
| McN-A-343 | mAChR1 | Baseline vs. Drug :1.91 ± 0.89 vs. 2.35 ± 0.87 Hz, p<0.0001, n=99 ROIs |
| Muscarine + Pirenzepine | mAChR3 | Baseline vs. Drug :2.05 ± 0.72 vs. 2.65 ± 0.79 Hz, p<0.0001, n=104 ROIs |
| Calcitonin | CalcitoninR | Baseline vs. Drug :2.16 ± 0.76 vs. 3.35 ± 0.86 Hz, p<0.0001, n=95 ROIs |

| Group II Agonist | Target Receptor | Repeat measures two-way ANOVA followed by Bonferroni test; Across Pharmacology: F(5,643)=12.89, p<0.0001; Baseline-Drug: F(1,651)=34.04, p<0.0001; Across Pharmacology x Baseline-Drug: F(5,643)=324.2, p<0.0001; Subjects(ROI): F(643,643)=18.97, p<0.0001 |
| --- | --- | --- |
| U50488 | KOR | Baseline vs. Drug: 1.37 ± 0.73 vs. 1.41 ± 0.73 Hz, p>0.9999, n=108 ROIs |
| DAMGO | MOR | Baseline vs. Drug: 1.42 ± 0.74 vs. 0.85 ± 0.70 Hz, p<0.0001, n=108 ROIs |
| SNC162 | DOR | Baseline vs. Drug: 1.33 ± 0.81 vs. 1.30 ± 0.82 Hz, p>0.9999, n=109 ROIs |
| N/O FQ | NOPR | Baseline vs. Drug: 1.52 ± 0.73 vs. 1.22 ± 0.76 Hz, p<0.0001, n=108 ROIs |
| CRF | CRFRs | Baseline vs. Drug: 1.25 ± 0.68 vs. 1.46 ± 0.75 Hz, p<0.0001, n=108 ROIs |
| Substance P | NK1R | Baseline vs. Drug: 1.33 ± 0.69 vs. 2.44 ± 0.78 Hz, p<0.0001, n=108 ROIs |

Table 2
| Group III Agonist | Target Receptor | Repeat measures two-way ANOVA followed by Bonferroni test; Across Pharmacology: F(5,352)=9.014, p<0.0001; Baseline-Drug: F(1,352)=23.70, p<0.0001; Across Pharmacology x Baseline-Drug: F(5,352)=156.8, p<0.0001; Subjects(ROI): F(352,352)=23.37, p<0.0001 |
| --- | --- | --- |
| Adenosine | AdenosineRs | Baseline vs. Drug: 1.76 ± 0.68 vs. 1.82 ± 0.75 Hz, p>0.5382, n=52 ROIs |
| Baclofen | GABABR | Baseline vs. Drug: 1.65 ± 0.79 vs. 1.24 ± 0.74 Hz, p<0.0001, n=63 ROIs |
| OrexinA | OrexinRs | Baseline vs. Drug: 1.76 ± 0.91 vs. 1.25 ± 0.85 Hz, p<0.0001, n=61 ROIs |
| 8OH-DPAT | 5HTR1a | Baseline vs. Drug: 1.83 ± 0.77 vs. 1.50 ± 0.67 Hz, p<0.0001, n=60 ROIs |
| Anandamide | CBRs | Baseline vs. Drug: 1.31 ± 0.69 vs. 1.54 ± 0.74 Hz, p<0.0001, n=61 ROIs |
| Somatostatin | SomatostatinR | Baseline vs. Drug: 1.32 ± 0.66 vs. 0.76 ± 0.51 Hz, p<0.0001, n=61 ROIs |

### Modular-selective pharmacological profile of LC-NE neurons

The non-homogenous response of LC-NE neurons to agonists activating various GPCRs suggests that these receptors may underly endogenous control of LC modules. It could be that GPCR-mediated regulation across anatomically defined LC efferent modules enables rapid reallocation of noradrenergic resources across brain circuits during distinct behaviors^20,21,27–29,42,43,56^. To investigate the pharmacological profile of anatomically defined LC modules, we next combined our *ex vivo* calcium imaging approach with fluorescent retrograde tracing to identify LC neurons by their efferent targets. To do so, we injected the common retrograde neuronal tracer cholera toxin subunit b conjugated to a CF 594 fluorescent tag (Ctb594) into the mPFC (**Fig. 3A&B**). The mPFC is a well-established downstream brain region that forms robust reciprocal axonal projections with LC. In the dorsal pons, almost all Ctb594 was colocalized with tyrosine hydroxylase (TH; the rate-limiting enzyme for catecholamine synthesis) immunoreactive neurons. These LC-mPFC neurons were scattered across the whole LC except for sparse distribution in the rostroventral LC (**Fig. 3C, S8**), where many spinally-projecting LC-NE neurons are located^28,47^. We next repeated the *ex vivo* multiplexed pharmacological scan and to determine the pharmacological response of LC-mPFC neurons to the eighteen GPCR agonists. To ensure accuracy of the Ctb594 signal, we reconstructed recorded slices using confocal z-stack scanning in concert with post-hoc immunohistochemistry of GCaMP8f and Ctb594. The extracted ROIs were then registered to their modular category (LC-mPFC or non-LC-mPFC) based on the presence Ctb594 (**Fig. 3C-F**). Surprisingly, some agonists drove differential effects between mPFC and non-mPFC-projecting LC-NE modules. In particular, we observed stronger inhibition of LC-mPFC neurons by DAMGO and the 5HT1a receptor agonist 8OH-DPAT. For the non-LC-mPFC-module, McN-A-343, a selective mAChR1 agonist, preferentially drove excitation, while baclofen, a GABAB receptor (GABABR) agonist, preferentially inhibited the non-mPFC projecting LC module (**Fig. 3G-J S9, S10**). The detailed statistical information for the modular multiplexed pharmacological scan is listed in **Table 3 and Table 4**. To eliminate the possibility that these effects could arise from differential spontaneous firing at baseline, we compared the baseline activity between modules and no significant difference was found across applications in any group of agonists (**Fig. S11**). Together these findings demonstrate pharmacological dissection of modular efferent function in the LC.

**Figure 3.**
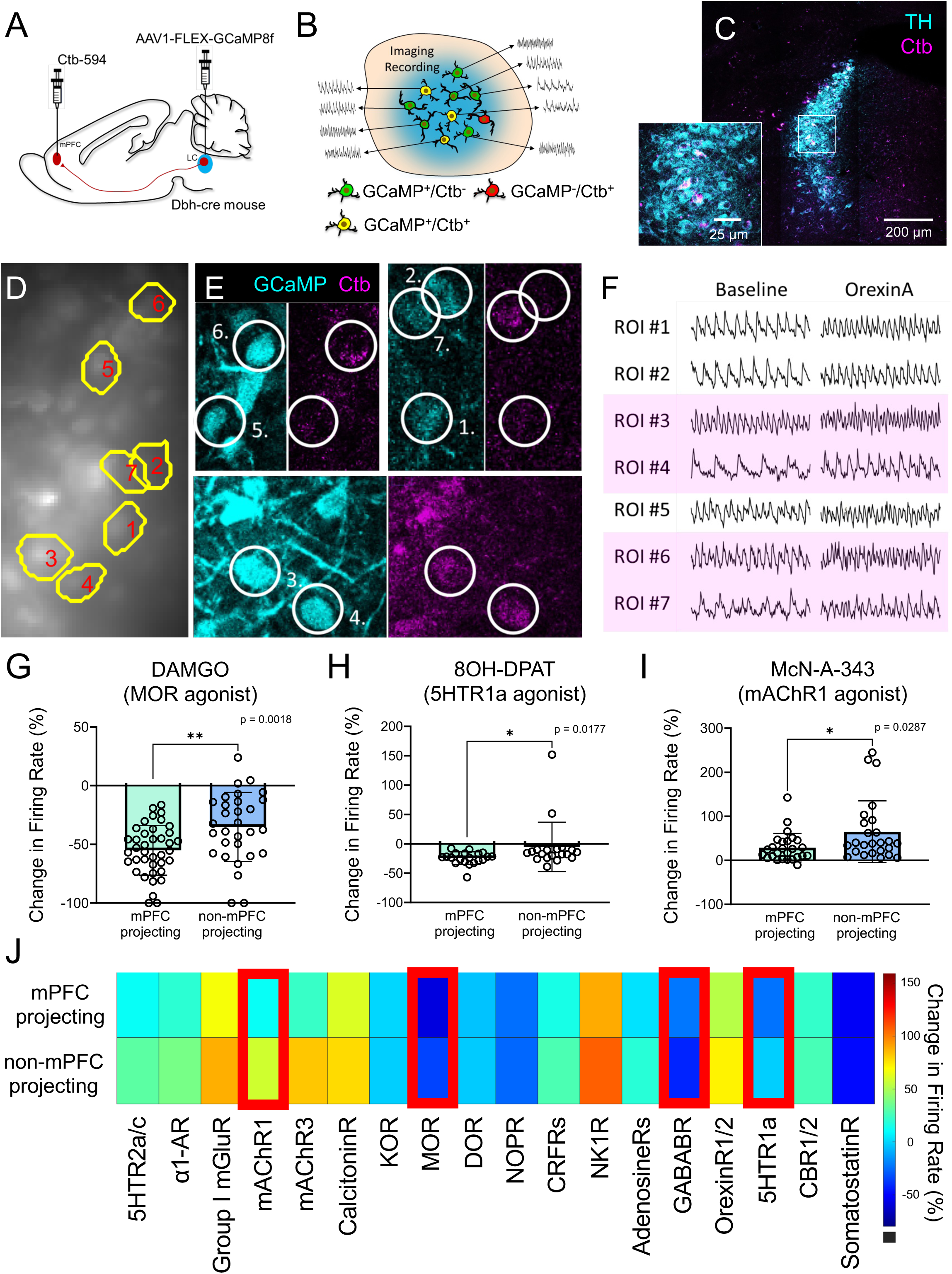
Pharmacological scan of GPCRs targeting the LC-mPFC module. (A) A schematic cartoon describing the viral and modular tracing strategies from mPFC to LC. (B) A schematic cartoon demonstrating the recording setup. (C) A representative fluorescent image showing the distribution of mPFC-projecting LC-NE neurons indicated by the colocalization of TH (cyan) and Ctb (magenta) immunoreactive signals. (D-F) Representative examples from 7 extracted ROIs showing their spatial information in D, and modular identity indicated by post-hoc morphological reconstruction in E, F shows pharmacological responses of orexin A that activates orexin receptors. Note the consistent numerical naming of ROIs across figures. GCaMP8f and Ctb immunoreactive signals are shown by cyan and magenta color in E, and pink backgrounds in F denote the mPFC-projecting modular identity. (G-I) Plots showing the firing rate results of DAMGO, 8OH-DPAT and McN-A-343 across mPFC- and non-mPFC-projecting LC-NE neurons. DAMGO: Student’s t-test, t = 3.260, **p<0.01. 8OH-DPAT: Mann-Whitney test, U = 106, *p<0.05. McN-A-343: Mann-Whitney test, U = 209, *p<0.05. (J) A color plot demonstrating the visualized results of modular pharmacological scan, pharmacological effects are displayed by color and the targeted GPCRs by each agonist are listed. The red rectangles indicate significant difference between pharmacological effects in different LC modules. Please see **Table 3** and **Table 4** for statistical details. All of data are represented as mean ± SD. Abbreviations: 5HTR2a/c: serotonin receptor 2a/c, α1-AR: alpha1-adrenergic receptor, mAChR1/3: muscarinic receptor 1/3, CalcitoninR: calcitonin receptor, KOR: kappa opioid receptor, DOR: delta opioid receptor, MOR: mu opioid receptor, NOPR: nociceptin receptor, CRFR: corticotropin-releasing factor receptor, NK1R: neurokinin 1 receptor, AdenosineRs: adenosine receptors, GABABR: GABAB receptor, Orexin1/2R: orexin receptors 1/2, 5HTR1a: serotonin receptor 1a, CB1/2R: cannabinoid receptor 1/2, SomatostatinR: somatostatin receptor.

**Table 3.**

| Group I | Firing Rate (mPFC-projecting vs. non mPFC-projecting) |
| --- | --- |
| DOI<br>(5HTR2a/c) | Repeat measures two-way ANOVA followed by Bonferroni test; Pharmacology : F(1,57)=36.35, p<0.0001; Modular : F(1,57)=5.464, p=0.0229; Pharmacology x Modular : F(1,57)=0.7062, p=0.4042; Subjects(ROI) : F(57,57)=38.23, p<0.0001; n=28/27 ROIs; mPFC-projecting : 2.23 ± 0.85 vs. 2.41 ± 0.90 Hz, p=0.001; non mPFC-projecting : 1.67 ± 0.74 vs. 1.92 ± 0.75 Hz, p<0.0001 |
| Phenylephrine<br>(αAR) | Repeat measures two-way ANOVA followed by Bonferroni test; Pharmacology : F(1,57)=30.16, p<0.0001; Modular : F(1,57)=3.396, p=0.0705; Pharmacology x Modular : F(1,57)=0.8966, p=0.3477; Subjects(ROI) : F(57,57)=45.89, p<0.0001; n=28/27 ROIs; mPFC-projecting : 2.16 ± 0.80 vs. 2.38 ± 0.82 Hz, p<0.0001; non mPFC-projecting : 1.79 ± 0.91 vs. 1.94 ± 0.88 Hz, p=0.0047 |
| DHPG<br>(Group I mGluR) | Repeat measures two-way ANOVA followed by Bonferroni test; Pharmacology : F(1,54)=133.7, p<0.0001; Modular : F(1,54)=0.00, p>0.9999; Pharmacology x Modular : F(1,54)=0.00, p>0.9999; Subjects(ROI) : F(54,54)=5.850, p<0.0001; n=26/20 ROIs; mPFC-projecting : 2.12 ± 0.89 vs. 3.21 ± 0.68 Hz, p<0.0001; non mPFC-projecting : 1.58 ± 0.95 vs. 2.45 ± 0.90 Hz, p<0.0001 |
| McN-A-343<br>(mAChR1) | Repeat measures two-way ANOVA followed by Bonferroni test; Pharmacology : F(1,53)=111.6, p<0.0001; Modular : F(1,53)=3.749, p=0.0582; Pharmacology x Modular : F(1,53)=4.862, p=0.318; Subjects(ROI) : F(53,53)=18.33, p<0.0001; n=26/25 ROIs; mPFC-projecting : 2.14 ± 0.88 vs. 2.57 ± 0.82 Hz, p<0.0001; non mPFC-projecting : 1.59 ± 0.82 vs. 2.26 ± 0.74 Hz, p<0.0001 |
| Muscarine +<br>Pirenzepine<br>(mAChR) | Repeat measures two-way ANOVA followed by Bonferroni test; Pharmacology : F(1,52)=60.16, p<0.0001; Modular : F(1,52)=3.702, p=0.0598; Pharmacology x Modular : F(1,52)=0.1291, p=0.7208; Subjects(ROI) : F(52,52)=5.217, p<0.0001; n=27/27 ROIs; mPFC-projecting : 2.25 ± 0.82 vs. 2.90 ± 0.72 Hz, p<0.0001; non mPFC-projecting : 1.82 ± 0.80 vs. 2.48 ± 0.87 Hz, p<0.0001 |
| Calcitonin<br>(CalcitoninR) | Repeat measures two-way ANOVA followed by Bonferroni test; Pharmacology : F(1,49)=139.9, p<0.0001; Modular : F(1,49)=2.581, p=0.1146; Pharmacology x Modular : F(1,49)=0.0930, p=0.7617; Subjects(ROI) : F(49,49)=3.970, p<0.0001; n=22/25 ROIs; mPFC-projecting : 2.36 ± 0.80 vs. 3.70 ± 0.39 Hz, p<0.0001; non mPFC-projecting : 1.91 ± 0.77 vs. 3.30 ± 1.08 Hz, p<0.0001 |

| Group II | Firing Rate (mPFC-projecting vs. non mPFC-projecting) |
| --- | --- |
| U50488<br>(KOR) | Repeat measures two-way ANOVA followed by Bonferroni test; Pharmacology : F(1,64)=0.1093, p=0.7420; Modular : F(1,64)=0.3974, p=0.5307; Pharmacology x Modular : F(1,64)=2.587, p=0.1126; Subjects(ROI) : F(64,64)=25.86, p<0.0001; n=38/28 ROIs; mPFC-projecting : 1.41 ± 0.78 vs. 1.48 ± 0.81 Hz, p=0.2830; non mPFC-projecting : 1.35 ± 0.84 vs. 1.30 ± 0.70 Hz, p=0.8057 |
| DAMGO<br>(MOR) | Repeat measures two-way ANOVA followed by Bonferroni test; Pharmacology : F(1,67)=129.9, p<0.0001; Modular : F(1,67)=2.138, p=0.1483; Pharmacology x Modular : F(1,67)=12.93, p=0.0006; Subjects(ROI) : F(67,67)=15.69, p<0.0001; n=39/30 ROIs; mPFC-projecting : 1.37 ± 0.71 vs. 0.68 ± 0.53 Hz, p<0.0001; non mPFC-projecting : 1.48 ± 0.96 vs. 1.12 ± 0.92 Hz, p<0.0001 |
| SNC 162<br>(DOR) | Repeat measures two-way ANOVA followed by Bonferroni test; Pharmacology : F(1,70)=1.953, p=0.1666; Modular : F(1,70)=0.3459, p=0.5583; Pharmacology x Modular : F(1,70)=0.0938, p=0.7604; Subjects(ROI) : F(70,70)=82.44, p<0.0001; n=38/30 ROIs; mPFC-projecting : 1.41 ± 0.87 vs. 1.38 ± 0.87 Hz, p=0.8316; non mPFC-projecting : 1.30 ± 0.83 vs. 1.26 ± 0.88 Hz, p=0.5139 |
| N/O FQ<br>(NOPR) | Repeat measures two-way ANOVA followed by Bonferroni test; Pharmacology : F(1,69)=87.53, p<0.0001; Modular : F(1,69)=0.6730, p=0.4148; Pharmacology x Modular : F(1,69)=0.1874, p=0.6664; Subjects(ROI) : F(69,69)=33.01, p<0.0001; n=37/30 ROIs; mPFC-projecting : 1.52 ± 0.69 vs. 1.22 ± 0.73 Hz, p<0.0001; non mPFC-projecting : 1.69 ± 0.84 vs. 1.36 ± 0.93 Hz, p<0.0001 |
| CRF<br>(CRFRs) | Repeat measures two-way ANOVA followed by Bonferroni test; Pharmacology : F(1,71)=59.72, p<0.0001; Modular : F(1,71)=0.3489, p=0.5566; Pharmacology x Modular : F(1,71)=1.019, p=0.3163; Subjects(ROI) : F(71,71)=43.97, p<0.0001; n=39/30 ROIs; mPFC-projecting : 1.25 ± 0.65 vs. 1.44 ± 0.70 Hz, p<0.0001; non mPFC-projecting : 1.33 ± 0.87 vs. 1.57 ± 0.97 Hz, p<0.0001 |
| Substance P<br>(NK1R) | Repeat measures two-way ANOVA followed by Bonferroni test; Pharmacology : F(1,70)=308.8, p<0.0001; Modular : F(1,70)=0.3076, p=0.5810; Pharmacology x Modular : F(1,70)=0.0981, p=0.7551; Subjects(ROI) : F(70,70)=8.334, p<0.0001; n=39/30 ROIs; mPFC-projecting : 1.32 ± 0.65 vs. 2.41 ± 0.74 Hz, p<0.0001; non mPFC-projecting : 1.40 ± 0.90 vs. 2.54 ± 0.88 Hz, p<0.0001 |

**Table 3**
| Group III | Firing Rate (mPFC-projecting vs. non mPFC-projecting) |
| --- | --- |
| Adenosine<br>(AdenosineR) | Repeat measures two-way ANOVA followed by Bonferroni test; Pharmacology : F(1,39)=1.449, p=0.2360; Modular : F(1,39)=1.739, p=0.1949; Pharmacology x Modular : F(1,39)=0.1638, p=0.2082; Subjects(ROI) : F(39,39)=23.22, p<0.0001; n=18/19 ROIs; mPFC-projecting : 1.90 ± 0.56 vs. 2.02 ± 0.61 Hz, p=0.1813; non mPFC-projecting : 1.63 ± 0.87 vs. 1.62 ± 0.94 Hz, p>0.9999 |
| Baclofen<br>(GABABR) | Repeat measures two-way ANOVA followed by Bonferroni test; Pharmacology : F(1,43)=86.07, p<0.0001; Modular : F(1,43)=1.166, p=0.2862; Pharmacology x Modular : F(1,43)=5.174, p=0.0280; Subjects(ROI) : F(43,43)=23.19, p<0.0001; n=21/20 ROIs; mPFC-projecting : 1.73 ± 0.83 vs. 1.40 ± 0.71 Hz, p<0.0001; non mPFC-projecting : 1.60 ± 0.85 vs. 1.03 ± 0.82 Hz, p<0.0001 |
| OrexinA<br>(OrexinRs) | Repeat measures two-way ANOVA followed by Bonferroni test; Pharmacology : F(1,42)=179.2, p<0.0001; Modular : F(1,42)=0.5095, p=0.4793; Pharmacology x Modular : F(1,42)=0.8579, p=0.3596; Subjects(ROI) : F(42,42)=20.24, p<0.0001; n=21/19 ROIs; mPFC-projecting : 1.39 ± 0.53 vs. 2.08 ± 0.63 Hz, p<0.0001; non mPFC-projecting : 1.28 ± 0.74 vs. 1.88 ± 0.90 Hz, p=0.002 |
| 8OH-DAPT<br>(5HTR1a) | Repeat measures two-way ANOVA followed by Bonferroni test; Pharmacology : F(1,41)=69.41, p<0.0001; Modular : F(1,41)=0.5580, p=0.4593; Pharmacology x Modular : F(1,41)=10.63, p=0.0022; Subjects(ROI) : F(41,41)=34.01, p<0.0001; n=20/19 ROIs; mPFC-projecting : 2.11 ± 0.64 vs. 1.62 ± 0.58 Hz, p<0.0001; non mPFC-projecting : 1.78 ± 0.99 vs. 1.57 ± 0.87 Hz, p<0.0001 |
| Anadamide<br>(CBRs) | Repeat measures two-way ANOVA followed by Bonferroni test; Pharmacology : F(1,42)=35.05, p<0.0001; Modular : F(1,42)=0.4614, p=0.5007; Pharmacology x Modular : F(1,42)=0.7580, p=0.3889; Subjects(ROI) : F(42,42)=30.24, p<0.0001; n=21/19 ROIs; mPFC-projecting : 1.40 ± 0.64 vs. 1.60 ± 0.64 Hz, p=0.0014; non mPFC-projecting : 1.21 ± 0.76 vs. 1.48 ± 0.90 Hz, p<0.0001 |
| Somatostatin<br>(SomatostatinR) | Repeat measures two-way ANOVA followed by Bonferroni test; Pharmacology : F(1,40)=160.4, p<0.0001; Modular : F(1,40)=0.9097, p=0.3459; Pharmacology x Modular : F(1,40)=0.0271, p=0.8700; Subjects(ROI) : F(40,40)=16.14, p<0.0001; n=20/18 ROIs; mPFC-projecting : 1.45 ± 0.52 vs. 0.89 ± 0.43 Hz, p<0.0001; non mPFC-projecting : 1.28 ± 0.82 vs. 0.70 ± 0.58 Hz, p<0.0001 |

**Table 4.**
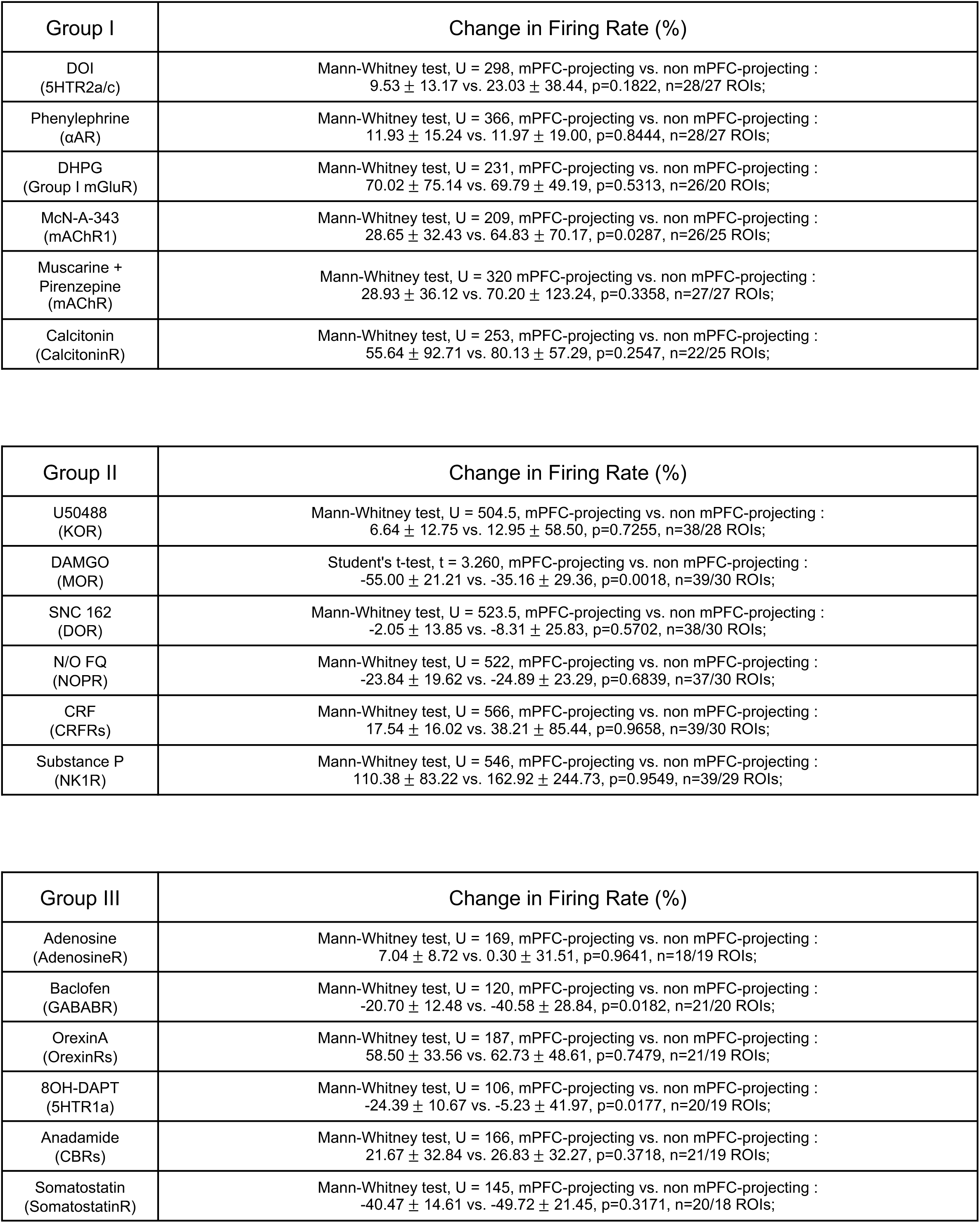

| Group I | Change in Firing Rate (%) |
| --- | --- |
| DOI<br>(5HTR2a/c) | Mann-Whitney test, U = 298, mPFC-projecting vs. non mPFC-projecting :<br>9.53 ± 13.17 vs. 23.03 ± 38.44, p=0.1822, n=28/27 ROIs; |
| Phenylephrine<br>(αAR) | Mann-Whitney test, U = 366, mPFC-projecting vs. non mPFC-projecting :<br>11.93 ± 15.24 vs. 11.97 ± 19.00, p=0.8444, n=28/27 ROIs; |
| DHPG<br>(Group I mGluR) | Mann-Whitney test, U = 231, mPFC-projecting vs. non mPFC-projecting :<br>70.02 ± 75.14 vs. 69.79 ± 49.19, p=0.5313, n=26/20 ROIs; |
| McN-A-343<br>(mAChR1) | Mann-Whitney test, U = 209, mPFC-projecting vs. non mPFC-projecting :<br>28.65 ± 32.43 vs. 64.83 ± 70.17, p=0.0287, n=26/25 ROIs; |
| Muscarine +<br>Pirenzepine<br>(mAChR) | Mann-Whitney test, U = 320 mPFC-projecting vs. non mPFC-projecting :<br>28.93 ± 36.12 vs. 70.20 ± 123.24, p=0.3358, n=27/27 ROIs; |
| Calcitonin<br>(CalcitoninR) | Mann-Whitney test, U = 253, mPFC-projecting vs. non mPFC-projecting :<br>55.64 ± 92.71 vs. 80.13 ± 57.29, p=0.2547, n=22/25 ROIs; |

| Group II | Change in Firing Rate (%) |
| --- | --- |
| U50488<br>(KOR) | Mann-Whitney test, U = 504.5, mPFC-projecting vs. non mPFC-projecting :<br>6.64 ± 12.75 vs. 12.95 ± 58.50, p=0.7255, n=38/28 ROIs; |
| DAMGO<br>(MOR) | Student's t-test, t = 3.260, mPFC-projecting vs. non mPFC-projecting :<br>-55.00 ± 21.21 vs. -35.16 ± 29.36, p=0.0018, n=39/30 ROIs; |
| SNC 162<br>(DOR) | Mann-Whitney test, U = 523.5, mPFC-projecting vs. non mPFC-projecting :<br>-2.05 ± 13.85 vs. -8.31 ± 25.83, p=0.5702, n=38/30 ROIs; |
| N/O FQ<br>(NOPR) | Mann-Whitney test, U = 522, mPFC-projecting vs. non mPFC-projecting :<br>-23.84 ± 19.62 vs. -24.89 ± 23.29, p=0.6839, n=37/30 ROIs; |
| CRF<br>(CRFRs) | Mann-Whitney test, U = 566, mPFC-projecting vs. non mPFC-projecting :<br>17.54 ± 16.02 vs. 38.21 ± 85.44, p=0.9658, n=39/30 ROIs; |
| Substance P<br>(NK1R) | Mann-Whitney test, U = 546, mPFC-projecting vs. non mPFC-projecting :<br>110.38 ± 83.22 vs. 162.92 ± 244.73, p=0.9549, n=39/29 ROIs; |

**Table 4**
| Group III | Change in Firing Rate (%) |
| --- | --- |
| Adenosine<br>(AdenosineR) | Mann-Whitney test, U = 169, mPFC-projecting vs. non mPFC-projecting :<br>7.04 ± 8.72 vs. 0.30 ± 31.51, p=0.9641, n=18/19 ROIs; |
| Baclofen<br>(GABABR) | Mann-Whitney test, U = 120, mPFC-projecting vs. non mPFC-projecting :<br>-20.70 ± 12.48 vs. -40.58 ± 28.84, p=0.0182, n=21/20 ROIs; |
| OrexinA<br>(OrexinRs) | Mann-Whitney test, U = 187, mPFC-projecting vs. non mPFC-projecting :<br>58.50 ± 33.56 vs. 62.73 ± 48.61, p=0.7479, n=21/19 ROIs; |
| 8OH-DAPT<br>(5HTR1a) | Mann-Whitney test, U = 106, mPFC-projecting vs. non mPFC-projecting :<br>-24.39 ± 10.67 vs. -5.23 ± 41.97, p=0.0177, n=20/19 ROIs; |
| Anadamide<br>(CBRs) | Mann-Whitney test, U = 166, mPFC-projecting vs. non mPFC-projecting :<br>21.67 ± 32.84 vs. 26.83 ± 32.27, p=0.3718, n=21/19 ROIs; |
| Somatostatin<br>(SomatostatinR) | Mann-Whitney test, U = 145, mPFC-projecting vs. non mPFC-projecting :<br>-40.47 ± 14.61 vs. -49.72 ± 21.45, p=0.3171, n=20/18 ROIs; |

### Synergistic antinociceptive effects from modular-selective pharmacology

Important prior studies have shown clear modularity in LC-mediated pain modulation. Activation of the LC-mPFC projection is thought to drive a pronociceptive tone while noradrenergic projections to the spinal cord exert descending antinociception^28,44^. Furthermore, LC efferent targeting other brain regions has been found to engage in different aspects of aversive and pain-related behaviors^27,28,30,42,43,45,46,49,51,52,54,56,73^. Accordingly, to drive antinociception, an ideal strategy for LC modulation is to selectively dampen the pronociceptive activity of LC-mPFC projection, leading a reallocation of noradrenergic modulation across the central nervous system. We next sought to test whether the agonists we identified to shift neural activity away from the LC-mPFC module *ex vivo* (**Fig. 3G-J**) could lead to *in vivo* antinociception. Here we selected DAMGO, 8OH-DPAT and McN-A-343 for *in vivo* testing. To do so, we implanted bilateral cannula above the LC in C57BL/6J mice. We began with single compound infusions and tested thermal paw withdrawal latencies using the Hargreaves test (**Fig. 4A&B**). We calculated the ratio between measurements under baseline and pharmacological conditions (**Fig. 4A&B, S12, S13**). We constructed dose-response curves from single compound infusions and found that DAMGO and 8OH-DPAT produced significant antinociception (**Fig. 4C&D**). McN-A-343 elicited a smaller, subtly antinociceptive effect when infused to LC (**Fig. 4E**). Following these dose-response curves, we then designed two pharmacological cocktails of these agonists based on their *in vivo* EC_50_ values (DAMGO: 0.46μM,8OH-DPAT: 1.96μM, McN-A-343: 2.27μM), namely, Cocktail A (DAMGO + 8OH-DPAT = 1:4) and Cocktail B (DAMGO + 8OH-DPAT + McN-A-343 = 1:4:5). Interestingly, when we generated isobolograms from the dose-response curves of these two pharmacological cocktails, there was significant synergistic antinociception driven by Cocktail A, but not Cocktail B (EC_50_: 0.46μM for Cocktail A, 2.1μM for Cocktail B) (**Fig. 4F-J, S14**). Surprisingly, Cocktail B caused more potent antinociception than any other ligand or cocktail (**Fig. 4G-H, J, S12**). However, three-dimensional isobolographic analysis^74^ shows an additive agonist interaction where Cocktail B only enhances antinociception past the other drug conditions once Cocktail B reaches a minimum dose threshold **(Fig. S14)**. Because these tests were performed within-subject over many weeks, we considered the possibility that baseline nociception could drift across the course of pharmacological treatments and behavioral examination, however no significant differences were found across baseline measurements in seven trials (**Fig. S13**). Furthermore, the thermal plantar assay relies on intact locomotor responses such that any sedative effect could lead to the fictive appearance of antinociception. Cocktail B, which provides the greatest paw withdrawal latency, did not alter locomotion in the open field test (**Fig. S15**), suggesting that the increased paw withdrawal latency is indeed driven by reduced detection of the noxious thermal stimulus. We then returned to *ex vivo* slice imaging to determine whether the differential response across LC modules during a co-application of these three agonists indeed drives activity away from the LC-mPFC. As hypothesized, the application of Cocktail B preferentially inhibited the LC-mPFC module (**Fig. 4K, S16**). Taken together, our *ex vivo* calcium imaging approach appears to have identified a novel pharmacological strategy for LC-mediated antinociception.

**Figure 4.**
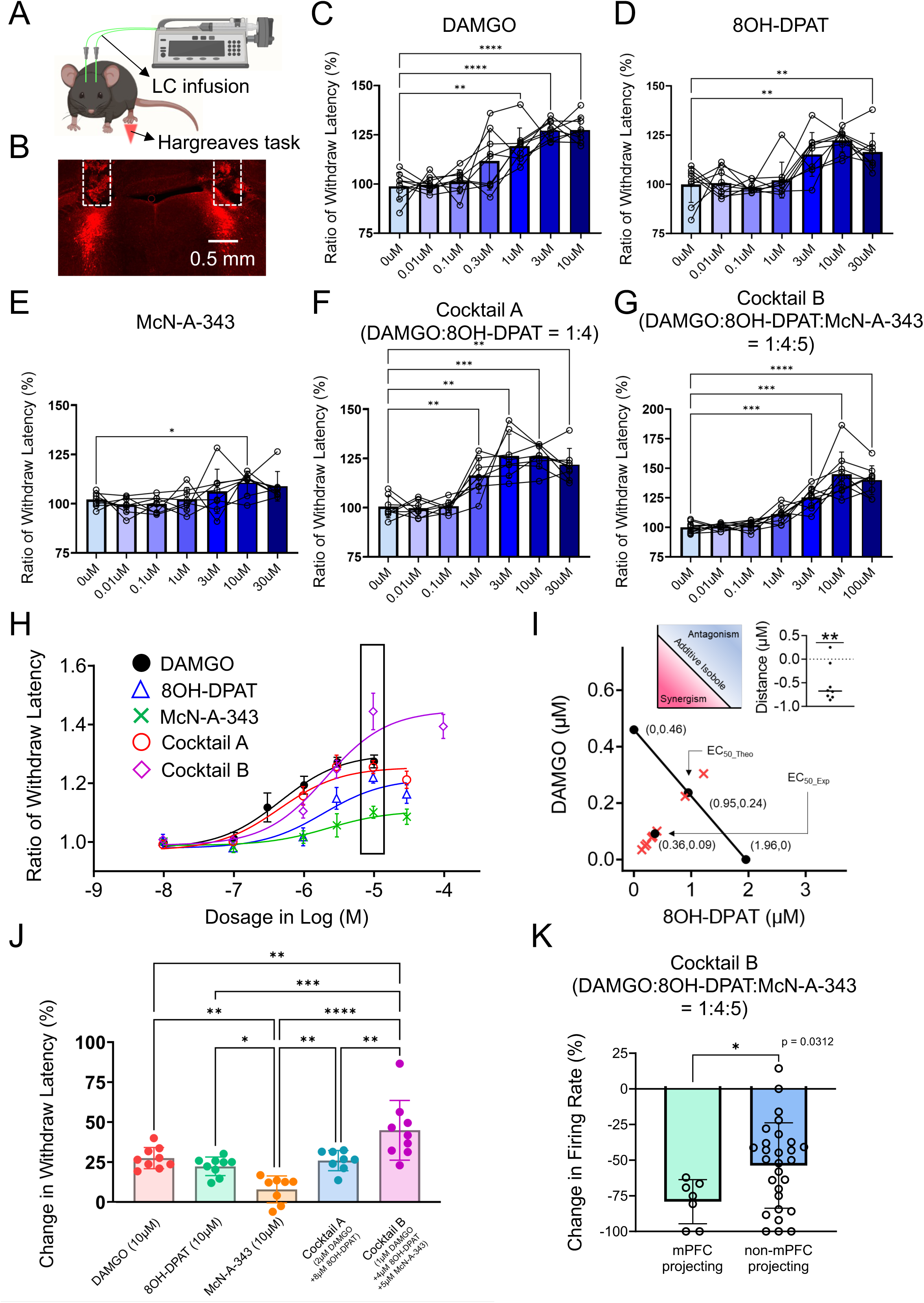
Multiple pharmacological approaches drive antinociception by shifting activity away from the LC-mPFC module. (A) A schematic cartoon illustrating the experimental design of Hargreaves thermal plantar assay in concert with LC local pharmacological infusions. (B) A fluorescent image showing a representative cannula implantation toward bilateral LC, which is denoted by TH immunoreactive signals (Red). The dashed rectangles indicate the trajectory of cannula. (C-G) Plots demonstrating the analgesia led by pharmacological infusions of DAMGO, 8OH-DPAT, McN-A-343, Cocktail A and Cocktail B toward LC with different dosages. The data are calculated from ratios between paw withdrawal latencies under baseline and pharmacological conditions. DAMGO: Repeated measures one-way ANOVA followed by Dunnett’s test, F = 20.77, **p<0.01, ****p<0.0001. 8OH-DPAT: Repeated measures one-way ANOVA followed by Dunnett’s test, F = 11.90, **p<0.01. McN-A-343: Friedman test followed by Dunn’s test, Friedman statistic = 22.82, *p<0.05. Cocktail A: Repeated measures one-way ANOVA followed by Dunnett’s test, F = 22.07, **p<0.01, ***p<0.001. Cocktail B: Repeated measures one-way ANOVA followed by Dunnett’s test, F = 33.86, ***p<0.001, ****p<0.0001. (H) A plot showing pharmacological dose-response curves from data shown in C. The rectangle denotes the pharmacological response with 10μM in dosage. (I) An isobobogram showing the synergistic interaction within Cocktail A, the experimental and theoretical EC_50_ of Cocktail A as well as the EC_50_ of DAMGO and 8OH-DPAT are indicated by black dots, the experimental EC_50_s of individual mice are indicated by red cross and the black line denotes the additive isobole. The upper right plot shows a statistic evaluation of synergism. One-sample Wilcoxon Signed Rank Test, *p<0.05. (J) A plot showing the comparison of antinociceptive effects upon infusions of different pharmacological approaches in fixed dosage shown in D (10μM). Note the greater analgesia led by Cocktail B. One-way ANOVA followed by Holm-Šídák’s test, F = 13.52, *p<0.05, **p<0.01, ***p<0.001, ****p<0.0001. (K) A plot demonstrating the percentile of changes in firing rate by administration of Cocktail B using *ex vivo* pharmacological scan. Student’s t-test, t = 2.150, *p<0.05. All data are represented as mean ± SD except SEMs used in the isologram in I.

### Modular deletion of LC-MOR disrupts cocktail-induced antinociception

One possibility for how LC modules drive discreet behavioral functions is through interaction, either competitive or cooperative, between two or more efferent modules. To test whether such modular interaction is required for the intra-LC cocktail-mediated antinociception, we created an LC-mPFC module-selective conditional knockout of *oprm1*^fl/fl^ (mKO). To do so, we bilaterally injected a noradrenergic selective, retrogradely trafficked canine adenovirus to drive Cre recombinase expression under the synthetic PRSx8 promoter (CAV-PRS-Cre-V5) into the mPFC of C57BL/6J and *oprm1*^fl/fl^ homozygote mice^47,75,76^ (**Fig. 5A&B**). O*prm1*^fl/fl^ mice have loxP sites flanking the 2^nd^-3^rd^ exons of *oprm1* gene^77^. Consequently, selective expression of Cre recombinase drives a genetic deletion of MOR in LC-mPFC neurons. To visualize successful Cre recombination, in a subset of mice, we also introduced Cre-dependent expression of the red fluorophore, mCherry, into the LC by local delivery of AAV8-DIO-mCherry (**Fig. 5C**). The expression of mCherry in LC-mPFC neurons showed a consistent pattern to the retrograde tracing by Ctb594 (**Fig. 5C, S8, S17**). Having used this approach previously^42^, we corroborated functional modular MOR exclusion using DAMGO-induced inhibition in cell-attached recordings (**Fig. 5D&E**). We next injected another cohort of *oprm1*^fl/fl^ mice with either CAV-PRS-Cre-V5 or CAV-mCherry in the mPFC in concert with bilateral cannula implantations above the LC (**Fig. 5F**). Following deletion of LC-mPFC MOR, we once again tested the antinociceptive effect of different pharmacological infusions using the thermal paw withdrawal assay using a fixed concentration of ligands or cocktails (10μM, **Fig. 5G**). Consistent with our electrophysiological findings, we found LC-mPFC mKO mice lost LC MOR-mediated antinociception. In addition to DAMGO infusion, the LC-mPFC mKO surprisingly disrupted the antinociceptive action of Cocktail B (**Fig. 5H, S18, Table 5**). This suppression of Cocktail B-mediated antinociception could be caused by MOR on other LC-NE neurons in the non-LC-mPFC-projecting module. To test this hypothesis, we designed Cocktail C as a pharmacological combination of 8OH-DPAT and McN-A-343 in a fixed ratio (4:5). The infusion of Cocktail C elicited robust antinociception that was not affected by LC-mPFC MOR mKO (**Fig. 5H, S18**). This finding suggests that the activation of MOR on LC-NE neurons not projecting to mPFC could yield a pronociceptive response that competes against the antinociception driven by the MOR-insensitive portion of Cocktail B^26–28,47,56^. Further, these results show that both LC-mPFC and non-LC-mPFC neurons are actively involved in the pharmacologically-mediated antinociception, providing a net outcome from the relative stronger inhibition of the LC-mPFC innervating module. This differential pharmacologically mediated modulation was outflanked by the modular impairment of MOR-mediated regulation. This inter-module interaction in nociception is likely the consequence of integration across brain and spinal regions that receive noradrenergic inputs from distinct LC efferent modules. A recent study also demonstrated alpha2-adrenergic receptor-mediated lateral inhibition underlies some of the modular organization of LC^65^. This cross-modular inhibition is theoretically scaled by the concurrent excitation of LC modules, however the *in vivo* dynamics of cross-modular integration in response to nociceptive stimuli remains elusive. Altogether, here we demonstrate a novel approach to identify neural circuit-selective pharmacological strategies that identify a means of driving GPCR-mediated antinociception across LC modules. As hypothesized by us and others^28,42,43^, this neural circuit-selective antinociception requires an active interaction between different efferent LC modules.

**Figure 5.**
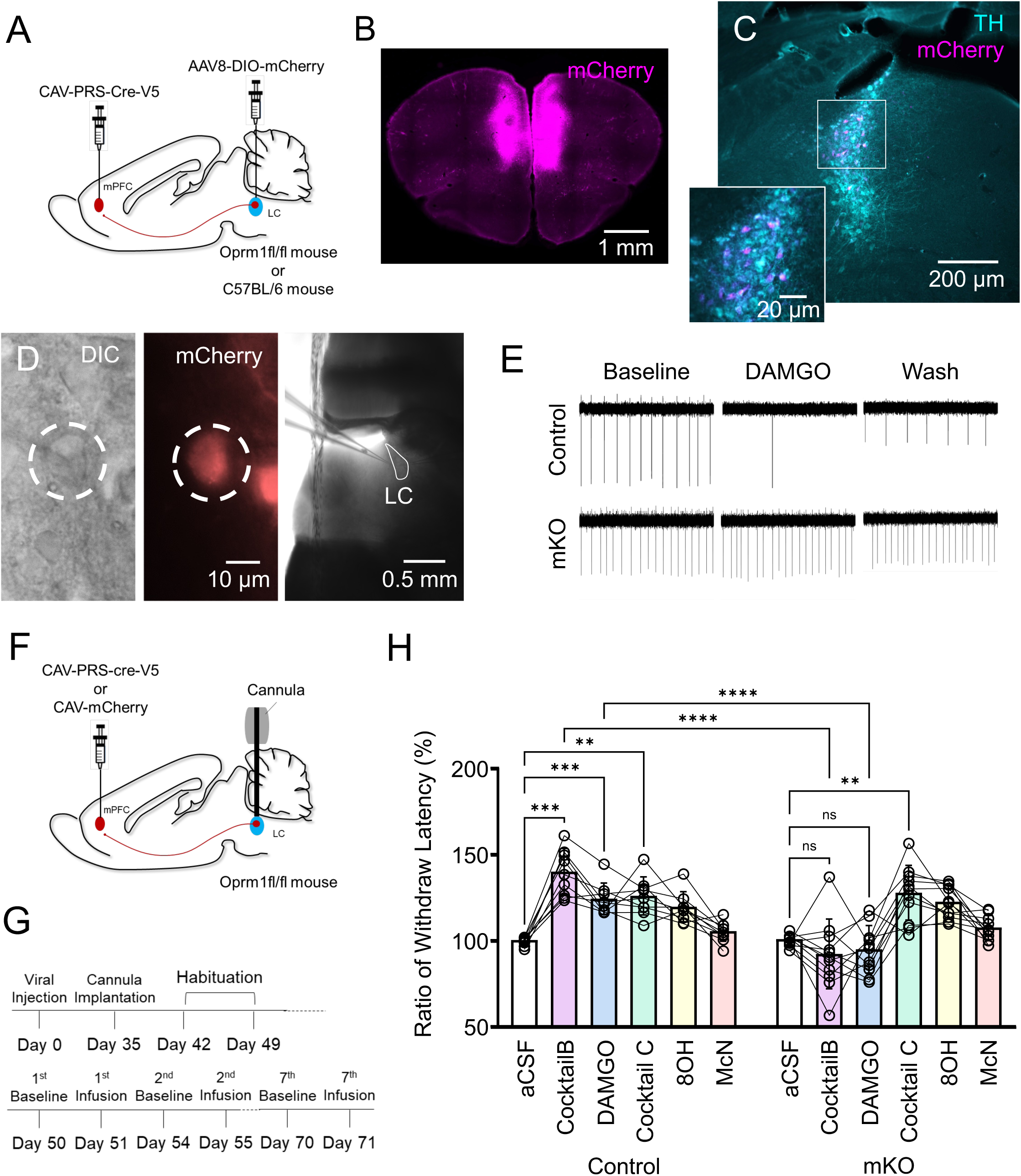
Modular, conditional LC-mPFC MOR knockout disrupts Cocktail B-mediated antinociception. (A) A schematic cartoon illustrating the viral strategies for the modular deletion of MOR with fluorescent labeling in LC. (B) A fluorescent image showing the injection site of CAV-mCherry into mPFC. (C) A representative fluorescent image demonstrating the distribution of mPFC innervating LC-NE neurons by the colocalization of TH (cyan) and mCherry (magenta) immunoreactive signals. (D) An example of live DIC and fluorescent images during a cell-attached recording (left & middle), the right panel shows a DIC image under 5x objective lens. (E) A representative cell-attached recording demonstrating a clear genetic deletion of modular MOR in LC. The recorded cell without functional MORs failed to respond to the application of DAMGO. (F) A schematic cartoon illustrating the viral strategies for the modular deletion of MOR in LC. (G) A timeline showing the experimental design for Hargreaves task in concert with pharmacological infusions. (H) A plot showing the impact of modular knockout of MOR in LC on pharmacological analgesia. Note that this genetic deletion disrupts the pharmacological effects led by DAMGO and Cocktail B but no other applications. Repeated measures two-way ANOVA followed by Bonferroni’s test, **p<0.01, ***p<0.001, ****p<0.0001. Only selective statistical comparisons are shown in H, please see **Table 5** for detailed statistics. All of data are represented as mean ± SD.

**Table 5.**
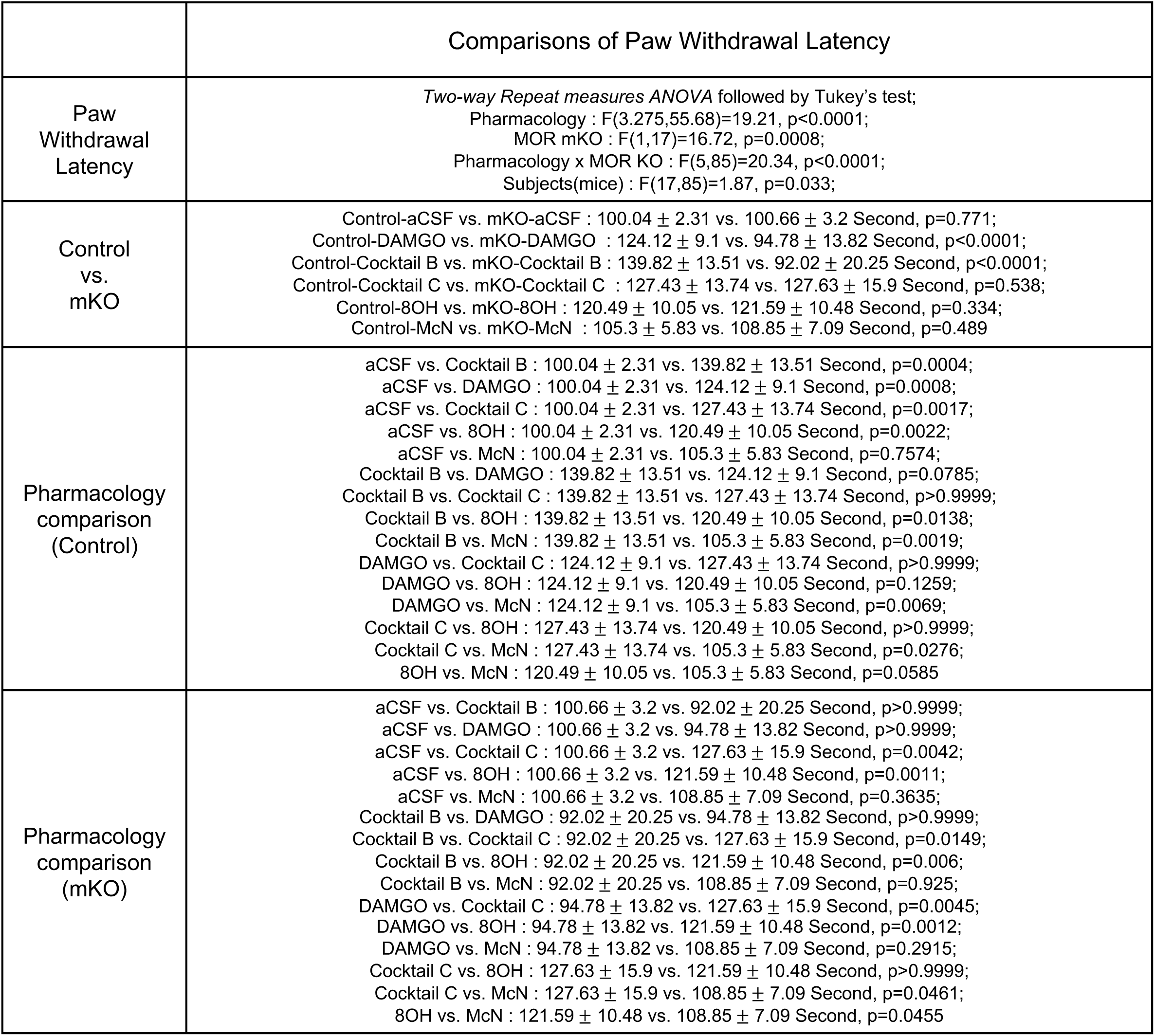

|  | Comparisons of Paw Withdrawal Latency |
| --- | --- |
| Paw Withdrawal Latency | *Two-way Repeat measures ANOVA followed by Tukey's test;*<br>Pharmacology : $F(3.275,55.68)=19.21$, $p<0.0001$;<br>MOR mKO : $F(1,17)=16.72$, $p=0.0008$;<br>Pharmacology x MOR KO : $F(5,85)=20.34$, $p<0.0001$;<br>Subjects(mice) : $F(17,85)=1.87$, $p=0.033$; |
| Control vs. mKO | Control-aCSF vs. mKO-aCSF : $100.04 \pm 2.31$ vs. $100.66 \pm 3.2$ Second, $p=0.771$;<br>Control-DAMGO vs. mKO-DAMGO : $124.12 \pm 9.1$ vs. $94.78 \pm 13.82$ Second, $p<0.0001$;<br>Control-Cocktail B vs. mKO-Cocktail B : $139.82 \pm 13.51$ vs. $92.02 \pm 20.25$ Second, $p<0.0001$;<br>Control-Cocktail C vs. mKO-Cocktail C : $127.43 \pm 13.74$ vs. $127.63 \pm 15.9$ Second, $p=0.538$;<br>Control-8OH vs. mKO-8OH : $120.49 \pm 10.05$ vs. $121.59 \pm 10.48$ Second, $p=0.334$;<br>Control-McN vs. mKO-McN : $105.3 \pm 5.83$ vs. $108.85 \pm 7.09$ Second, $p=0.489$ |
| Pharmacology comparison (Control) | aCSF vs. Cocktail B : $100.04 \pm 2.31$ vs. $139.82 \pm 13.51$ Second, $p=0.0004$;<br>aCSF vs. DAMGO : $100.04 \pm 2.31$ vs. $124.12 \pm 9.1$ Second, $p=0.0008$;<br>aCSF vs. Cocktail C : $100.04 \pm 2.31$ vs. $127.43 \pm 13.74$ Second, $p=0.0017$;<br>aCSF vs. 8OH : $100.04 \pm 2.31$ vs. $120.49 \pm 10.05$ Second, $p=0.0022$;<br>aCSF vs. McN : $100.04 \pm 2.31$ vs. $105.3 \pm 5.83$ Second, $p=0.7574$;<br>Cocktail B vs. DAMGO : $139.82 \pm 13.51$ vs. $124.12 \pm 9.1$ Second, $p=0.0785$;<br>Cocktail B vs. Cocktail C : $139.82 \pm 13.51$ vs. $127.43 \pm 13.74$ Second, $p>0.9999$;<br>Cocktail B vs. 8OH : $139.82 \pm 13.51$ vs. $120.49 \pm 10.05$ Second, $p=0.0138$;<br>Cocktail B vs. McN : $139.82 \pm 13.51$ vs. $105.3 \pm 5.83$ Second, $p=0.0019$;<br>DAMGO vs. Cocktail C : $124.12 \pm 9.1$ vs. $127.43 \pm 13.74$ Second, $p>0.9999$;<br>DAMGO vs. 8OH : $124.12 \pm 9.1$ vs. $120.49 \pm 10.05$ Second, $p=0.1259$;<br>DAMGO vs. McN : $124.12 \pm 9.1$ vs. $105.3 \pm 5.83$ Second, $p=0.0069$;<br>Cocktail C vs. 8OH : $127.43 \pm 13.74$ vs. $120.49 \pm 10.05$ Second, $p>0.9999$;<br>Cocktail C vs. McN : $127.43 \pm 13.74$ vs. $105.3 \pm 5.83$ Second, $p=0.0276$;<br>8OH vs. McN : $120.49 \pm 10.05$ vs. $105.3 \pm 5.83$ Second, $p=0.0585$ |
| Pharmacology comparison (mKO) | aCSF vs. Cocktail B : $100.66 \pm 3.2$ vs. $92.02 \pm 20.25$ Second, $p>0.9999$;<br>aCSF vs. DAMGO : $100.66 \pm 3.2$ vs. $94.78 \pm 13.82$ Second, $p>0.9999$;<br>aCSF vs. Cocktail C : $100.66 \pm 3.2$ vs. $127.63 \pm 15.9$ Second, $p=0.0042$;<br>aCSF vs. 8OH : $100.66 \pm 3.2$ vs. $121.59 \pm 10.48$ Second, $p=0.0011$;<br>aCSF vs. McN : $100.66 \pm 3.2$ vs. $108.85 \pm 7.09$ Second, $p=0.3635$;<br>Cocktail B vs. DAMGO : $92.02 \pm 20.25$ vs. $94.78 \pm 13.82$ Second, $p>0.9999$;<br>Cocktail B vs. Cocktail C : $92.02 \pm 20.25$ vs. $127.63 \pm 15.9$ Second, $p=0.0149$;<br>Cocktail B vs. 8OH : $92.02 \pm 20.25$ vs. $121.59 \pm 10.48$ Second, $p=0.006$;<br>Cocktail B vs. McN : $92.02 \pm 20.25$ vs. $108.85 \pm 7.09$ Second, $p=0.925$;<br>DAMGO vs. Cocktail C : $94.78 \pm 13.82$ vs. $127.63 \pm 15.9$ Second, $p=0.0045$;<br>DAMGO vs. 8OH : $94.78 \pm 13.82$ vs. $121.59 \pm 10.48$ Second, $p=0.0012$;<br>DAMGO vs. McN : $94.78 \pm 13.82$ vs. $108.85 \pm 7.09$ Second, $p=0.2915$;<br>Cocktail C vs. 8OH : $127.63 \pm 15.9$ vs. $121.59 \pm 10.48$ Second, $p>0.9999$;<br>Cocktail C vs. McN : $127.63 \pm 15.9$ vs. $108.85 \pm 7.09$ Second, $p=0.0461$;<br>8OH vs. McN : $121.59 \pm 10.48$ vs. $108.85 \pm 7.09$ Second, $p=0.0455$ |

## Discussion

Here we introduce a framework for novel, genetic tool-free neural circuit-selective pharmacology. Using *ex vivo* calcium imaging in concert with multiplexed pharmacology, we tested pharmacological responses during application of agonists targeting eighteen different GPCRs in LC-NE neurons. Next, we imprinted this pharmacological response profile onto efferent-defined LC modular organization. Co-labeling the LC-mPFC projection revealed differential pharmacological responses led by activation of the GPCR agonists. To test whether pharmacological, modular-selective control of the LC could elicit antinociception, we generated pharmacological dose-response curves using the thermal paw withdrawal assay in mice following local infusions of single compounds and cocktail combinations. Here we find infusion of a combination of GPCR agonists targeting MOR, 5HTR1a and mAChR1 caused a greater antinociception by shifting the neural activity away from LC-mPFC neurons. Furthermore, genetic exclusion of MOR in the LC-mPFC module disrupted antinociception from this pharmacological combination, but not when the cocktail did not activate MOR. This latter finding suggests the dependence of modular interaction underlying enhanced antinociceptive action mediated by activating multiple GPCRs within the LC.

### Technical potential and limitations

Here we demonstrate an *ex vivo* multiplexed pharmacological scan to quickly test multiple responses across many LC-NE neurons. We were able to deconvolute individual spikes during drug application using a machine learning-base model based on spontaneous fluctuating calcium signals. This technical combination has broad compatibility with cell types in different brain regions. For example, it would be straightforward to implement the same strategy in striatal cholinergic interneurons which also spontaneously fire around 0.5-5 Hz in normal slice preparation^78^. The firing frequency-dependent deconvolution allows us to dissect neural activity in compactly arranged nuclei without bias from bleaching issues. However, the requirement of stable, spontaneous firing also presents challenges when applying the approach to cells that do not fire under typical *ex vivo* conditions. This issue can be partially mitigated in other *ex vivo* or *in vivo* prepratations that allow repeated drug applications, such as superficial layers of spinal cord^79^. Although spontaneous activity can be generated by adjusting recording buffers^80–82^, technical difficulties remain when confronting cells with discrete firing modes at different membrane potentials like thalamic relay neurons^83,84^, or cells with adaptive spiking profiles^85,86^. Moreover, instrumental spatiotemporal resolution constraints in imaging data acquisition restrict compatibility with higher firing frequency neurons. Despite these technical limitations, this fast, efficient, and largely affordable survey of neural responses during pharmacological application provides a useful toolbox to understand the cell type-selective pharmacological action in physiological or pathological conditions.

### Interaction between GPCRs

To conduct the multiplexed agonist scan, we consecutively applied agonists targeting various GPCRs with sufficient washout periods for firing to return to baseline (**Table 1**). Despite our best efforts, this approach likely gives rise to inherent interactions between GPCR signaling cascades. We found that the application of some compounds caused robust firing effects that were resistant to the prolonged washing process (**Fig. S4**). These instances could arise from saturating doses, high ligand affinity, or the persistence of cellular mechanisms underpinning the effect on firing rate. In general, GPCRs activating Gi/o-coupled G-protein pathways were more difficult to entirely washout than Gq-coupled GPCRs (**Fig. S4**). The sustained potassium conductance through GIRK channels may underestimate responses from receptors using a similar mechanism^87,88^. The weak correlation between the baseline neural activity, DAMGO, and clonidine responses (**Fig. S5**) at least partially alleviates this possibility. It is also possible that the cell signaling pathway triggered activating G proteins might interact with subsequent GPCR activation^89,90^. For instance, MOR activation is known to cross-desensitize the alpha2-adrenergic receptor and somatostatin receptor^91^ and consecutive carbachol, a potent cholinergic agonist, applications desensitize mAChR1/3-mediated excitation in A7 noradrenergic neurons^92^. Phosphorylation of various second messengers like CREB, beta-arrestin, GRK, ERK in the LC could result in a complicated interaction between different receptors^57,71,93,94^. This issue was exaggerated between when receptor subtypes regulated by similar cellular mechanisms such as opioid receptors and muscarinic acetylcholine receptors were in the same group for pharmacological imaging. Furthermore, heteromerization occurs with discrete pairs of GPCRs and different ligands can be dependent on such heteromerization^95–99^. Whether such heteromers endogenously form in LC neurons is not yet clear, but our approach could be modified to identify such functions. To minimize bias from crosstalk between receptors we used subsaturating concentrations of agonists in randomized order. Agonists with irreversible effects (i.e., calcitonin, substance P and somatostatin) were pre-identified by cell-attached recordings and always applied last, prior to clonidine.

### Comparisons between optogenetics, chemogenetics and neural circuit-selective multiple pharmacological approach

Optogenetics and chemogenetics are powerful tools widely used in neuroscience for functional dissection of specific cell populations during discrete behaviors. Optogenetic tools such photoactivatable ion channels, pumps and receptors with various selectivity in light wavelength provide a toolbox for precise control of neural activity with excellent spatiotemporal resolution^3,4,100,101^. Chemogenetic tools such as DREADDs and PSAMs designed from mAChRs and ligand-gated channels, respectively, offer easy, long-term control of specific neural circuits^1,2,102,103^. Bridging these tools with photosensitive GPCRs enables spatiotemporal control of intracellular signaling cascades^104–106^. Nevertheless, a few concerns using optogenetics and chemogenetics have been reported. Proton pumps are limited in presynaptic inhibition, while chloride pumps and channels are often subject to rebound excitation^107,108^. While these issues are mitigated by photoactivatable Gi/o coupled GPCR, these tools are often more challenging to validate^98,101^. Additionally, the highly synchronous neural activity under optogenetic excitation might also interfere with, or otherwise non-physiologically entrain, intact brain oscillations. Ultrastructural evidence suggests the cellular location of chemogenetic constructs are largely affected tagged proteins^109^. The occupancy of intracellular signaling pathways due to overexpressed receptors could also interfere the intact transmission through endogenous GPCRs. Further research is needed to evaluate the cross desensitization of endogenous GPCRs during the activation of chemogenetic receptors as well as photoactivatable GPCRs. In contrast, our neural circuit-selective pharmacology approach merely relies on the intact cellular machinery triggered by endogenous GPCRs. Therefore, this approach is largely constrained by the nature of GPCR expression in given cell populations. Furthermore, presynaptic modulation can also be included in the pharmacological response during agonist applications^66,110–113^. Given that, in normal slice preparations, distal afferents remain mostly silent without axonal action potentials, we could underestimate the net effects caused by presynaptic action *in vivo*. Generally speaking, our approach likely yields a less potent and less selective control of neural activity compared to clear excitation or suppression using optogenetics and chemogenetics. However, several critical benefits still arise from the recruitment of endogenous GPCRs: (1) It helps to alleviate the off-target effects from overwhelming manipulations by optogenetics or chemogenetics while providing adequate, more physiological control on neural activity. (2) It sidesteps the need for genetic editing, lending itself to more translational applications. (3) It reduces unexpected effects from overexpression and off-target expression of foreign genes^9–12,114–116^. Nevertheless, substantial efforts are required to build our understanding of circuit-selective pharmacological profiles to expand this approach for non-genetic circuit-selective manipulations in other brain regions.

### Insights in multimodal analgesia

From a clinical perspective, combinations of multiple analgesic agents like opiates/opioids, acetaminophen, non-steroidal anti-inflammatory drugs, gabapentinoids, alpha-2 adrenoreceptor agonists, NMDA receptor antagonists, and serotonin-noradrenaline reuptake inhibitors are commonly used to treat postoperative and chronic pain conditions^117–119^. Taking advantage of additive or synergistic pharmacological antinociception, can alleviate non-desirable side effects such as nausea, vomiting and respiratory depression led by opioid administration^120–122^. However, most efforts toward multimodal analgesia are conceptually derived from a combination of known analgesics. Here we demonstrate a novel circuit-driven perspective to derive antinociception from multiple pharmacological approaches that may not directly link to antinociception alone (i.e., McN-A-343). The synergistic and enhanced antinociception driven by Cocktail A and Cocktail B, respectively, each which contain a MOR agonist, provides a route for reduced opioid use by leveraging the modular interaction in LC-NE system with other receptor ligands. Considering the prevalence of comorbid anxiodepressive disorders and sleep dysfunction during the chronification and maintenance of chronic pain^123–125^, the LC-NE system could be an excellent target for pain therapeutics due to its contribution to cognitive function, stress processing, and pain regulation^27–30,38,43,51,56,125^. Previous studies have shown decreased mRNA expression and faster desensitization of MOR in LC during the development of neuropathic pain^126,127^, and selective antagonism of alpha1/2 adrenergic receptors reversed neuropathic pain-induced depression^45^. Accordingly, one interesting future direction would be to test whether the neural circuit-selective agonist cocktails designed in this study could alleviate the comorbid hypersensitivity and anxiodepressive symptoms in chronic pain.

### GPCR-mediated control of functional LC modules

The rich expression of various GPCRs regulates LC-NE neurons to flexibly tune responses to diverse environmental stimuli^19,20,22–25^. A growing number of studies has interrogated the functional contribution of modular LC efferent architecture^27–30,42,54,56^, but relatively little effort has been made to determine how the afferent input rapidly shifts activity between LC functional modules. To address this issue, our work demonstrates differential modulation of LC modules by GPCR activation. We found that subsaturating concentrations of many GPCR agonists cause heterogenous effects across the LC. This variability in cellular response might also happen after the release of endogenous ligands *in vivo*, something difficult to precisely mimic exogenously. The lack of comprehensive understanding of presynaptic endogenous ligands and the postsynaptic locations of many GPCRs presents challenges in studying the physiological role of LC GPCRs. We recently showed that conditional knockout of MOR in noradrenergic neurons increased the baseline mechanical and thermal sensitivity, and this phenomenon was replicated by modular selective deletion of MOR in LC-mPFC neurons, demonstrating a modular role of LC MOR in pain regulation^42^. In this study, interestingly, the same LC-mPFC modular deletion of MOR disrupted Cocktail B-mediated antinociception. This was not the case for Cocktail C that did not include a MOR agonist. This latter observation suggests an active interaction between LC modules during pharmacological activation. Although the maximum antinociception was not increased, we also found a synergistic interaction between DAMGO and 8OH-DPAT from Cocktail A. This potentially opioid-sparing phenomenon might be due to rapid activation of shared downstream signaling pathways targeting Gi/o-coupled GPCRs and suggests a full recruitment of GIRK-mediated hyperpolarization^70,71,90^. In contrast, the greater antinociception from Cocktail B might reflect the competition between Gq- and Gi/o-coupled GPCRs on individual cells at lower doses, while higher doses unlock important modular interactions to enhance antinociception^91^. In summary, here we present a novel *ex vivo* imaging approach to rapidly guide the development of neural circuit-selective pharmacological approaches. Our work leveraged GPCR-mediated regulation of the LC-mPFC circuit to provide a clear target for multiplexed pharmacological antinociception.

## Materials and Methods

### Subjects

Both sex of adult C57BL/6J (JAX:000664), *Dbh*-Cre+/-(JAX:033951), *Oprm1*^fl/fl^ (JAX: 030074) mice over 8 weeks-old in age were used in this study. Animals were originally introduced from The Jackson Laboratory (Bar Harbor, ME, USA) then group housed and bred in a barrier facility with an ad libitum access to food pellets and water under a 12:12-hour light/dark cycle (lights on at 7:00 AM). Animal were transferred to the experimental facility and allowed to habituate for at least 2 weeks. All experiments and procedures were approved by the Institutional Animal Care and Use Committee of Washington University School of Medicine in accordance with National Institutes of Health guidelines.

### Stereotaxic Surgeries

Mice were anesthetized in an induction chamber (3% isoflurane) then mounted on a stereotaxic apparatus (Model 940, Kopf Instruments, CA, USA) and maintained at 1-2% isoflurane. Small craniotomies were performed and glass pipettes filled with tracers and viruses were slowly advanced to the brain area of interest as the following coordinates (in mm), mPFC : AP +2.0, ML 0.4, DV 0.9 & 1.8 from Bregma; LC: AP -0.9, ML 0.9, DV 2.9 from Lambda. 250nL of viruses were gently delivered through Nanoject III injector (Drummond Scientific Company, PA, USA) at rate of 40nL/mins. Tracers and viruses used included Ctb-CF594 (00072, Biotium, CA, USA), AAV1-syn-FLEX-jGCaMP8f-WPRE (162379-AAV1, Addgene, MA, USA), CAV-mCherry and CAV-PRS-Cre-V5 (PVM, Biocampus, Montpellier, France). Mice were allowed to recover for 2 weeks for neuronal tracing and at least 6 weeks for viral expression. For cannula implantation, 26-gauge bilateral guide cannulas coupled with blind cannulas (Protech Int. Inc) were slowly directed above the LC region (AP -0.9, ML 0.9, DV 2.8 from Lambda), 33-gauge injector cannulas (Protech Int. Inc) were then used in intra-cerebral infusion. Crowns of implantation were secured using C&B-Metabond (Parkell Inc., NY, USA) and super glue. Postoperative analgesia was made upon administration of carprofen (5 mg/kg) through oral tablets or subcutaneous injection for 3 days. Mice were allowed to have 2 weeks of recovery after surgery prior to behavioral tests. For the genetic deletion of modular MOR by retrograde virus, mice were allowed to have 7 weeks of recovery to complete the genetic knock out.

### Slice Preparation and Electrophysiology

Adult mice were deeply anesthetized with an i.p. injection of cocktail containing ketamine, xylazine & acepromazine then perfused with ice-cold slicing-aCSF consisting of (in mM) 92 N-methyl-d-glucose (NMDG), 2.5 KCl, 1.25 NaH_2_PO_4_, 10 MgSO_4_, 20 HEPES, 30 NaHCO_3_, 25 glucose, 0.5 CaCl_2_, 5 sodium ascorbate, 3 sodium pyruvate and 2 thiourea, oxygenated with 95% O_2_ and 5% CO_2_, then adjusted to pH 7.3-7.4 and 315-320 mOsm of osmolarity with sucrose. Brains were dissected rapidly then embedded and mounted with 2% agarose made in slicing-aCSF, coronal brainstem slices containing LC region were cut using vibratome (VF310-0Z, Precisionary Instruments, MA, USA). Slices were incubated in warm (32°C) slicing-aCSF for 30 minutes then transferred to holding-aCSF consisting of (in mM) 92 NaCl, 2.5 KCl, 1.25 NaH_2_PO_4_, 30 NaHCO_3_, 20 HEPES, 25 glucose, 2 MgSO_4_, 2 CaCl_2_, 5 sodium ascorbate, 3 sodium pyruvate and 2 thiourea, oxygenated with 95% O_2_ and 5% CO_2_, then adjusted to pH 7.3-7.4 and 310-315 mOsm of osmolarity with sucrose. The slice preparation procedure was modified from our previous study^42,113^. Slices were placed in a recording chamber mounted on an upright microscope (BX51WI, Olympus Optical Co., Ltd, Japan) with epifluorescent equipment and continuously perfused with warm (29-31°C) recording-aCSF consisting of (in mM) 124 NaCl, 2.5 KCl, 1.25 NaH_2_PO_4_, 24 NaHCO_3_, 5 HEPES, 12.5 glucose, 2 MgSO_4_ and 2 CaCl_2_, oxygenated with 95% O_2_ and 5% CO_2_, then adjusted to pH 7.3-7.4 and 305-310 mOsm of osmolarity with sucrose at a rate around 2.5-3mL/minute. Live tissue images were monitored by a digital CMOS camera (ORCA-Flash4.0LT, Hamamatsu Photonics, Japan) through a 40x water immersion objective lens (LUMPLFLN-40xW, Olympus, Tokyo, Japan) coupled with HC Image program (Hamamatsu Photonics, Japan). For cell-attached recordings, glass pipettes pulled by borosilicate glass capillary (GC150F-10, Warner Instruments, CT, USA) with a resistance around 5 MΩ when filled with recording-aCSF, either 470 or 505nm LED light from the epifluorescent system were used to identify the fluorescent expression in recorded cells. All electrophysiological data were collected by Multiclamp 700B amplifier (Molecular Devices, CA, USA) upon a low pass filtered at 2 kHz and digitalized at 10k Hz by Axon Digidata 1440A interface (Molecular Devices, CA, USA) coupled with Clampex software (Molecular Devices, CA, USA). Recording traces were collected and further analyzed by Clampx software, MATLAB (MathWorks, MA, USA) and GraphPad Prism9 (GraphPad Software, MA, USA).

### Multiplexed Pharmacological Scan

For *ex vivo* calcium imaging recordings, brain slices were transferred in the recording chamber with continuous perfusion of warm recording-aCSF as described above. The excitation and emission illuminations were delivered and collected using the live imaging system and epifluorescent equipment mounted on the Olympus BX51WI microscope. A 470 nm LED light under command of Axon Digidata 1440A interface coupled with Clampex software was used for field illumination of blue light at intensity of 0.5-1mW/mm^2^ based on the expression of calcium indicator. Fluorescent images were taken as 512 x 512-pixel square resolution after 4 x 4 binning covering 350 x 350 μm of FOV at 20 Hz with ≤ 50ms in exposure time (**Fig. 1B**), the focal plane was fixed at typically around 25-75 μm from the upper surface of recorded slices along the entire experiment, a sturdy anchor made of platinum was used to reduce unwanted motions and maximize the number of detectable cells in the FOV at selected focal plane. Upon the pharmacological applications, image stacks with length of 95 seconds was captured during baseline conditions or pharmacological bath perfusions. Simultaneous single or dual cell-attached recordings were made during the entire procedure to obtain raw material for model fitting or act as a live indicator of pharmacological action. The intensity of 470 nm excitation light was mildly raised depending on the bleaching of fluorescent signals along the pharmacological scanning protocol but remained consistent within each recording epoch to maintain the quality of imaging data.

### Imaging Data Processing and Spike deconvolution

All image stacks collected from the same slice were applied with 4 x 4 binning into 128 x 128 pixel-square then concatenated along the order of imaging recordings following the contrast adjustment to the same level across stacks by build-in functions of FUJI program^128^. The non-rigid motion correction was processed onto the image stacks by using NoRMCorre algorithm^129^ operated in MATLAB, followed by the baseline subtraction adopted a floating 2 second baseline along the whole stack then the first 5 seconds of each epoch were discarded due to the initial bleaching of fluorescent signals. The temporospatial information of each ROI was automatically extracted by CNMF-E, an open-source algorithm for analysis of microendoscopic calcium imaging data^130^, typically 15-30 ROIs were extracted per slice, temporal traces across ROIs were further separated by recording epochs for spike deconvolution (**Fig. S1A**). For the training of deconvolution model, ROIs represented the somata of recorded LC-NE neurons were aligned with simultaneous cell-attached recordings and formatted into algorithm-digestible files using MATLAB. CASCADE, a machine learning-based algorithm for spike inference from calcium signals using Python^59^, was used to fit the model for spike deconvolution based on training materials including around 6 hours of simultaneous recordings. For model evaluation, deconvoluted traces showing likelihood of spiking were calculated by the trained model upon the import of a pre-preserved set (15% of total materials, around 1 hour in length) of simultaneous recordings. Note that the recordings with relative high firing frequency were intentionally collected for model training and evaluation due to the smaller cell population in normal brain slice preparation (**Fig. 1C**). Spike inferences were made based on peaks and waveform of traces from both spike likelihood and raw calcium signal, then aligned to the ground truth representing the concomitant spikes by custom MATLAB scripts (**Fig. 1F, S2A-C**). Upon the alignment, pairs between ground truth and predicted spike with temporal difference < 2 image frames (±125ms) were considered as a correct prediction (**Fig. S2D**). For the spike deconvolution for pharmacological scan, calcium signals under baseline and pharmacological conditions were converted into traces of spike likelihood then the individual spike inferences were made as described above. To verify the cell identity by applications of clonidine, comparisons were made between inter-spike-intervals under baseline and clonidine application, simple judgements upon spike number were used when the firing rate was lower than 0.1 Hz. In a small subset of ROIs, spontaneous burst activities represented by the transient increase of bulk calcium signal as well as a short period of firing inhibition following the burst^113^, were manually removed prior to the deconvolution process.

### Immunohistochemistry

Mice were deeply anesthetized with an i.p. injection of cocktail containing ketamine, xylazine & acepromazine then perfused with ice-cold 4% paraformaldehyde in 0.1M PB. Brains were dissected and postfixed for 24 hours at 4°C with the same fixative. Upon the infiltration of 30% sucrose in 0.05M PB, frozen sections with 50μm thickness were cut through a sliding microtome (SM2000R, Leica, Germany). Sections were rinsed 3 times with PBS then incubated in blocking solution containing 2% bovine serum albumin (BSA) plus 5% normal goat serum (NGS) in PBST for 1 hour before transferred to PBST solutions containing primary antibodies plus 10% blocking solution for 24 hours at 4°C. Sections were then washed by PBS for 3 times and incubated in PBST solutions with secondary antibodies for 2-3 hours at room temperature followed by a rinse with PBS and wash with PB for 3 times. Sections were then mounted on glass slides with Vectashield mounting medium (Vector Labs, CA, USA). For post-hoc morphological reconstruction of slices used in the *ex vivo* calcium imaging recording, slices were fixed by 4% paraformaldehyde in 0.1M PB for few days and rinsed with PBS for 3 times. Slices were incubated in blocking solution consisting of 2% bovine serum albumin (BSA) plus 10% normal goat serum (NGS) in PBST for 1 hour followed by PBST solutions with primary antibodies plus 10% blocking solution for 72 hours at 4°C. Then slices were washed with PBS for 3 times and transferred to PBST solutions with secondary antibodies for 24 hours at 4°C followed by a rinse with PBS and wash for 3 times with PB. Slices were then incubated in RapiClear 1.47 (SUNJin Lab, Taiwan) for 1 hour and mounted with the same clearing agent for imaging scan. Images were collected using Leica confocal microscope (SP8, Leica, Germany), z-stacks with 2-3 μm intervals were scanned for the reconstruction of tissue morphology of recorded slices after post-hoc immunohistochemistry. The usage of antibodies is listed below:

| Antibody | Catalog | Supplier | Dilution | Figure |
| --- | --- | --- | --- | --- |
| mouse anti-TH | MAB318 | MilliporeSigma, MA, USA | 1:500 | 3C-E, 5C, S15 |
| rabbit anti-TH | AB152 | EMD Millipore, MA, USA | 1:2000 | S8, 4B, S17 |
| chicken anti-GFP | GP1010 | Aves Labs, CA, USA | 1:1000 | 3C-E |
| rabbit anti-Ctb | ab34992 | Abcam, Cambridge, UK | 1:500 | 3C-E, S8 |
| rabbit anti-mCherry | 600-401-P16 | Rockland, NJ, USA | 1:1000 | 5B-C, S15 |

### Hargreaves thermal paw withdrawal assay and intracerebral Infusion

Mice were allowed to daily habituate with Hargreaves apparatus (IITC Life Science, CA, USA) for a week with 1 hour per day. Prior to behavioral examinations, mice were placed in the apparatus for 30 minutes for habituation. Upon thermal stimulation, hind paw withdrawal behavior was identified as the paw being removed from the glass surface of Hargreaves apparatus and the maximum duration was set at 20 seconds. Intensity of stimuli were adjusted between 25 to 40% of setting to obtain a paw withdrawal latency around 9 to 12 seconds for each mouse, measurements were made with this fixed intensity for the entire protocol. For the examinations coupled with intra-cerebral infusions, injector cannulas were connected to 2.0 µL Hamilton syringe (Neuros Model 7002 KH, Hamilton, NV, USA) mounted on a syringe pump (70-4500, Harvard Apparatus, MA, USA) and pre-filled with recording-aCSF based pharmacological solutions. Mice were connected to the injector cannulas before habituation session, 250 nL of solutions were infused at a rate of 50 nL/mins. During baseline measurements for each trial, mice didn’t receive any infusion but still connected to the setup. At least 3 days were allowed to washout the pharmacological effect before next tests. Measurements were made between 5-30 minutes after the end of infusions with 5 minutes intervals, results from both paws were averaged for further analysis.

### Isobolographic Analysis

The isobolographic analysis was used to evaluate the drug interaction within pharmacological combinations^74,131,132^. Shortly, EC_50_s were first calculated by a non-linear regression adopting the three-parameter dose-response formula as a built-in function in GraphPad Prism 10.0. Isobolograms were drawn by the EC_50_s of pharmacological effect from single compound infusions, the theoretical EC_50_s and variance were calculated on the additive isobole depending on the recipe of drug combinations. An additive interaction is considered when the experimental EC_50_ shows no difference to the theoretical EC_50_, whereas the significantly smaller or larger experimental EC_50_ compared to theoretical EC_50_ denote a synergism or antagonism, respectively.

### Open field test

Mice were allowed to habituate to the test room for 1 hour prior to the test then connected to infusion setup and an additional 15 minutes of habituation was made. Mice then received 250 nL solutions at rate of 50 nL/min with either vehicle or 10μM Cocktail B, 5 minutes were allowed for the diffusion of injected solutions. Mice were individually placed into a 50 cm x 50 cm x 30 cm square acrylic box for 15 minutes. The behavioral activity was recorded through a Google Pixel 3 XL and the video. Traveled distances were further analyzed via Ethovision XT 13 (Noldus Information Technology, The Netherlands).

### Statistics

All data are expressed as mean ± SD except the isobologram with SEM demonstrations (**Fig. 4I**). One-sample t-Test, Student’s t test, paired t test, One-way ANOVA or Two-way ANOVA followed by posthoc tests were used to analyze data that pass the Shapiro-Wilk normality test. Otherwise, nonparametric tests were used. Statistical significance was indicated as ∗p < 0.05, ∗∗p < 0.01, ∗∗∗p < 0.001, ∗∗∗∗p < 0.0001 and ns indicates not significant. Statistical comparisons were conducted in GraphPad Prism 10.0. All statistical tests performed as part of this manuscript are available in **Supplemental Datasheet 1.**

## Supporting information

Supplemental Datasheet 1

Supplemental Datasheet 2

## Acknowledgements

We thank the other members of the Al-Hasani and McCall labs, particularly Jenny R. Kim, Manish K. Madasu, and Rui-Ni Wu for helpful feedback on this project. Special thanks to Patricia Jensen for the *Dbh-*Cre mice and EJ Kremer for the CAVs. This work was financially supported by the National Institutes of Health (R01NS117899; R01NS135401), the McDonnell Center for Systems Neuroscience (J.G.M.), and the Rita Allen Foundation (J.G.M.) with added financial help from the Open Philanthropy Project (J.G.M.). We would like to acknowledge biorender.com for figure cartoons, the Washington University School of Medicine Hope Center for Neurological Disorders viral vector core, and the Osage Nation, Missouria, Illinois Confederacy and many other tribes as the ancestral, traditional, and contemporary custodians of the land where this work was conducted.

## Author contributions

C.C.K. and J.G.M conceived the project and designed the detailed experimental protocols. C.C.K. performed the experiments and analyzed the data. J.G.M acquired funding and provided research supervision. C.C.K. and J.G.M wrote and edited the paper.

## Data availability

All data presented in this manuscript is available in **Supplemental Datasheet 2.**

## Code availability

The custom code used in this manuscript is included in Supplemental information.

---

**Figure S1.**
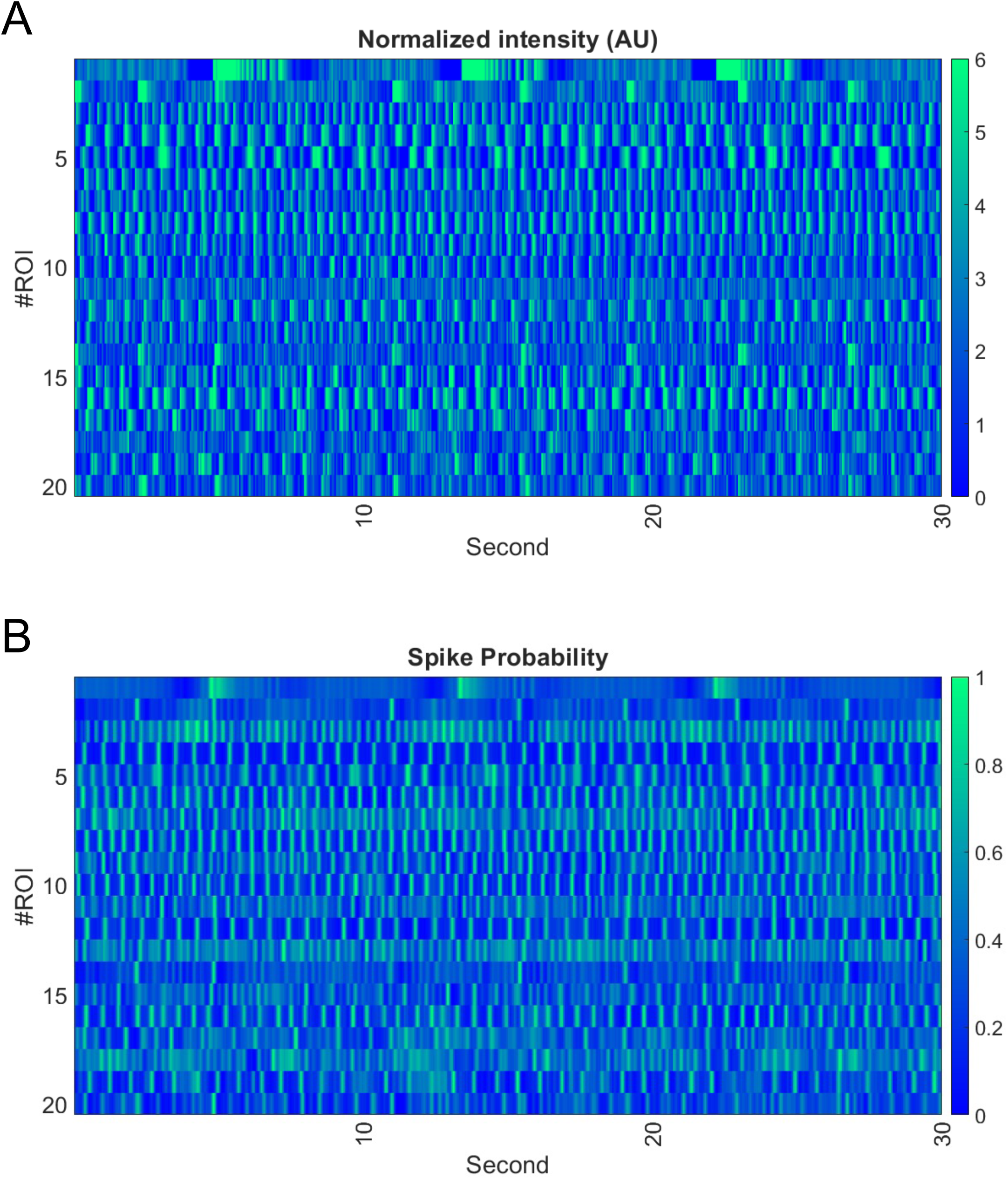
Conversion of calcium signal into spike likelihood. (A) A color map illustrating the raw calcium signals across 20 ROIs extracted from the same slice. (B) A color map illustrating traces of spike probability converted by the data shown in A by pre-trained deconvolution model.

**Figure S2.**
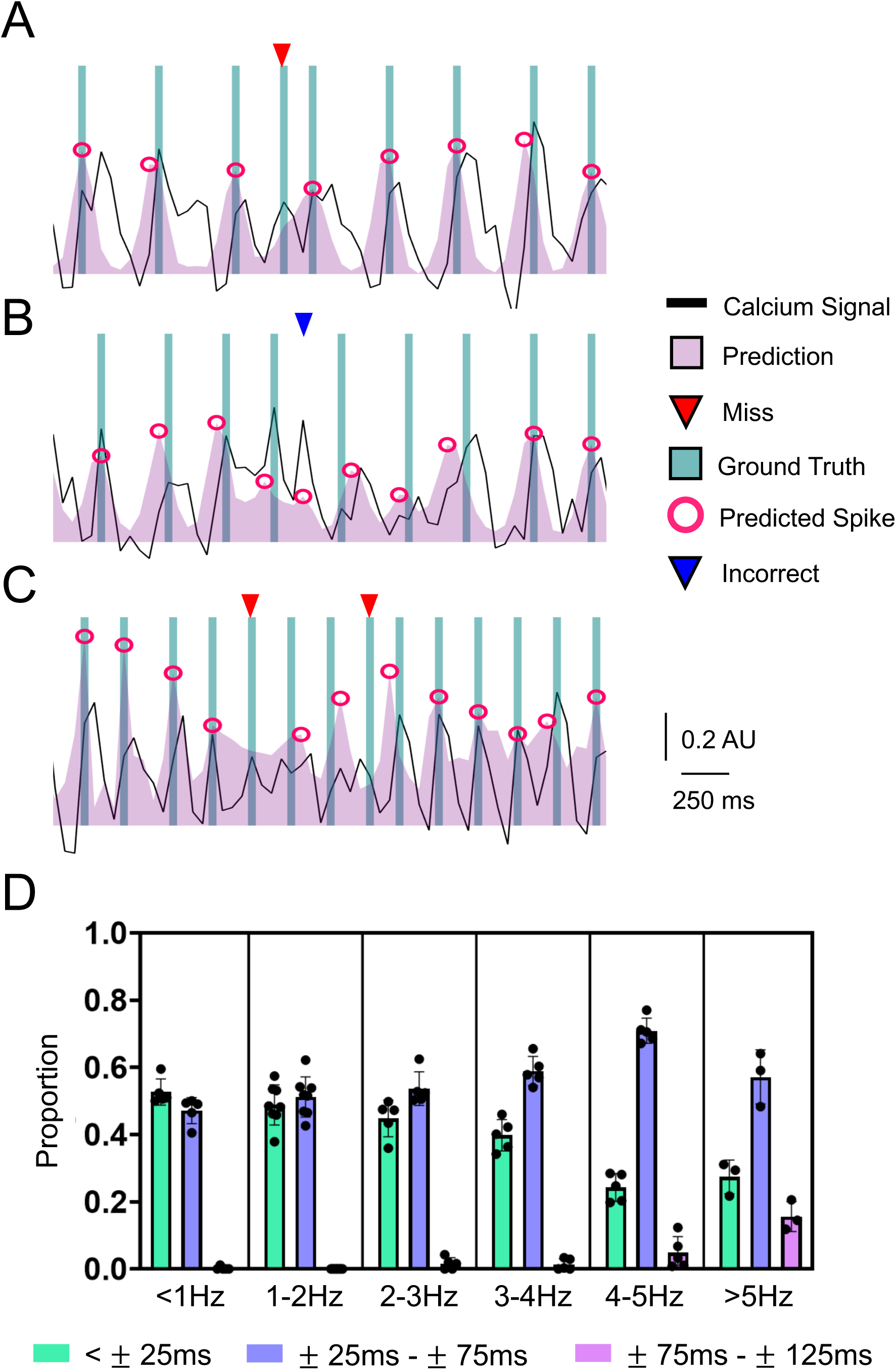
Prediction error in spike inference. (A) An example of spike missing as a deconvolution failure. (B) An example of erroneous prediction as a deconvolution failure. (C) An example showing the increased possibility of spike missing when cells fired action potential in higher rate. (D) A plot summarizes the proportional distribution of temporal difference between predictive and real spikes along the action potential firing rate. All of data are represented as mean ± SD.

**Figure S3.**
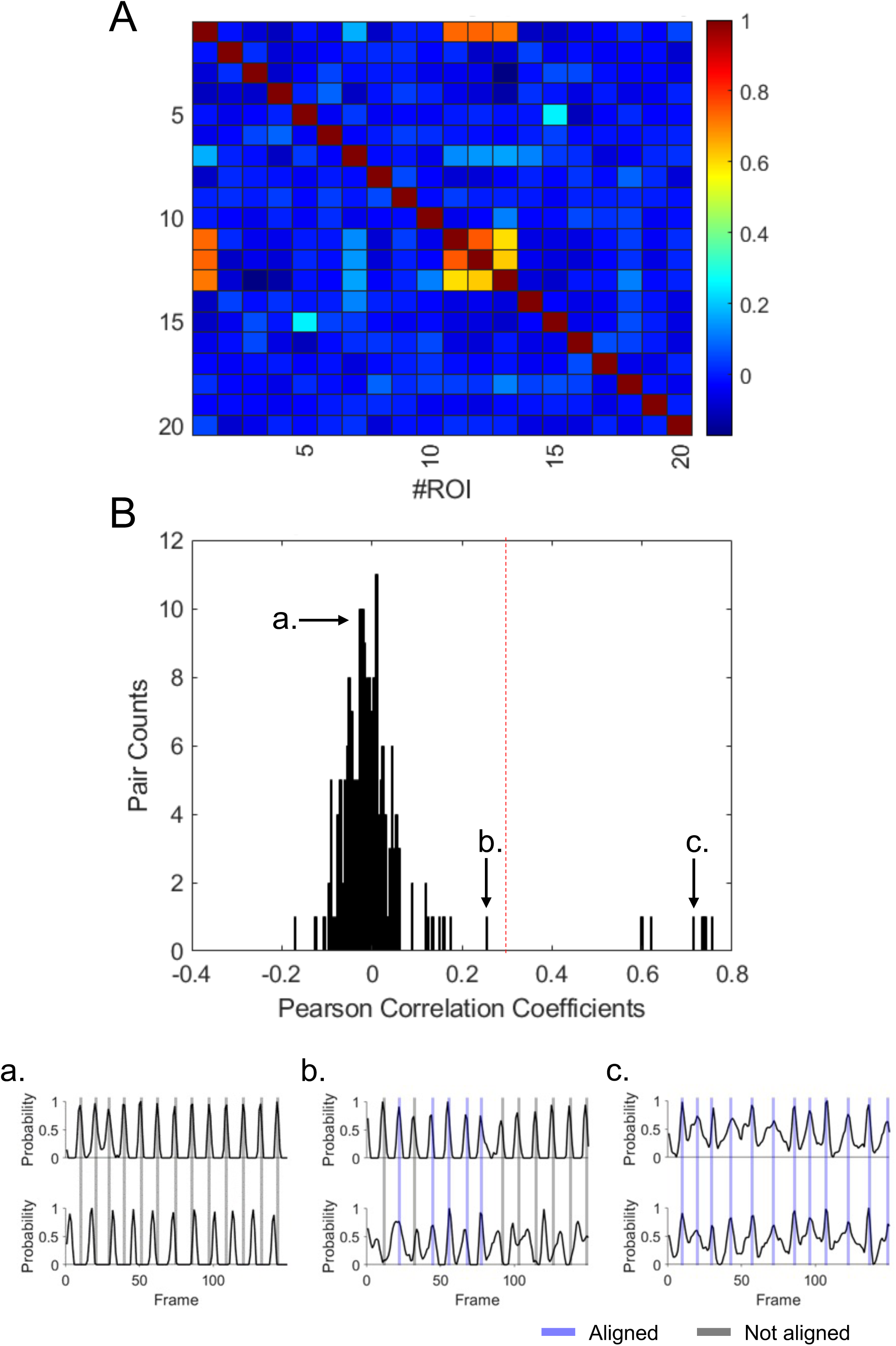
Identification of repeated ROIs. (A) A cross-correlation matrix from 20 ROIs collected from the same slice, note the high correlation coefficient in some of ROI pairs. (B) A plot demonstrating the distribution of correlation coefficients of ROI pairs shown in A and small panels below showing the none (left), partial (middle) and complete (right) synchronicity of predictive spikes.

**Figure S4.**
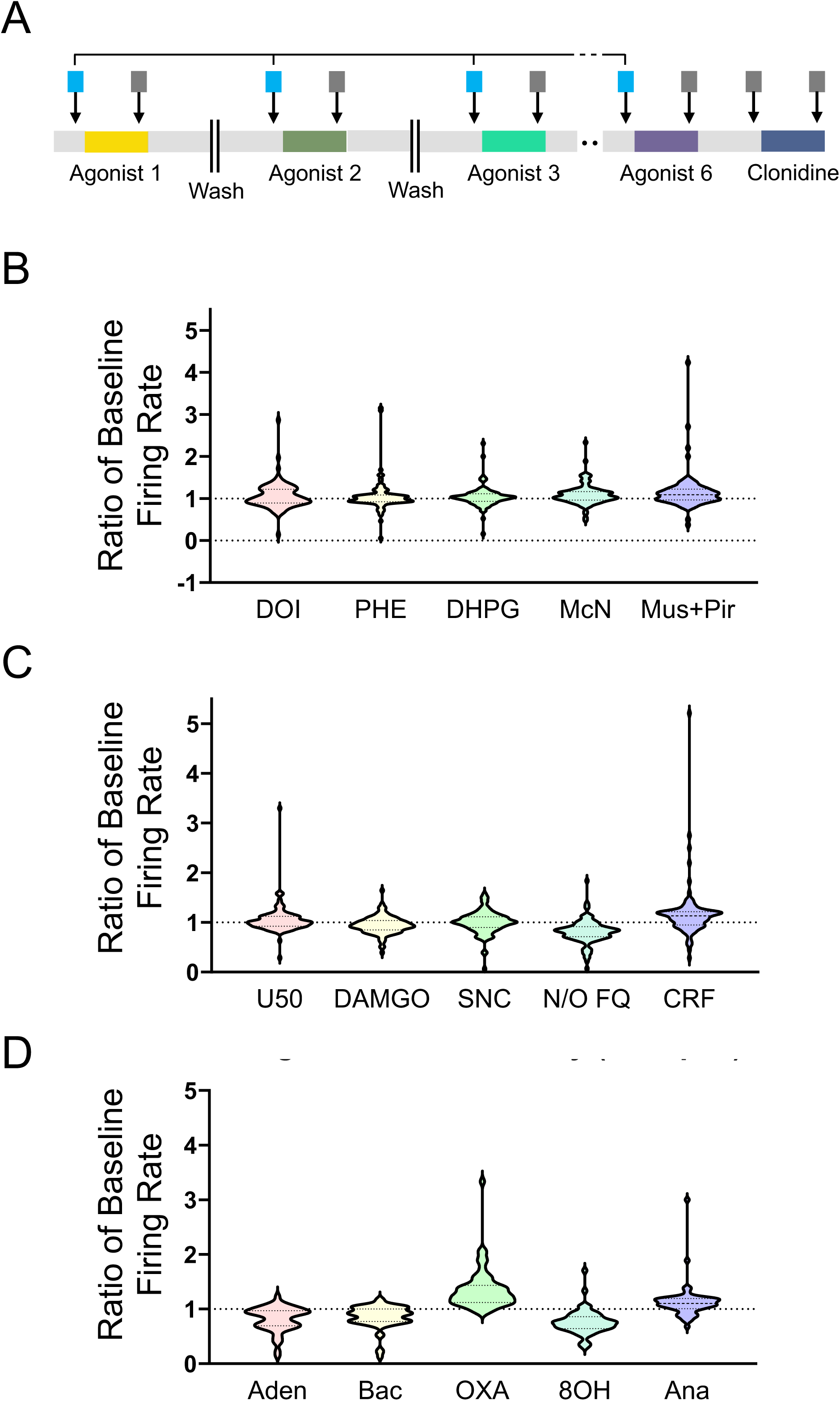
Washout efficiency across pharmacological recordings. (A) A schematic diagram demonstrating the quantification of wash efficiency as comparisons between baseline firing frequency. (B-D) Summarized results of drug washout efficiency across three groups of agonists. The wash efficiency is calculated by the ratio between firing rates in baseline epochs of adjacent applications.

**Figure S5.**
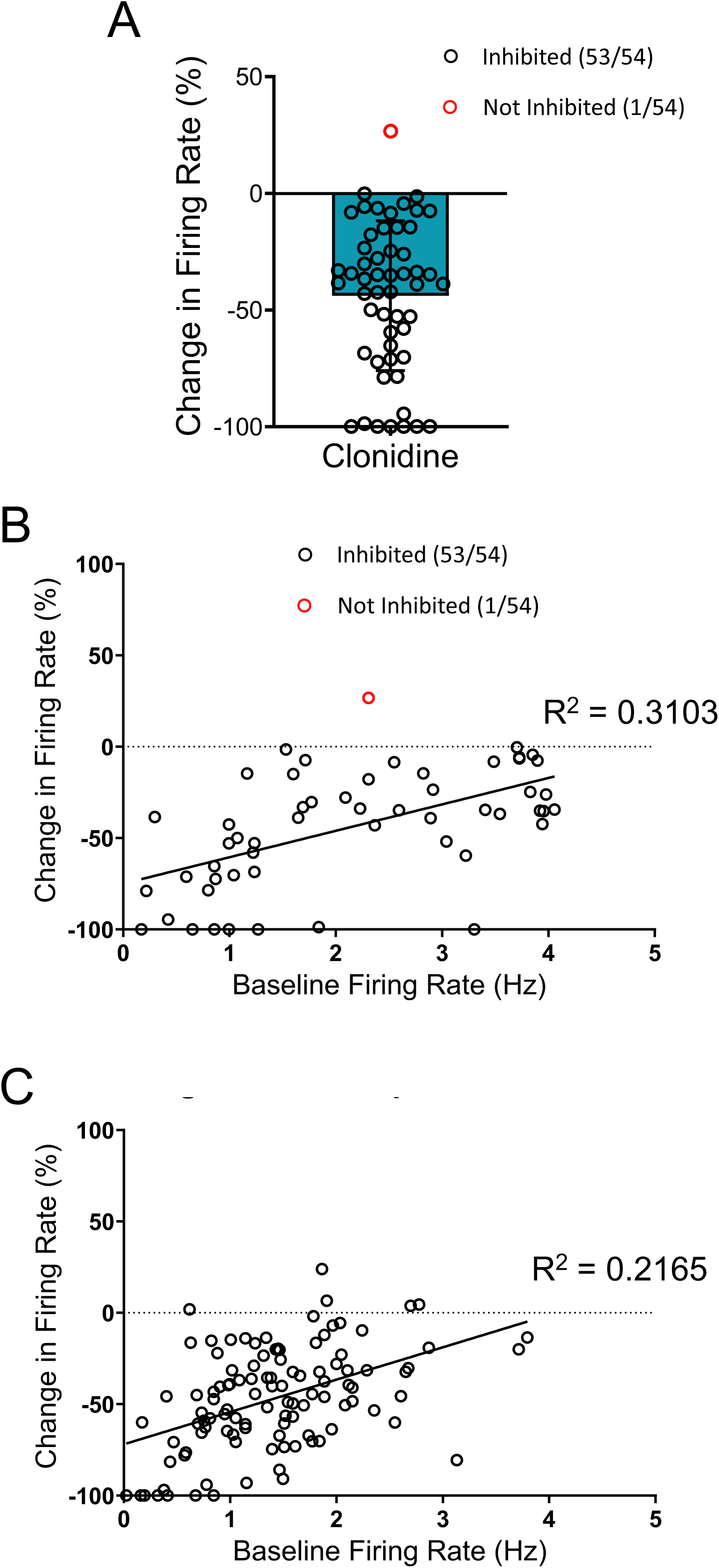
Relationship between basal firing rate and pharmacological effects from clonidine and DAMGO. (A) A plot showing the pooled results of pharmacological inhibition upon the application of clonidine, note that the red dot indicates a ROI that didn’t respond to the application. (B) A scatter plot displaying a neutral relationship between basal firing rate and the pharmacological inhibition from clonidine using data shown in A. (C) A scattered plot displaying the weak relationship between basal firing rate and the pharmacological effect from DAMGO. Data represented as mean ± SD.

**Figure S6.**
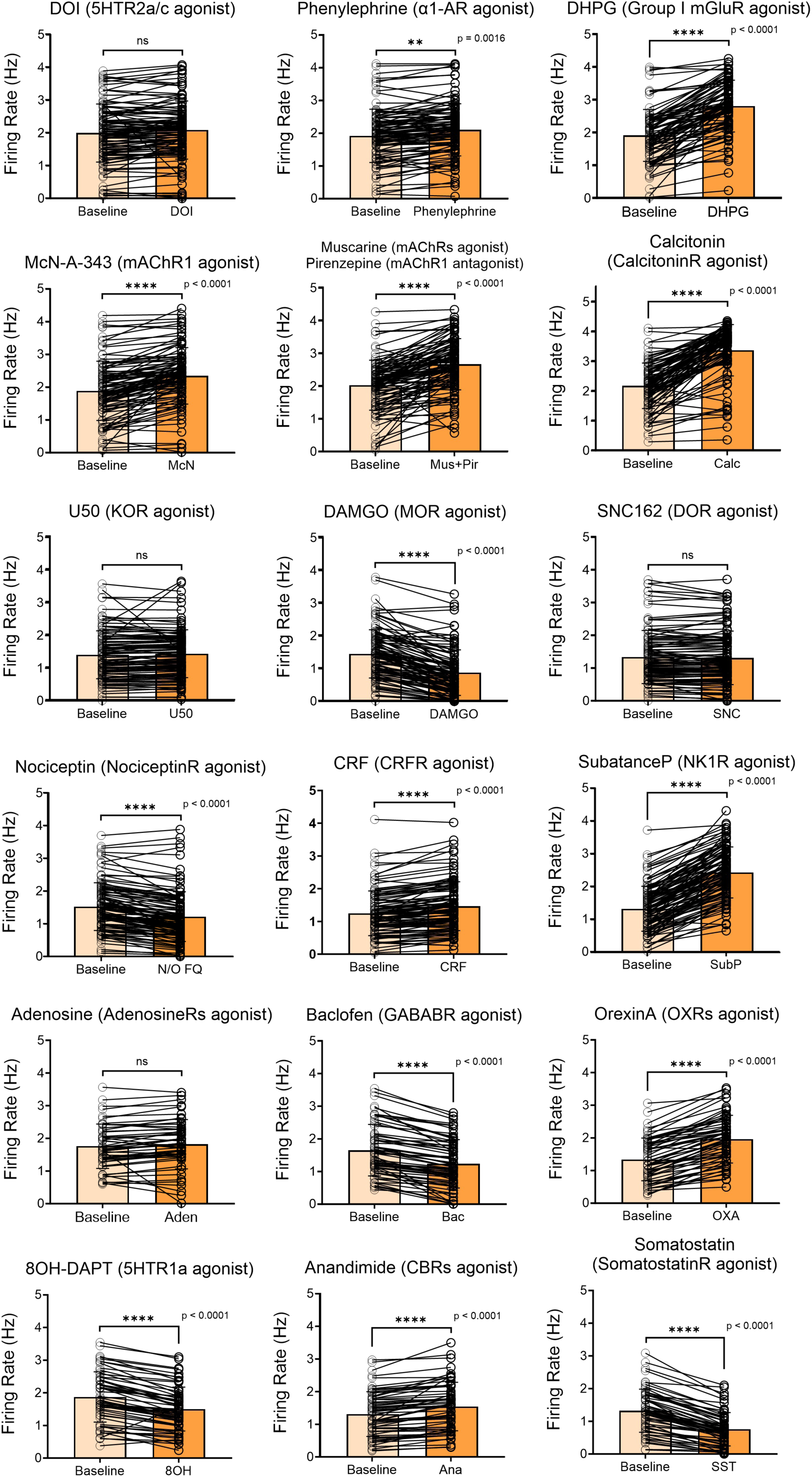
Pharmacological effects of LC-NE GPCR activation. The list of plots demonstrating the pharmacological effects in firing rate led by applications of agonists targeting 18 GPCRs. All of data are represented as mean ± SD. **p<0.01, ****p<0.0001, ns = not significant., please see **Table 2** for statistical details.

**Figure S7.**
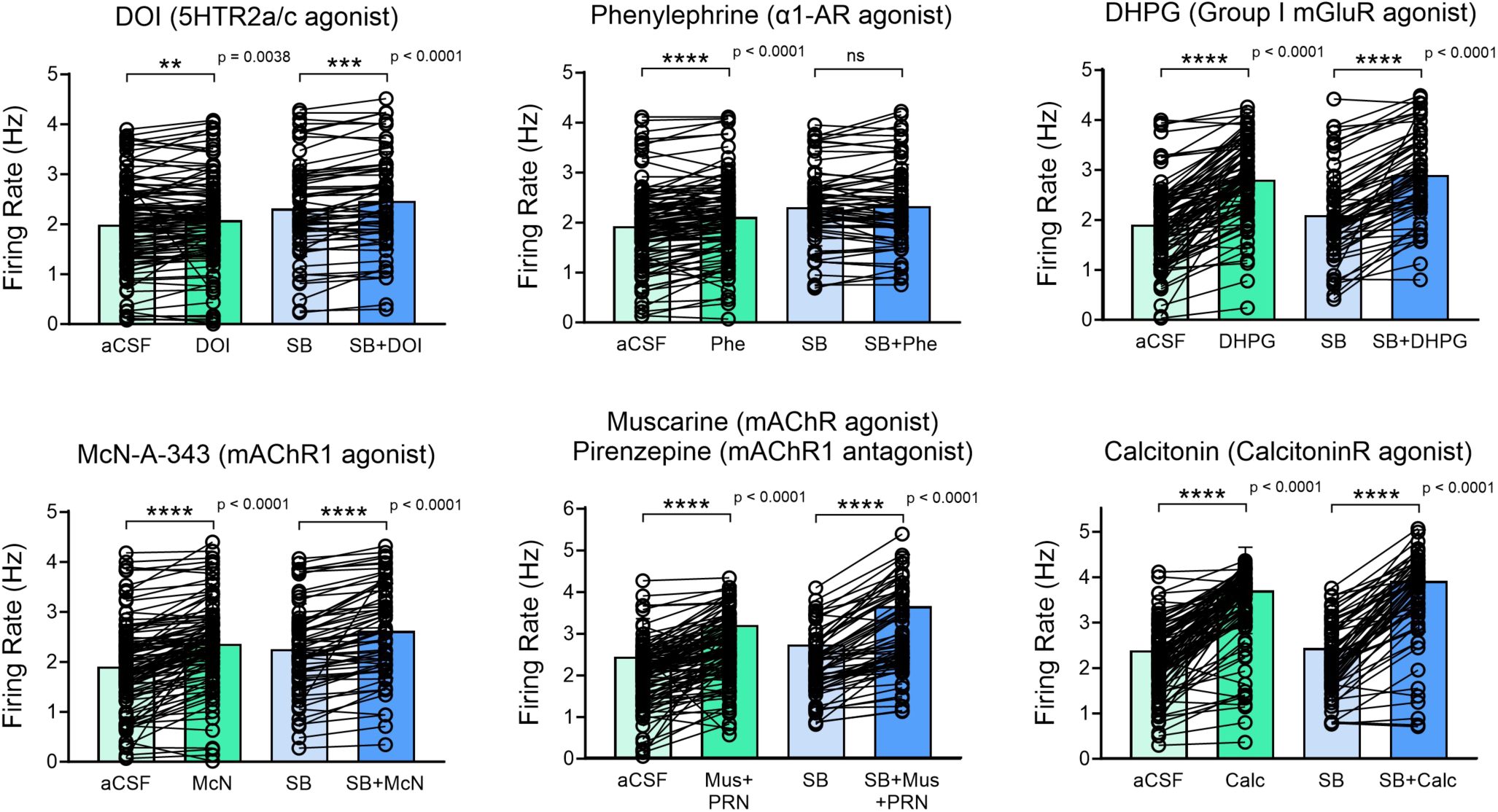
GPCR-mediated presynaptic modulation onto LC-NE neurons. The list of plots demonstrating the pharmacological response of Group I GPCRs under conditions of normal recording-aCSF perfusion and pre-administration with synaptic blockers. All of data are represented as mean ± SD, **p<0.01, ***p<0.001, ****p<0.0001, ns indicating not significant. Please see **Supplementary Datasheet 1** for detailed statistics.

**Figure S8.**
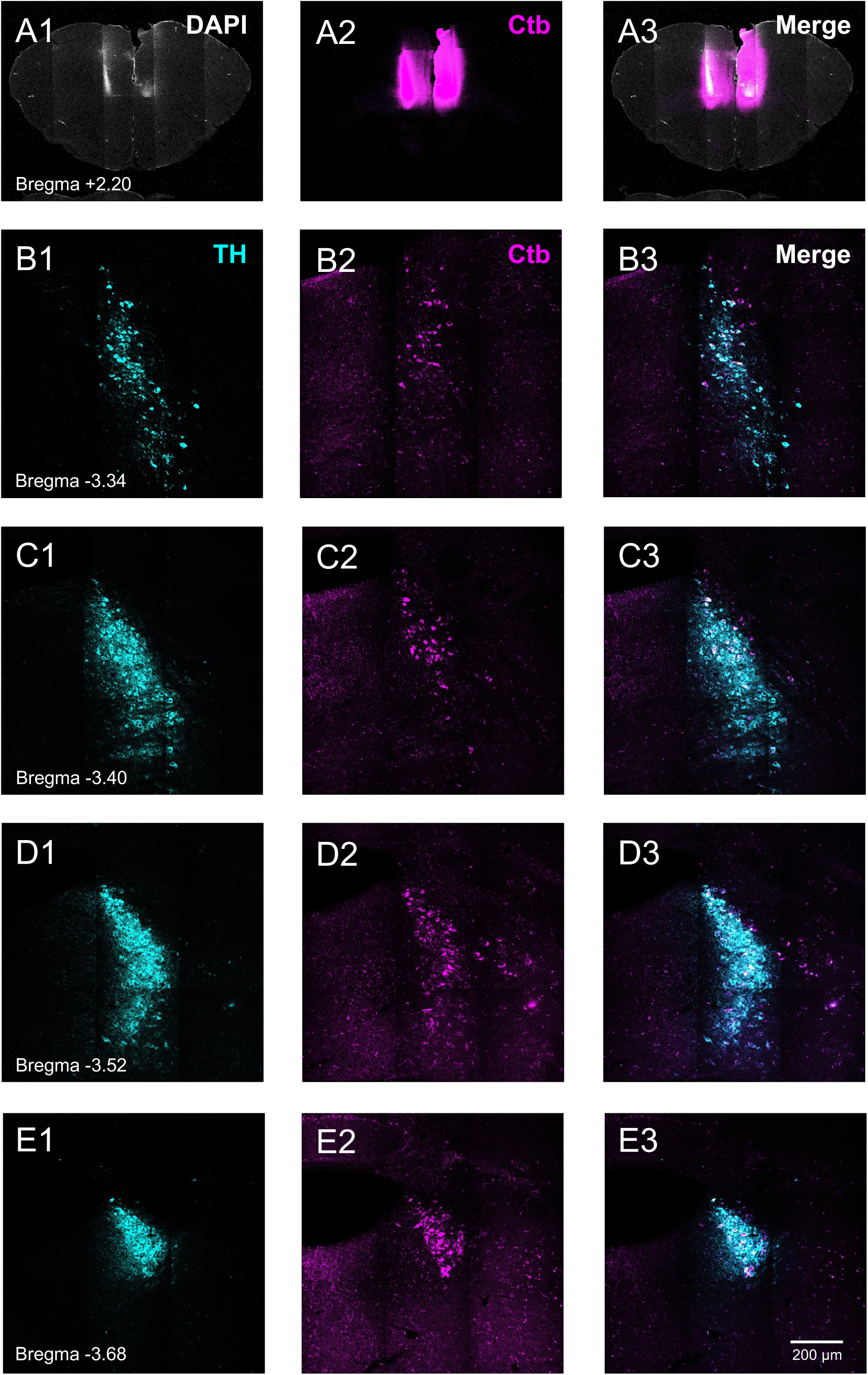
Distribution of mPFC-projecting LC-NE neurons revealed by retrograde tracer. (A) Fluorescent images showing the injection of Ctb594 into bilateral mPFC. (B-E) Fluorescent images displaying the distribution of mPFC-innervating LC-NE neurons with colocalization of TH and Ctb immunoreactive signals indicated by cyan (TH) and magenta (Ctb) colors. Note the fewer cells showing Ctb in the rostroventral part of LC.

**Figure S9.**
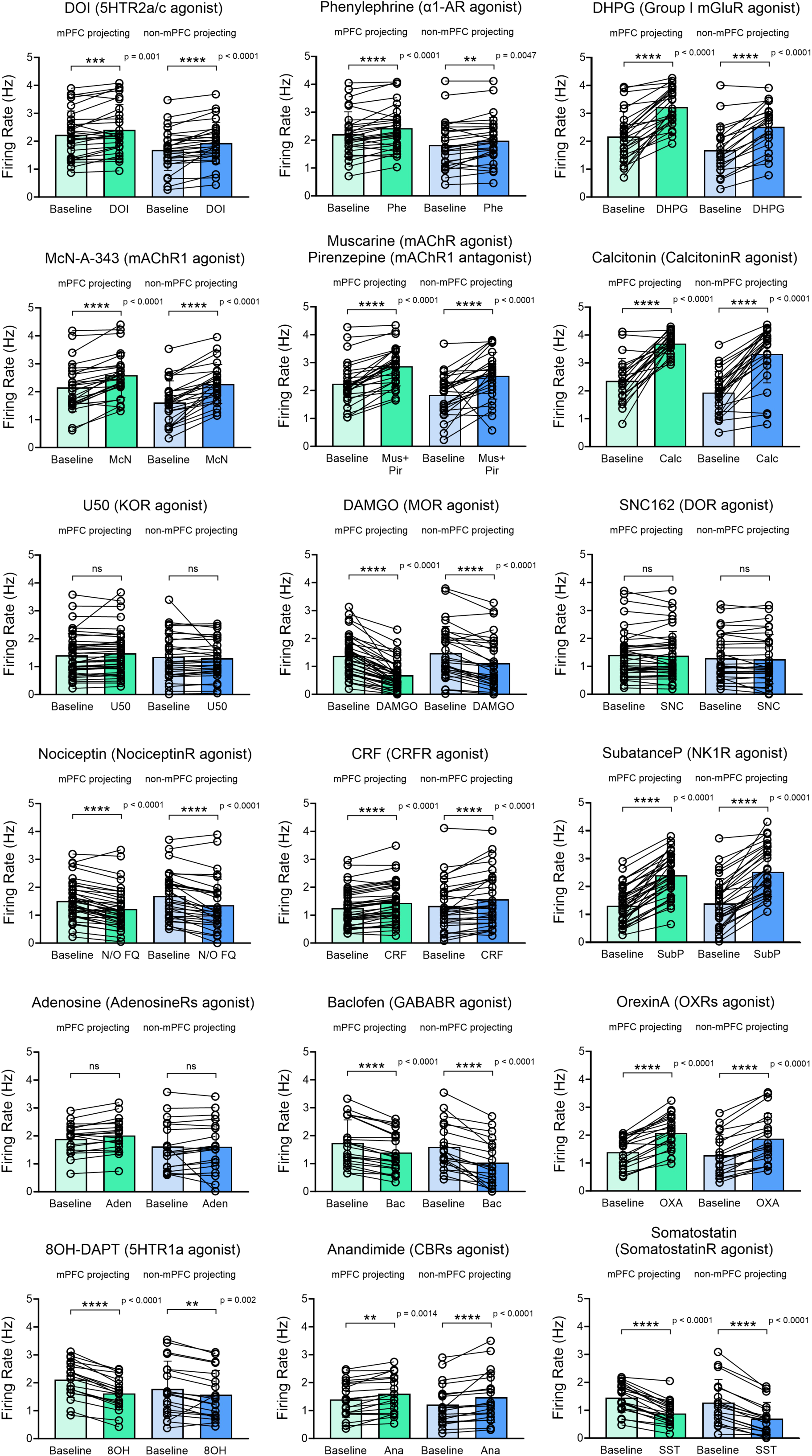
Raw data of pharmacological effects across LC modular organization. The list of plots demonstrating the pharmacological effects in firing rate by agonists targeting 18 GPCRs across LC modules. All of data are represented as mean ± SD. **p<0.01, ***p<0.001, ****p<0.0001, ns = not significant. Please see **Table 3** for statistical details.

**Figure S10.**
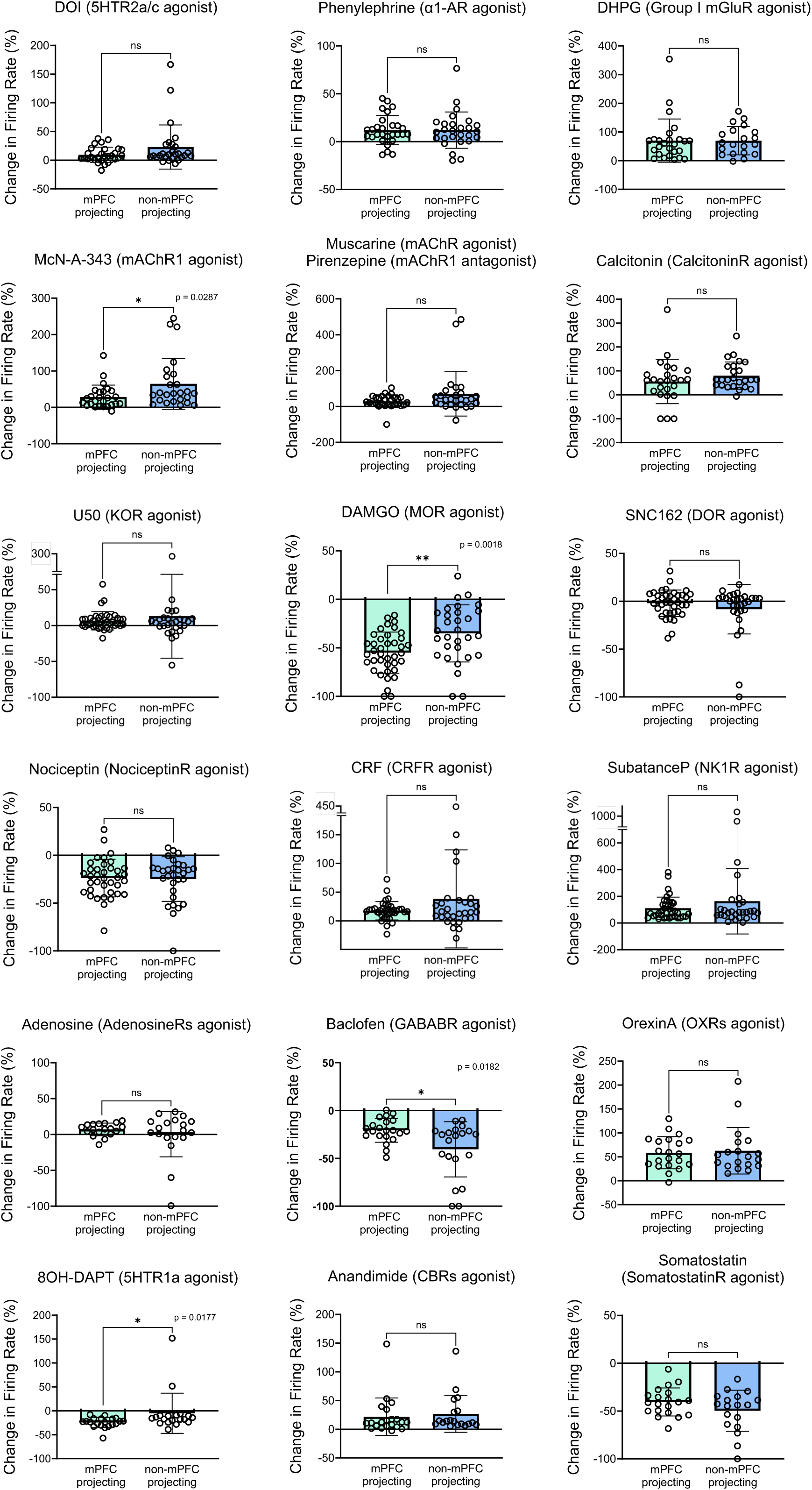
Comparison of pharmacological responses across LC modules. The list of plots demonstrating the percentile of changes in firing rate by agonists targeting 18 GPCRs across LC modules. All of data are represented as mean ± SD, *p<0.05, **p<0.01, ns = not significant, please see **Table 4** for statistical details.

**Figure S11.**
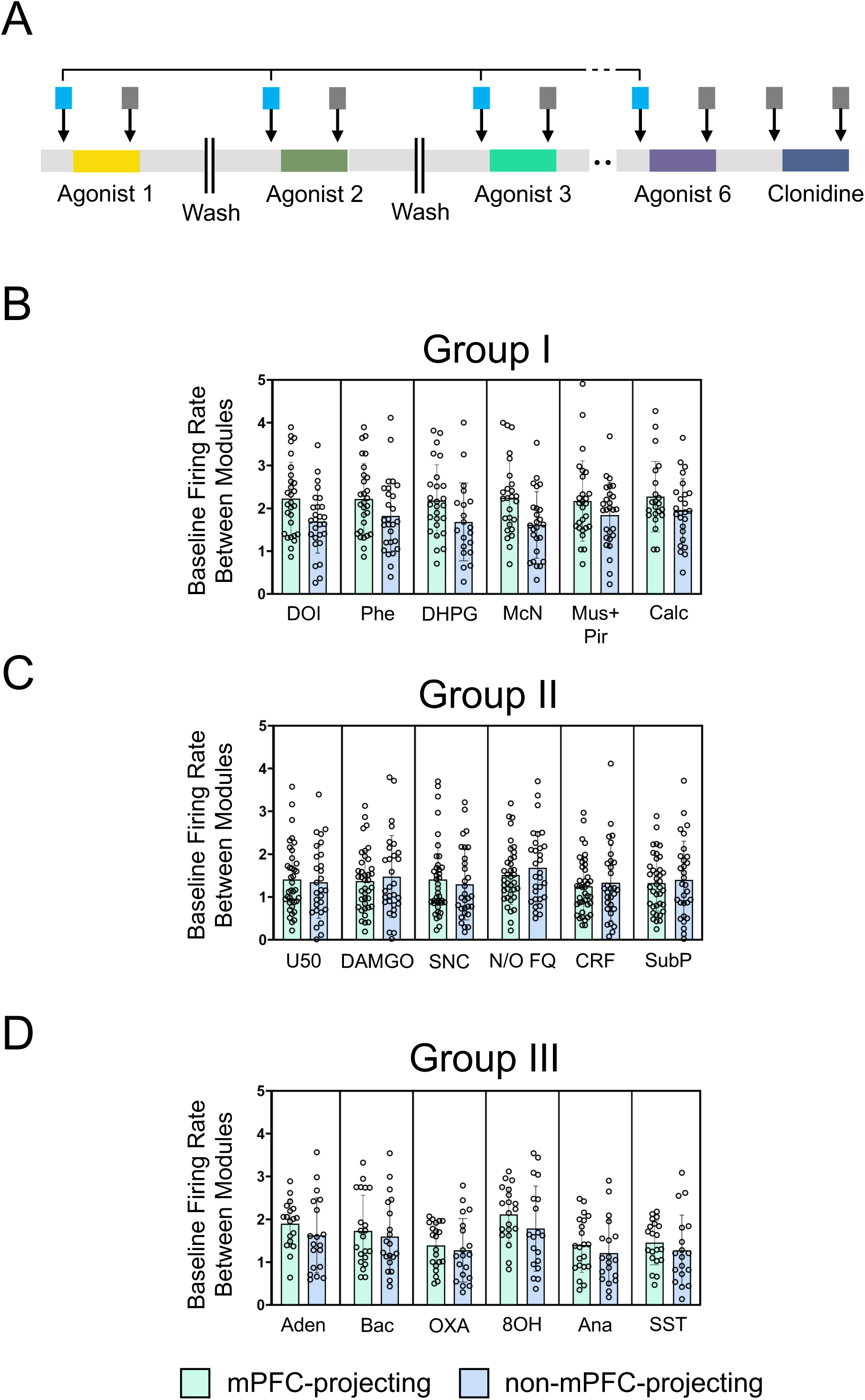
Comparisons of basal firing rate during modular pharmacological scan between LC modules. (A) A schematic diagram demonstrating the comparisons of baseline firing frequency between LC-NE neurons in different modules. (B-D) Plots showing the firing rate in baseline epochs across LC modules during multiplexed pharmacological scan. No significant difference is found among all of comparisons. Please see **Supplementary Datasheet 1** for detailed statistics. All of data are represented as mean ± SD.

**Figure S12.**
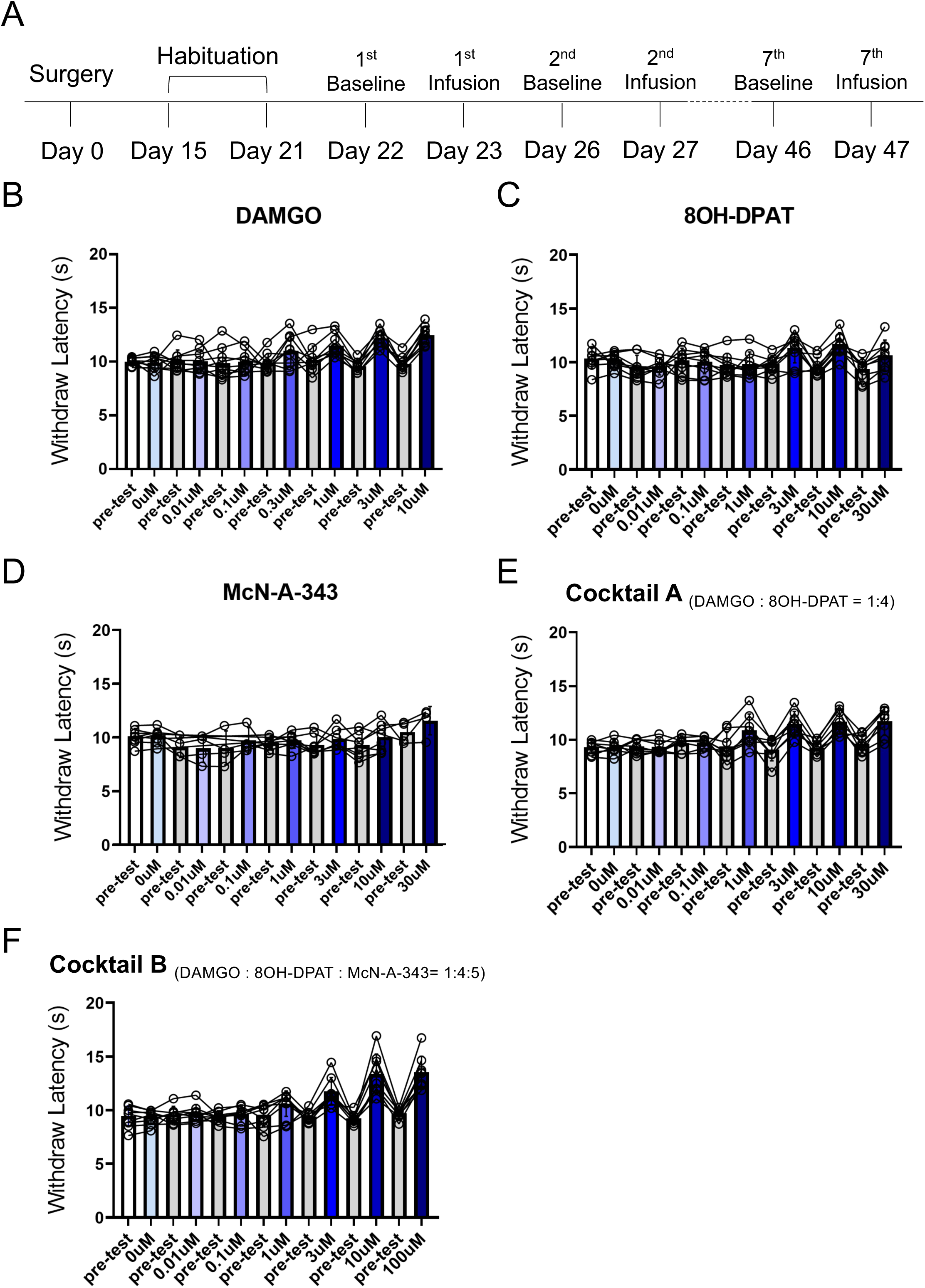
Hargreaves assay in concert with pharmacological infusions toward LC. (A) A timeline showing the experimental design. (B-F) Plots showing raw data of behavioral examinations, baselines for trials of each pharmacological dosage are collected from pre-test measurements.

**Figure S13.**
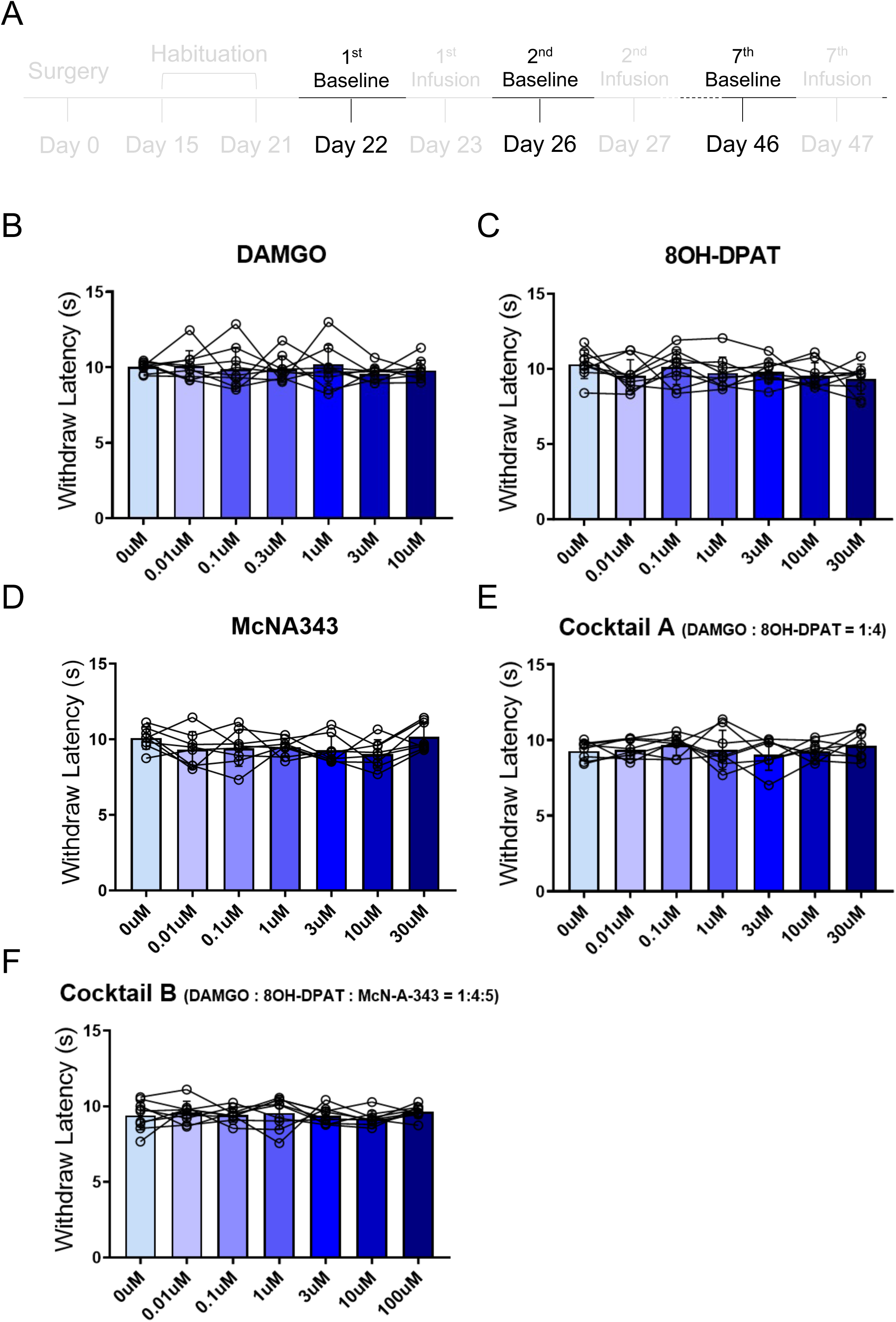
Comparisons across baseline tests. (A) A timeline showing the experimental design, only baselines are used for the comparisons. (B-F) Plots showing baseline behavioral examinations across the entire protocol. No significant difference was found. Please see **Supplementary Datasheet 1** for detailed statistics.

**Figure S14.**
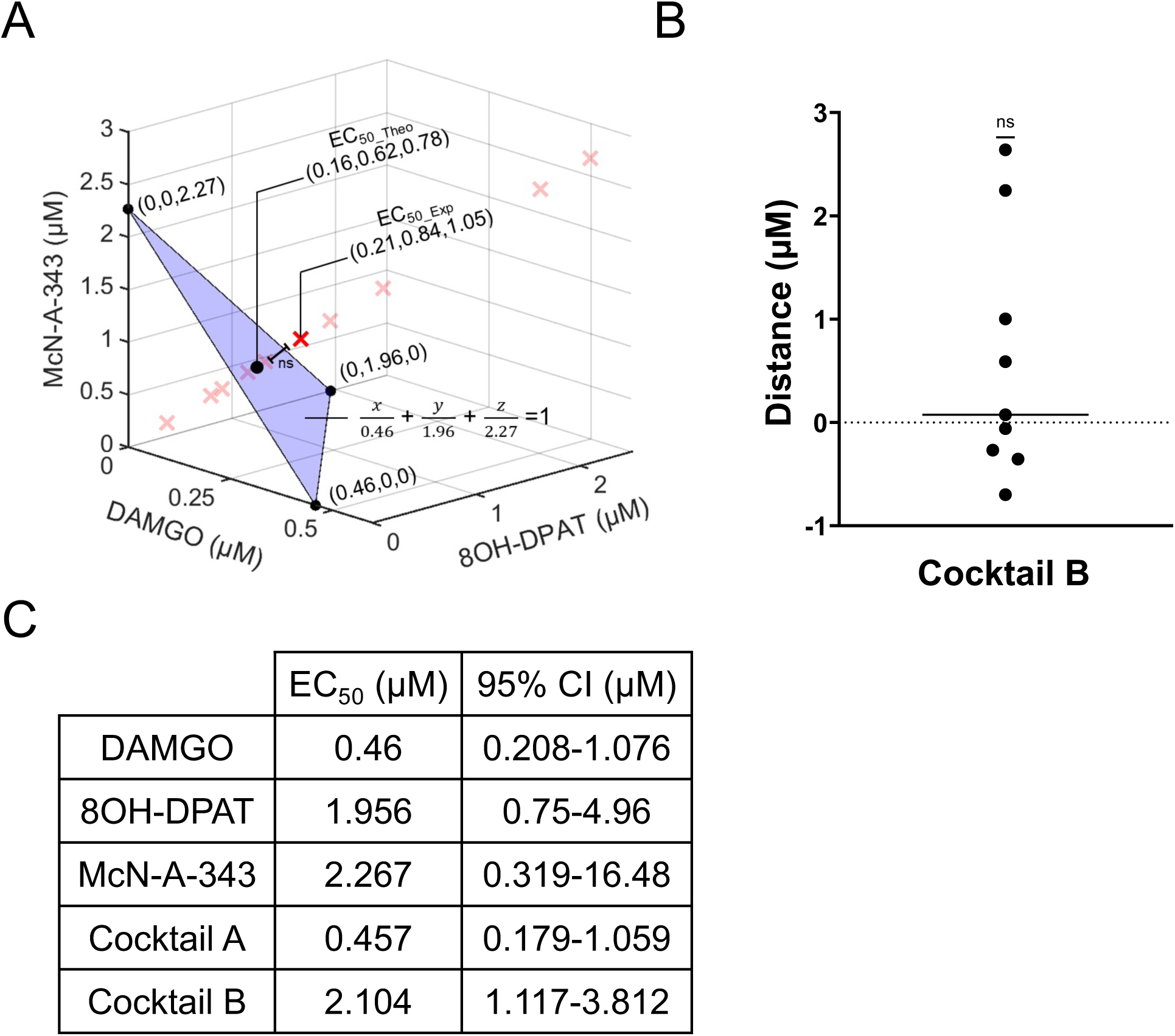
Detailed information and control experiments for pharmacologically induced antinociception. (A) A three-dimensional isobologram showing an additive interaction within Cocktail B, the theoretical EC_50_ of Cocktail B, EC_50_s of DAMGO, 8OH-DPAT and McN-A-343 are indicated by black dots, the overall and individual experimental EC_50_s of Cocktail B are indicated by solid and pale red cross and the blue plane denotes the additive isobole. (B) A plot demonstrating the additive interaction within Cocktail B with no significant difference between the theoretical and experimental EC_50_s. One sample t-test, t = 1.464, ns = not significant. (C) A table summarizes the EC_50_ and 95% confidence interval across pharmacological approaches.

**Figure S15.**
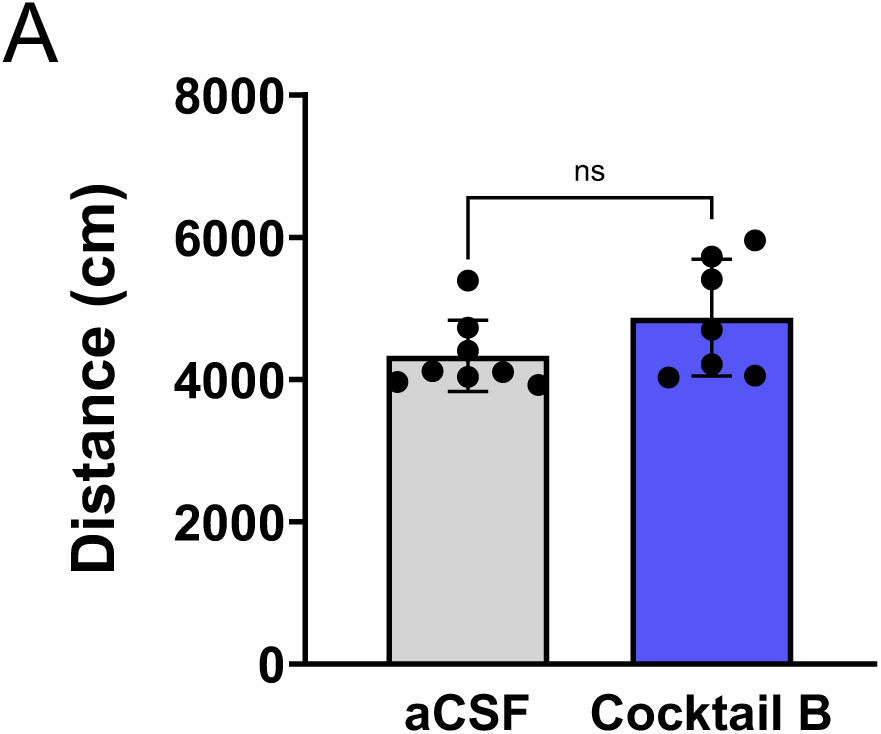
Effect of intra-LC infusion of Cocktail B on locomotion. (A) A plot showing the travel distance during open-field test upon the LC local infusions of either vehicle or Cocktail B. Mann-Whitney test, U = 16, ns = not significant. All of data are represented as mean ± SD.

**Figure. S16.**
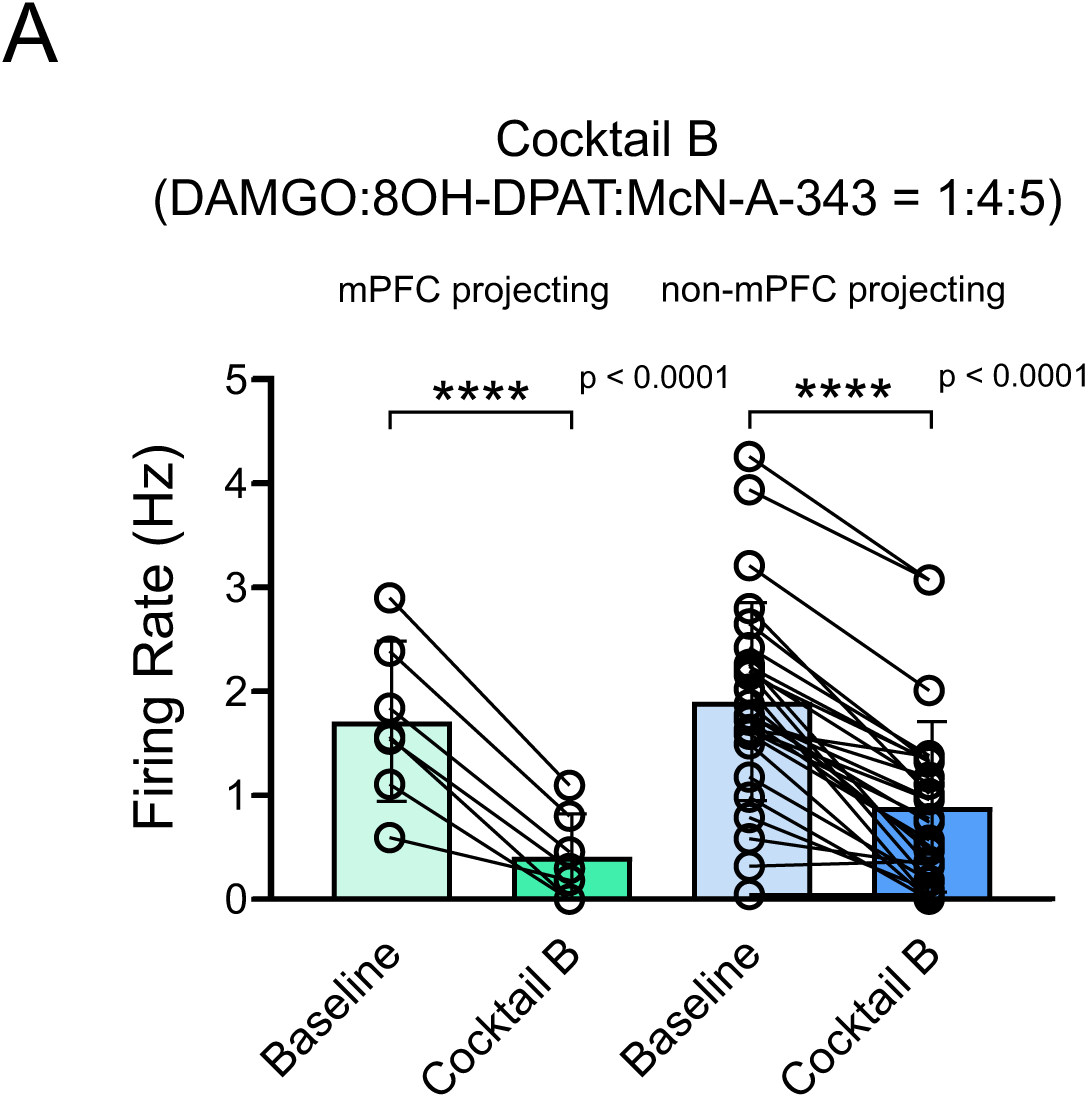
Effects of Cocktail B application on firing rate across LC modular organization. (A) A plot showing pharmacological effects on firing rate between LC modules under an administration of Cocktail B. Repeated measures two-way ANOVA, F = 102.3 (Pharmacology); 0.1369 (Modular); 0.4091 (interaction); 4.518 (ROI). ****p<0.0001. All of data are represented as mean ± SD.

**Figure S17.**
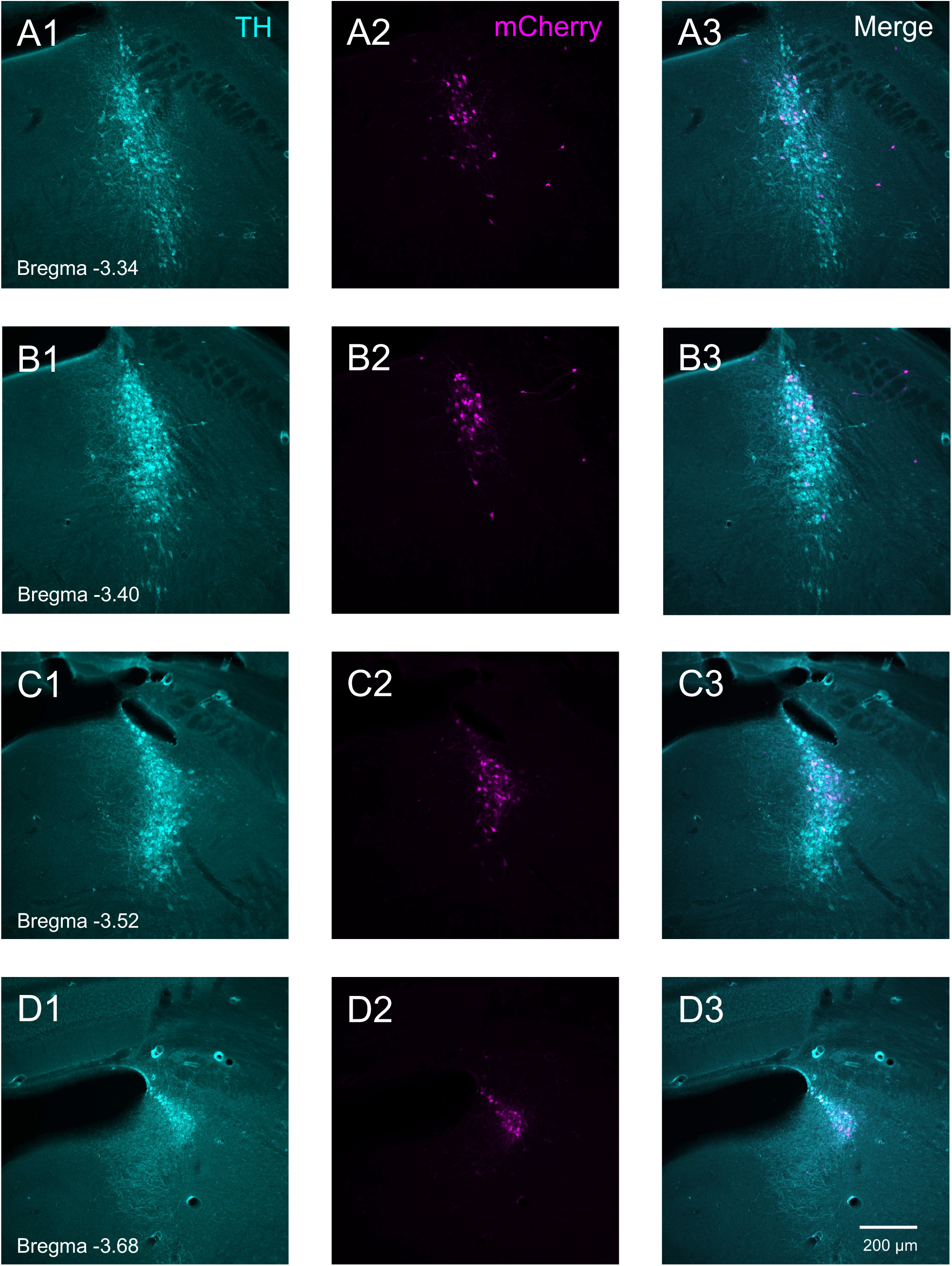
Distribution of mPFC-projecting LC-NE neurons revealed by retrograde virus. (A-D) Fluorescent images displaying the distribution of mPFC-innervating LC-NE neurons with colocalization of TH and mCherry immunoreactive signals indicated by cyan (TH) and magenta (mCherry) colors. Note the strikingly similarity between tracing results from retrograde tracer and virus (**Fig. S8**).

**Figure S18.**
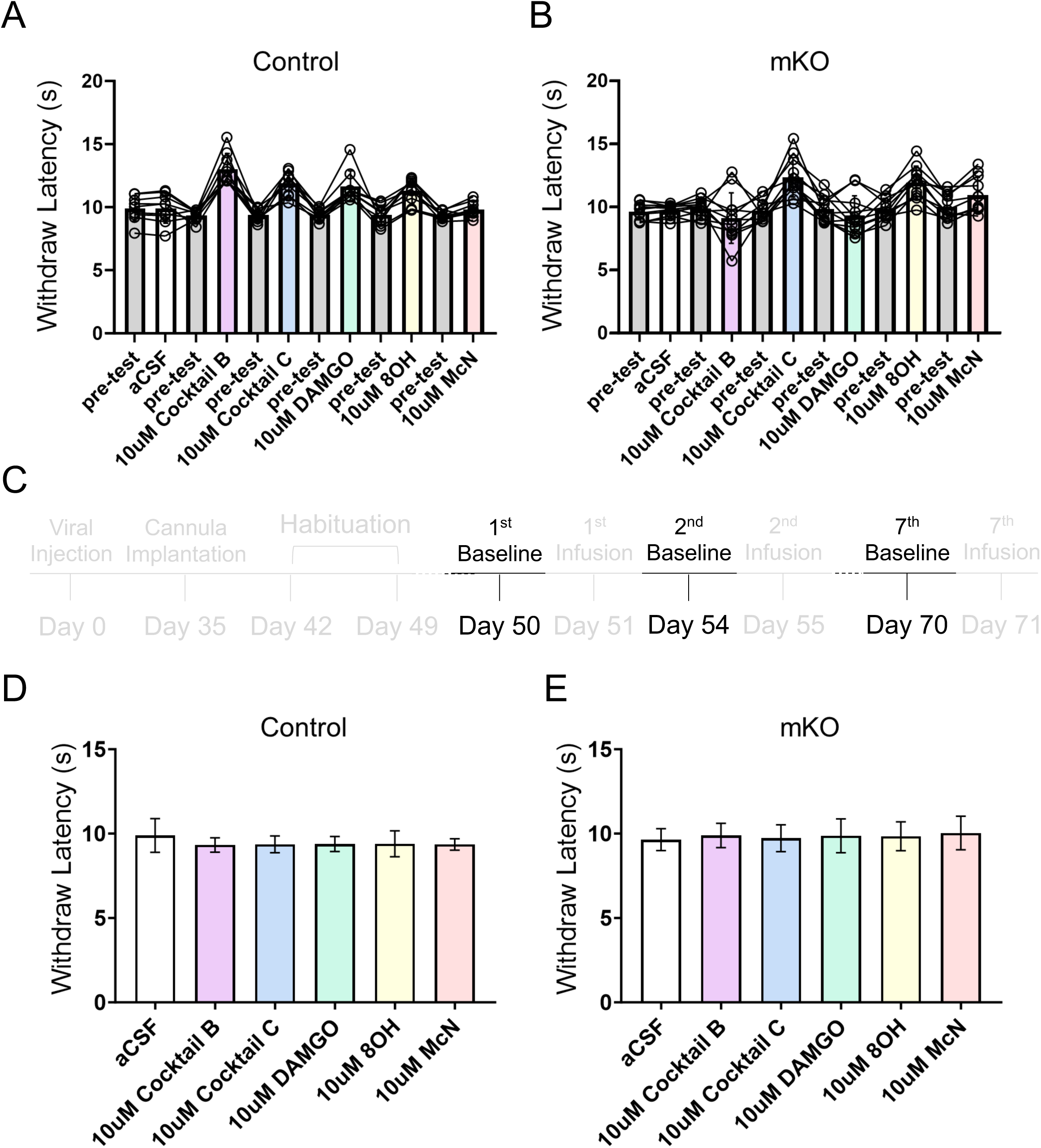
Hargreaves assay in concert with pharmacological infusions in mice with modular MOR knock out. (A-B) Plots demonstrating the raw data of behavioral examinations, baseline for each pharmacological dosage is collected from pre-test measurements. (C) A timeline showing the experimental design, only baselines are used in D and E. (D-E) Plots showing the comparison across baseline measurements pooled from all of data shown in A and B. Repeated measures one-way ANOVA, F = 1.269 (Control); 1.023 (mKO). No significant difference is found. All of data are represented as mean ± SD.

**Figure S19.**
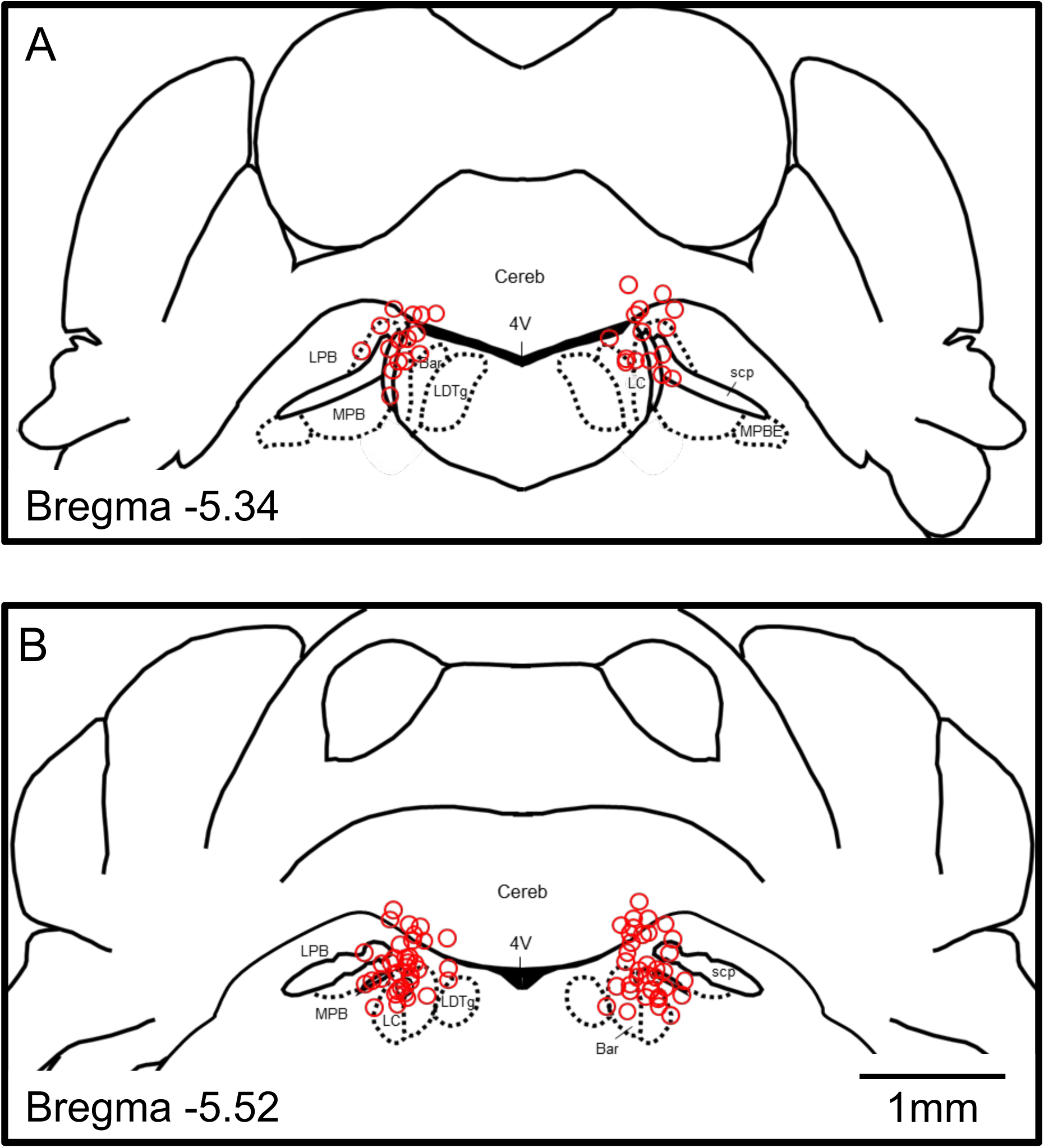
Cannula implantation site. (A-B) Superimposed brain maps show the reconstructed cannula implantation site (Red circles) from 51 mice included in this study. Abbreviation: 4V, fourth ventricle; Bar: Barrington’s nucleus; Cereb: cerebellum; LdTg: lateral dorsal tegmental area; LPB, lateral parabrachial nucleus; MPB, medial parabrachial nucleus; MPBE: medial parabrachial nucleus, external part; scp superior cerebellar peduncle.

## References

1. Armbruster, B. N., Li, X., Pausch, M. H., Herlitze, S. & Roth, B. L. Evolving the lock to fit the key to create a family of G protein-coupled receptors potently activated by an inert ligand. Proc Natl Acad Sci U S A 104, 5163–5168 (2007).

2. Roth, B. L. DREADDs for Neuroscientists. Neuron 89, 683–694 (2016).

3. Kim, C. K., Adhikari, A. & Deisseroth, K. Integration of optogenetics with complementary methodologies in systems neuroscience. Nat Rev Neurosci 18, 222–235 (2017).

4. Deisseroth, K. Optogenetics: 10 years of microbial opsins in neuroscience. Nat Neurosci 18, 1213–1225 (2015).

5. Tervo, D. G. R. et al. A Designer AAV Variant Permits Efficient Retrograde Access to Projection Neurons. Neuron 92, 372–382 (2016).

6. DeNardo, L. A. et al. Temporal evolution of cortical ensembles promoting remote memory retrieval. Nat Neurosci 22, 460–469 (2019).

7. Bhuiyan, S. A. et al. Harmonized cross-species cell atlases of trigeminal and dorsal root ganglia. 2023.07.04.547740 Preprint at 10.1101/2023.07.04.547740 (2023).

8. Liu, X. et al. Optogenetic stimulation of a hippocampal engram activates fear memory recall. Nature 484, 381–385 (2012).

9. Bansal, A., Shikha, S. & Zhang, Y. Towards translational optogenetics. Nat. Biomed. Eng 7, 349–369 (2022).

10. Song, J., Patel, R. V., Sharif, M., Ashokan, A. & Michaelides, M. Chemogenetics as a neuromodulatory approach to treating neuropsychiatric diseases and disorders. Molecular Therapy 30, 990–1005 (2022).

11. Walker, M. C. & Kullmann, D. M. Optogenetic and chemogenetic therapies for epilepsy. Neuropharmacology 168, 107751 (2020).

12. Bhokisham, N. et al. CRISPR-Cas System: The Current and Emerging Translational Landscape. Cells 12, 1103 (2023).

13. Berridge, C. W. & Foote, S. L. Enhancement of Behavioral and Electroencephalographic Indices of Waking following Stimulation of Noradrenergic β-Receptors within the Medial Septal Region of the Basal Forebrain. J Neurosci 16, 6999– 7009 (1996).

14. Varazzani, C., San-Galli, A., Gilardeau, S. & Bouret, S. Noradrenaline and Dopamine Neurons in the Reward/Effort Trade-Off: A Direct Electrophysiological Comparison in Behaving Monkeys. J Neurosci 35, 7866–7877 (2015).

15. Aston-Jones, G. & Cohen, J. D. AN INTEGRATIVE THEORY OF LOCUS COERULEUS-NOREPINEPHRINE FUNCTION: Adaptive Gain and Optimal Performance. Annual Review of Neuroscience 28, 403–450 (2005).

16. Kempadoo, K. A., Mosharov, E. V., Choi, S. J., Sulzer, D. & Kandel, E. R. Dopamine release from the locus coeruleus to the dorsal hippocampus promotes spatial learning and memory. Proc Natl Acad Sci U S A 113, 14835–14840 (2016).

17. Llorca-Torralba, M., Borges, G., Neto, F., Mico, J. A. & Berrocoso, E. Noradrenergic Locus Coeruleus pathways in pain modulation. Neuroscience 338, 93–113 (2016).

18. Sara, S. J. & Segal, M. Plasticity of sensory responses of locus coeruleus neurons in the behaving rat: implications for cognition. in Progress in Brain Research (eds. Barnes, C. D. & Pompeiano, O.) vol. 88 571–585 (Elsevier, 1991).

19. Mulvey, B. et al. Molecular and Functional Sex Differences of Noradrenergic Neurons in the Mouse Locus Coeruleus. Cell Reports 23, 2225–2235 (2018).

20. Zitnik, G. A. Control of arousal through neuropeptide afferents of the locus coeruleus. Brain Research 1641, 338–350 (2016).

21. Olpe, H.-R. & Steinmann, M. Responses of locus coeruleus neurons to neuropeptides. in Progress in Brain Research (eds. Barnes, C. D. & Pompeiano, O.) vol. 88 241–248 (Elsevier, 1991).

22. Weber, L. M. et al. The gene expression landscape of the human locus coeruleus revealed by single-nucleus and spatially-resolved transcriptomics. eLife 12, RP84628 (2024).

23. Guajardo, H. M., Snyder, K., Ho, A. & Valentino, R. J. Sex Differences in μ-Opioid Receptor Regulation of the Rat Locus Coeruleus and Their Cognitive Consequences. Neuropsychopharmacol 42, 1295–1304 (2017).

24. McCall, J. G. et al. CRH engagement of the locus coeruleus noradrenergic system mediates stress-induced anxiety. Neuron 87, 605–620 (2015).

25. Walling, S. G., Nutt, D. J., Lalies, M. D. & Harley, C. W. Orexin-A Infusion in the Locus Ceruleus Triggers Norepinephrine (NE) Release and NE-Induced Long-Term Potentiation in the Dentate Gyrus. J. Neurosci. 24, 7421–7426 (2004).

26. Schwarz, L. A. et al. Viral-genetic tracing of the input–output organization of a central noradrenaline circuit. Nature 524, 88–92 (2015).

27. Uematsu, A. et al. Modular organization of the brainstem noradrenaline system coordinates opposing learning states. Nat Neurosci 20, 1602–1611 (2017).

28. Hirschberg, S., Li, Y., Randall, A., Kremer, E. J. & Pickering, A. E. Functional dichotomy in spinal- vs prefrontal-projecting locus coeruleus modules splits descending noradrenergic analgesia from ascending aversion and anxiety in rats. eLife 6, e29808 (2017).

29. Chandler, D. J. et al. Redefining Noradrenergic Neuromodulation of Behavior: Impacts of a Modular Locus Coeruleus Architecture. J. Neurosci. 39, 8239–8249 (2019).

30. Llorca-Torralba, M. et al. Chemogenetic Silencing of the Locus Coeruleus-Basolateral Amygdala Pathway Abolishes Pain-Induced Anxiety and Enhanced Aversive Learning in Rats. Biol Psychiatry 85, 1021–1035 (2019).

31. Hnasko, T. S., Hjelmstad, G. O., Fields, H. L. & Edwards, R. H. Ventral Tegmental Area Glutamate Neurons: Electrophysiological Properties and Projections. J Neurosci 32, 15076–15085 (2012).

32. Morales, M. & Margolis, E. B. Ventral tegmental area: cellular heterogeneity, connectivity and behaviour. Nat Rev Neurosci 18, 73–85 (2017).

33. Ma, S., Zhong, H., Liu, X. & Wang, L. Spatial Distribution of Neurons Expressing Single, Double, and Triple Molecular Characteristics of Glutamatergic, Dopaminergic, or GABAergic Neurons in the Mouse Ventral Tegmental Area. J Mol Neurosci 73, 345–362 (2023).

34. Rodenkirch, C., Liu, Y., Schriver, B. J. & Wang, Q. Locus coeruleus activation enhances thalamic feature selectivity via norepinephrine regulation of intrathalamic circuit dynamics. Nat Neurosci 22, 120–133 (2019).

35. Hung, W.-C. et al. GABAB receptor-mediated tonic inhibition of locus coeruleus neurons plays a role in deep anesthesia induced by isoflurane. NeuroReport 31, 557 (2020).

36. Baulmann, J., Spitznagel, H., Herdegen, T., Unger, T. & Culman, J. Tachykinin receptor inhibition and c-Fos expression in the rat brain following formalin-induced pain. Neuroscience 95, 813–820 (1999).

37. Imbe, H. et al. Activation of ERK in the locus coeruleus following acute noxious stimulation. Brain Research 1263, 50–57 (2009).

38. Brightwell, J. J. & Taylor, B. K. Noradrenergic Neurons in the Locus Coeruleus Contribute to Neuropathic Pain. Neuroscience 160, 174–185 (2009).

39. Jongeling, A. C., Johns, M. E., Murphy, A. Z. & Hammond, D. L. Persistent inflammatory pain decreases the antinociceptive effects of the mu opioid receptor agonist DAMGO in the locus coeruleus of male rats. Neuropharmacology 56, 1017–1026 (2009).

40. Guo, T.-Z., Jiang, J.-Y., Buttermann, A. E. & Maze, M. Dexmedetomidine Injection into the Locus Ceruleus Produces Antinociception. Anesthesiology 84, 873–881. (1996).

41. Llorca-Torralba, M., Mico, J. A. & Berrocoso, E. Behavioral effects of combined morphine and MK-801 administration to the locus coeruleus of a rat neuropathic pain model. Progress in Neuro-Psychopharmacology and Biological Psychiatry 84, 257–266 (2018).

42. Norris, M. R. et al. Endogenous opioids gate the locus coeruleus pain generator. 2023.10.20.562785 Preprint at 10.1101/2023.10.20.562785 (2023).

43. Suárez-Pereira, I. et al. The Role of the Locus Coeruleus in Pain and Associated Stress-Related Disorders. Biological Psychiatry 91, 786–797 (2022).

44. Bahari, Z. & Meftahi, G. H. Spinal α2-adrenoceptors and neuropathic pain modulation; therapeutic target. Br J Pharmacol 176, 2366–2381 (2019).

45. Llorca-Torralba, M. et al. Pain and depression comorbidity causes asymmetric plasticity in the locus coeruleus neurons. Brain 145, 154–167 (2022).

46. Gu, X. et al. Neurons in the caudal ventrolateral medulla mediate descending pain control. Nat Neurosci 26, 594–605 (2023).

47. Li, Y. et al. Retrograde optogenetic characterization of the pontospinal module of the locus coeruleus with a canine adenoviral vector. Brain Res 1641, 274–290 (2016).

48. Hickey, L. et al. Optoactivation of locus ceruleus neurons evokes bidirectional changes in thermal nociception in rats. J. Neurosci. 34, 4148–4160 (2014).

49. Kucharczyk, M. W., Di Domenico, F. & Bannister, K. Distinct brainstem to spinal cord noradrenergic pathways inversely regulate spinal neuronal activity. Brain 145, 2293–2300 (2022).

50. Kucharczyk, M. W., Di Domenico, F. & Bannister, K. A critical brainstem relay for mediation of diffuse noxious inhibitory controls. Brain 146, 2259–2267 (2023).

51. Camarena-Delgado, C. et al. Nerve injury induces transient locus coeruleus activation over time: role of the locus coeruleus–dorsal reticular nucleus pathway. PAIN 163, 943 (2022).

52. Lubejko, S. T. et al. Inputs to the locus coeruleus from the periaqueductal gray and rostroventral medulla shape opioid-mediated descending pain modulation. Science Advances 10, eadj9581 (2024).

53. Bouret, S. & Sara, S. J. Reward expectation, orientation of attention and locus coeruleus-medial frontal cortex interplay during learning. Eur J Neurosci 20, 791–802 (2004).

54. Watanabe, M., Uematsu, A. & Johansen, J. P. Bidirectional emotional regulation through prefrontal innervation of the locus coeruleus. 2024.02.07.579259 Preprint at 10.1101/2024.02.07.579259 (2024).

55. Zhang, Y. et al. Fast and sensitive GCaMP calcium indicators for imaging neural populations. Nature 615, 884–891 (2023).

56. McCall, J. G. et al. Locus coeruleus to basolateral amygdala noradrenergic projections promote anxiety-like behavior. eLife 6, e18247 (2017).

57. Wu, R.-N., Kuo, C.-C., Min, M.-Y., Chen, R.-F. & Yang, H.-W. Extracellular Signal-Regulated Kinases Mediate an Autoregulation of GABAB-Receptor-Activated Whole-Cell Current in Locus Coeruleus Neurons. Sci Rep 10, 7869 (2020).

58. Ishimatsu, M. & Williams, J. T. Synchronous Activity in Locus Coeruleus Results from Dendritic Interactions in Pericoerulear Regions. J Neurosci 16, 5196–5204 (1996).

59. Rupprecht, P. et al. A database and deep learning toolbox for noise-optimized, generalized spike inference from calcium imaging. Nat Neurosci 24, 1324–1337 (2021).

60. Rupprecht, P., Fan, W., Sullivan, S. J., Helmchen, F. & Sdrulla, A. D. Spike inference from mouse spinal cord calcium imaging data. 2024.07.17.603957 Preprint at 10.1101/2024.07.17.603957 (2024).

61. Totah, N. K., Neves, R. M., Panzeri, S., Logothetis, N. K. & Eschenko, O. The Locus Coeruleus Is a Complex and Differentiated Neuromodulatory System. Neuron 99, 1055–1068.e6 (2018).

62. McKinney, A. et al. Cellular composition and circuit organization of the locus coeruleus of adult mice. eLife 12, e80100 (2023).

63. Williams, J. T., Henderson, G. & North, R. A. Characterization of α2-adrenoceptors which increase potassium conductance in rat locus coeruleus neurones. Neuroscience 14, 95–101 (1985).

64. Arima, J., Kubo, C., Ishibashi, H. & Akaike, N. α2-Adrenoceptor-mediated potassium currents in acutely dissociated rat locus coeruleus neurones. J Physiol 508, 57–66 (1998).

65. Davy, O. et al. Noradrenergic cross-modular reciprocal inhibition within the locus coeruleus. 2022.09.07.506929 Preprint at 10.1101/2022.09.07.506929 (2022).

66. Kuo, C.-C. et al. Carbachol increases locus coeruleus activation by targeting noradrenergic neurons, inhibitory interneurons and inhibitory synaptic transmission. European Journal of Neuroscience 57, 32–53 (2023).

67. Pan, Y.-Z., Li, D.-P., Chen, S.-R. & Pan, H.-L. Activation of μ-opioid receptors excites a population of locus coeruleus-spinal neurons through presynaptic disinhibition. Brain Research 997, 67–78 (2004).

68. Harris, G. & Williams, J. Transient homologous mu-opioid receptor desensitization in rat locus coeruleus neurons. J Neurosci 11, 2574–2581 (1991).

69. Wagner-Altendorf, T. A., Fischer, B. & Roeper, J. Axonal projection-specific differences in somatodendritic α2 autoreceptor function in locus coeruleus neurons. European Journal of Neuroscience 50, 3772–3785 (2019).

70. Al-Hasani, R. & Bruchas, M. R. Molecular Mechanisms of Opioid Receptor-Dependent Signaling and Behavior. Anesthesiology 115, 1363–1381 (2011).

71. Che, T. & Roth, B. L. Molecular basis of opioid receptor signaling. Cell 186, 5203–5219 (2023).

72. Luyo, Z. N. M., Lawrence, A. B., Stathopoulos, T. G. & Mitrano, D. A. Localization and neurochemical identity of alpha1-adrenergic receptor-containing elements in the mouse locus coeruleus. J Chem Neuroanat 133, 102343 (2023).

73. Seo, D.-O. et al. A locus coeruleus to dentate gyrus noradrenergic circuit modulates aversive contextual processing. Neuron 109, 2116–2130.e6 (2021).

74. Huang, R. et al. Isobologram Analysis: A Comprehensive Review of Methodology and Current Research. Front Pharmacol 10, 1222 (2019).

75. Hwang, D. Y., Carlezon, W. A., Isacson, O. & Kim, K. S. A high-efficiency synthetic promoter that drives transgene expression selectively in noradrenergic neurons. Hum Gene Ther 12, 1731–1740 (2001).

76. Soudais, C., Laplace-Builhe, C., Kissa, K. & Kremer, E. J. Preferential transduction of neurons by canine adenovirus vectors and their efficient retrograde transport in vivo. FASEB J 15, 2283–2285 (2001).

77. Weibel, R. et al. Mu Opioid Receptors on Primary Afferent Nav1.8 Neurons Contribute to Opiate-Induced Analgesia: Insight from Conditional Knockout Mice. PLoS One 8, e74706 (2013).

78. Yorgason, J. T., Zeppenfeld, D. M. & Williams, J. T. Cholinergic Interneurons Underlie Spontaneous Dopamine Release in Nucleus Accumbens. J Neurosci 37, 2086–2096 (2017).

79. Warwick, C. et al. Cell type-specific calcium imaging of central sensitization in mouse dorsal horn. Nat Commun 13, 5199 (2022).

80. Boal, A. M., McGrady, N. R., Risner, M. L. & Calkins, D. J. Sensitivity to extracellular potassium underlies type-intrinsic differences in retinal ganglion cell excitability. Front Cell Neurosci 16, 966425 (2022).

81. Schmitz, G. P. et al. Psychedelic compounds directly excite 5-HT2A Layer 5 Pyramidal Neurons in the Prefrontal Cortex through a 5-HT2A Gq -mediated activation mechanism. 2022.11.15.516655 Preprint at 10.1101/2022.11.15.516655 (2022).

82. Padawer-Curry, J. A. et al. Psychedelic 5-HT2A receptor agonism: neuronal signatures and altered neurovascular coupling. 2023.09.23.559145 Preprint at 10.1101/2023.09.23.559145 (2023).

83. Hsu, C.-L., Yang, H.-W., Yen, C.-T. & Min, M.-Y. Comparison of synaptic transmission and plasticity between sensory and cortical synapses on relay neurons in the ventrobasal nucleus of the rat thalamus. J Physiol 588, 4347–4363 (2010).

84. Yi, G. & Grill, W. M. Average firing rate rather than temporal pattern determines metabolic cost of activity in thalamocortical relay neurons. Sci Rep 9, 6940 (2019).

85. Markram, H. et al. Interneurons of the neocortical inhibitory system. Nat Rev Neurosci 5, 793–807 (2004).

86. Bardoni, R. et al. Pain Inhibits GRPR Neurons via GABAergic Signaling in the Spinal Cord. Sci Rep 9, 15804 (2019).

87. Wang, H.-Y. et al. GABAB receptor-mediated tonic inhibition regulates the spontaneous firing of locus coeruleus neurons in developing rats and in citalopram-treated rats. J Physiol 593, 161–180 (2015).

88. Torrecilla, M., Quillinan, N., Williams, J. T. & Wickman, K. Pre- and postsynaptic regulation of locus coeruleus neurons after chronic morphine treatment: a study of GIRK knockout mice. Eur J Neurosci 28, 618–624 (2008).

89. Roth, S., Kholodenko, B. N., Smit, M. J. & Bruggeman, F. J. G Protein–Coupled Receptor Signaling Networks from a Systems Perspective. Mol Pharmacol 88, 604–616 (2015).

90. Jiang, H., Galtes, D., Wang, J. & Rockman, H. A. G protein-coupled receptor signaling: transducers and effectors. Am J Physiol Cell Physiol 323, C731–C748 (2022).

91. Fiorillo, C. & Williams, J. Opioid desensitization: interactions with G-protein-coupled receptors in the locus coeruleus. J Neurosci 16, 1479–1485 (1996).

92. Li, M.-J. et al. Cholinergic and glutamatergic transmission at synapses between pedunculopotine tegmental nucleus axonal terminals and A7 catecholamine cell group noradrenergic neurons in the rat. Neuropharmacology 110, 237–250 (2016).

93. Borges, G. P., Micó, J. A., Neto, F. L. & Berrocoso, E. Corticotropin-Releasing Factor Mediates Pain-Induced Anxiety through the ERK1/2 Signaling Cascade in Locus Coeruleus Neurons. Int J Neuropsychopharmacol 18, pyv019 (2015).

94. Lane-Ladd, S. B. et al. CREB (cAMP Response Element-Binding Protein) in the Locus Coeruleus: Biochemical, Physiological, and Behavioral Evidence for a Role in Opiate Dependence. J Neurosci 17, 7890–7901 (1997).

95. Moreno, E. et al. Pharmacological targeting of G protein-coupled receptor heteromers. Pharmacol Res 185, 106476 (2022).

96. Gomes, I., Ijzerman, A. P., Ye, K., Maillet, E. L. & Devi, L. A. G protein-coupled receptor heteromerization: a role in allosteric modulation of ligand binding. Mol. Pharmacol 79, 1044–1052 (2011).

97. De Oliveira, P. A. et al. Preferential Gs protein coupling of the galanin Gal1 receptor in the µ-opioid-Gal1 receptor heterotetramer. Pharmacol Res 182, 106322 (2022).

98. Levinstein, M. R. et al. Unique pharmacodynamic properties and low abuse liability of the µ-opioid receptor ligand (S)-methadone. Mol Psychiatry 29, 624–632 (2024).

99. Wong, T.-S. et al. G protein-coupled receptors in neurodegenerative diseases and psychiatric disorders. Sig Transduct Target Ther 8, 1–57 (2023).

100. Copits, B. A., Pullen, M. Y. & Gereau, R. W. Spotlight on pain: optogenetic approaches for interrogating somatosensory circuits. Pain 157, 2424–2433 (2016).

101. Fenno, L., Yizhar, O. & Deisseroth, K. The Development and Application of Optogenetics. Annu Rev Neurosci 34, 389–412 (2011).

102. Magnus, C. J. et al. Ultrapotent chemogenetics for research and potential clinical applications. Science 364, eaav5282 (2019).

103. Sternson, S. M. & Roth, B. L. Chemogenetic Tools to Interrogate Brain Functions. Annu. Rev. Neurosci. 37, 387–407 (2014).

104. Copits, B. A. et al. A photoswitchable GPCR-based opsin for presynaptic inhibition. Neuron 109, 1791–1809.e11 (2021).

105. Siuda, E. R. et al. Spatiotemporal control of opioid signaling and behavior. Neuron 86, 923–935 (2015).

106. Airan, R. D., Thompson, K. R., Fenno, L. E., Bernstein, H. & Deisseroth, K. Temporally precise in vivo control of intracellular signalling. Nature 458, 1025–1029 (2009).

107. Mahn, M., Prigge, M., Ron, S., Levy, R. & Yizhar, O. Biophysical constraints of optogenetic inhibition at presynaptic terminals. Nat Neurosci 19, 554–556 (2016).

108. Raimondo, J. V., Kay, L., Ellender, T. J. & Akerman, C. J. Optogenetic silencing strategies differ in their effects on inhibitory synaptic transmission. Nat Neurosci 15, 1102– 1104 (2012).

109. Galvan, A. et al. Ultrastructural localization of DREADDs in monkeys. Eur J Neurosci 50, 2801–2813 (2019).

110. Luskin, A. T. et al. A diverse network of pericoerulear neurons control arousal states. 2022.06.30.498327 Preprint at 10.1101/2022.06.30.498327 (2023).

111. Barcomb, K., Olah, S. S., Kennedy, M. J. & Ford, C. P. Properties and modulation of excitatory inputs to the locus coeruleus. J Physiol 600, 4897–4916 (2022).

112. Breton-Provencher, V. & Sur, M. Active control of arousal by a locus coeruleus GABAergic circuit. Nat Neurosci 22, 218–228 (2019).

113. Kuo, C.-C. et al. Inhibitory interneurons regulate phasic activity of noradrenergic neurons in the mouse locus coeruleus and functional implications. J Physiol 598, 4003–4029 (2020).

114. Martinez-Lage, M., Puig-Serra, P., Menendez, P., Torres-Ruiz, R. & Rodriguez-Perales, S. CRISPR/Cas9 for Cancer Therapy: Hopes and Challenges. Biomedicines 6, 105 (2018).

115. Botterill, J. J. et al. Off-Target Expression of Cre-Dependent Adeno-Associated Viruses in Wild-Type C57BL/6J Mice. eNeuro 8, ENEURO.0363-21.2021 (2021).

116. Fischer, K. B., Collins, H. K. & Callaway, E. M. Sources of off-target expression from recombinase-dependent AAV vectors and mitigation with cross-over insensitive ATG-out vectors. Proc Natl Acad Sci U S A 116, 27001–27010 (2019).

117. Beverly, A., Kaye, A. D., Ljungqvist, O. & Urman, R. D. Essential Elements of Multimodal Analgesia in Enhanced Recovery After Surgery (ERAS) Guidelines. Anesthesiology Clinics 35, e115–e143 (2017).

118. Chou, R. et al. Management of Postoperative Pain: A Clinical Practice Guideline From the American Pain Society, the American Society of Regional Anesthesia and Pain Medicine, and the American Society of Anesthesiologists’ Committee on Regional Anesthesia, Executive Committee, and Administrative Council. The Journal of Pain 17, 131–157 (2016).

119. Gilron, I., Jensen, T. S. & Dickenson, A. H. Combination pharmacotherapy for management of chronic pain: from bench to bedside. Lancet Neurol 12, 1084–1095 (2013).

120. Helander, E. M. et al. Multimodal Analgesia, Current Concepts, and Acute Pain Considerations. Curr Pain Headache Rep 21, 3 (2017).

121. Freo, U. Paracetamol for Multimodal Analgesia. Pain Management 12, 737–750 (2022).

122. Chen, T., Wang, J., Wang, Y.-Q. & Chu, Y.-X. Current Understanding of the Neural Circuitry in the Comorbidity of Chronic Pain and Anxiety. Neural Plast 2022, 4217593 (2022).

123. Zis, P. et al. Depression and chronic pain in the elderly: links and management challenges. Clin Interv Aging 12, 709–720 (2017).

124. Haack, M., Simpson, N., Sethna, N., Kaur, S. & Mullington, J. Sleep deficiency and chronic pain: potential underlying mechanisms and clinical implications. Neuropsychopharmacology 45, 205–216 (2020).

125. Donertas-Ayaz, B. & Caudle, R. M. Locus coeruleus-noradrenergic modulation of trigeminal pain: Implications for trigeminal neuralgia and psychiatric comorbidities. Neurobiol Pain 13, 100124 (2023).

126. Llorca-Torralba, M. et al. Opioid Activity in the Locus Coeruleus Is Modulated by Chronic Neuropathic Pain. Mol Neurobiol 56, 4135–4150 (2019).

127. Llorca-Torralba, M., Pilar-Cuéllar, F., da Silva Borges, G., Mico, J. A. & Berrocoso, E. Opioid receptors mRNAs expression and opioids agonist-dependent G-protein activation in the rat brain following neuropathy. Progress in Neuro-Psychopharmacology and Biological Psychiatry 99, 109857 (2020).

128. 128. Schindelin, J., et al. Fiji: an open-source platform for biological-image analysis. Nat Methods 9, 676–682 (2012).

129. Pnevmatikakis, E. A. & Giovannucci, A. NoRMCorre: An online algorithm for piecewise rigid motion correction of calcium imaging data. Journal of Neuroscience Methods 291, 83– 94 (2017).

130. Zhou, P. et al. Efficient and accurate extraction of in vivo calcium signals from microendoscopic video data. eLife 7, e28728.

131. Tallarida, R. J., Porreca, F. & Cowan, A. Statistical analysis of drug-drug and site-site interactions with isobolograms. Life Sciences 45, 947–961 (1989).

132. Nasehi, M., Hajikhani, M., Ebrahimi-Ghiri, M. & Zarrindast, M.-R. Interaction between NMDA and CB2 function in the dorsal hippocampus on memory consolidation impairment: an isobologram analysis. Psychopharmacology (Berl) 234, 507–514 (2017).

