## Supplemental Code for "Neural circuit-selective, multiplexed pharmacological targeting of prefrontal cortex-projecting locus coeruleus neurons drives antinociception": binary_spike_location.m

```
function [events]=binary_spike_location(data_in,spikes_loc)
% events: Binary form of spike location
% data_in: traces for prediction, default 'dF_traces'
% spikes_loc: spikes_locations, default 'spike_cascade'
% Example code: [events]=binary_spike_location(dF_traces,spikes_cascade);
events=zeros(size(data_in));
for roi=1:size(events,1)
    temp_spike=spikes_loc{roi};
    for time_point=1:size(temp_spike,1)
        events(roi,temp_spike(time_point,1))=1;
    end
end
events=events';
end
```
