## Supplemental Code for "Neural circuit-selective, multiplexed pharmacological targeting of prefrontal cortex-projecting locus coeruleus neurons drives antinociception": pharmacology_profiling.m

```
function [accept_ROI,pharmacology_results,spikes_bin]=pharmacology_profiling(spikes_cascade,ROI_information,time_all,time_span,FPS,bin)
% Example code: [accept_ROI,pharmacology_results,spikes_bin]=pharmacology_profiling(spikes_cascade,ROI_information_sheet,3500,1750,20,0.5);
% accept_ROI: Accepted ROIs
% pharmacology_results: Results of pharmacological profiling (firing rate during baseline & drug + Ratio of FR) 
% spikes_bin: Binned spike firing rate
% spikes_cascade: Predicted spike location, default 'spikes_cascade'
% ROI_information: Main sheet for spike inference, default 'ROI_information_sheet'
% time_all: total time frames of trace, usually 3500
% time_span: time frames for either baseline or drug epoch, usually 1750
% FPS: Frame rate
% bin: Bin size (Sec)


spikes_cascade_temp={};
accept_ROI=[];
for h=1:size(spikes_cascade,2)
    if ROI_information.quality(h)==1 && ROI_information.repeated_cell(h)==0
        spikes_cascade_temp=cat(2,spikes_cascade_temp,spikes_cascade{h});
        accept_ROI=cat(1,accept_ROI,h);
    end
end

for i=1:size(spikes_cascade_temp,2)
    pre=length(find(spikes_cascade_temp{i}<=time_all/2 & spikes_cascade_temp{i}>time_all/2-time_span));
    post=length(find(spikes_cascade_temp{i}>time_all-time_span));
    pharmacology_results(i,:)=[pre post];
end

pharmacology_results=pharmacology_results/((time_all/2)/FPS);
pharmacology_results(:,3)=pharmacology_results(:,2)./pharmacology_results(:,1);

for j=1:size(spikes_cascade_temp,2)
    for k=1:floor(time_span/(bin*FPS))
        spikes_bin.pre(k,j)=length(find(spikes_cascade_temp{j}>bin*FPS*(k-1) & spikes_cascade_temp{j}<=bin*FPS*k));
        spikes_bin.post(k,j)=length(find(spikes_cascade_temp{j}>time_all/2+bin*FPS*(k-1) & spikes_cascade_temp{j}<=time_all/2+bin*FPS*k));
    end
end

for k=1:size(accept_ROI,1)
    run=accept_ROI(k);
    if ROI_information.manual(run,1)>0
        pharmacology_results(k,1)=ROI_information.manual(run,2)/((time_all/2)/FPS);
        pharmacology_results(k,3)=pharmacology_results(k,2)/pharmacology_results(k,1);
    end
    if ROI_information.manual(run,3)>0
        pharmacology_results(k,2)=ROI_information.manual(run,4)/((time_all/2)/FPS);
        pharmacology_results(k,3)=pharmacology_results(k,2)/pharmacology_results(k,1);
    end
end

figure;
b=bar(pharmacology_results(:,1:2),'hist');
b(1).FaceColor = [.3 .7 .6];
b(2).FaceColor = [.8 .3 .5];
set(b, {'DisplayName'}, {'Baseline','Drug Application'}');
legend();
set(gcf,'color','w');
ax=gca();
ax.FontSize=14;
title('Overview of Pharmacology Effects','fontsize', 18)
xlabel('#ROI')
ylabel('Firing Rate (Hz)')
xticks([1:size(pharmacology_results,1)])
xticklabels(string(accept_ROI)')
box off

figure;
b=bar(pharmacology_results(:,3));
hold on
b(1).FaceColor = [.4 .5 .7];
set(gcf,'color','w');
ax=gca();
ax.FontSize=14;
title('Overview of Pharmacology Effects','fontsize', 18)
xlabel('#ROI')
ylabel('Ratio')
xticks([1:size(pharmacology_results,1)])
xticklabels(string(accept_ROI)')
box off
plot(ones(size(pharmacology_results,1),1),'r')

end
```
