## Supplemental Code for "Neural circuit-selective, multiplexed pharmacological targeting of prefrontal cortex-projecting locus coeruleus neurons drives antinociception": prediction_material_convorter.m

```
function [dF_traces]=prediction_material_convorter(data_in)
% data_in: Temporal ROI info, default as results.C_raw
% dF_traces: Selective format % filename for spike prediction
% Example code: [dF_traces]=prediction_material_convorter(results.C_raw);
dF_traces=data_in';
for roi=1:size(dF_traces(1:1800,:),2)
    dF_traces(1:1800,roi)=dF_traces(1:1800,roi)-min(dF_traces(1:1800,roi));
    dF_traces(1:1800,roi)=dF_traces(1:1800,roi)/max(dF_traces(1:1800,roi));
    dF_traces(1:1800,roi)=dF_traces(1:1800,roi)/2;
end
for roi=1:size(dF_traces(1801:3600,:),2)
    dF_traces(1801:3600,roi)=dF_traces(1801:3600,roi)-min(dF_traces(1801:3600,roi));
    dF_traces(1801:3600,roi)=dF_traces(1801:3600,roi)/max(dF_traces(1801:3600,roi));
    dF_traces(1801:3600,roi)=dF_traces(1801:3600,roi)/2;
end
dF_traces=dF_traces';
end
```
