## Supplemental Code for "Neural circuit-selective, multiplexed pharmacological targeting of prefrontal cortex-projecting locus coeruleus neurons drives antinociception": roi_contours.m

```
function roi_contours(mean_image,ROI_spatial,side1,side2,ROI_sel,contour_cutoff)
%Example code: roi_contours(mean_image,results.A,128,128,0,[0.25,0.15]);
%ROI_spatial: ROI spatial info from CNMF algorithm, usually 'results.A'
%mean_image: Mean of Z projection for whole imaging data
%side1/2: Image size, usually 128
%ROI_sel: Interested ROIs, 0 for all or [1 2 3...]
%contour_cutoff: Contour borders, ex: [0.25,0.15];

mean_image=mat2gray(imread(mean_image));
all_roi=double(full(ROI_spatial));
all_roi=reshape(all_roi,[side2,side1,size(all_roi,2)]);
roi_location=all_roi;
all_roi=all_roi/max(all_roi,[],'all');
all_roi_boundary=all_roi;
all_roi(all_roi>=contour_cutoff(1))=1;
all_roi(all_roi<contour_cutoff(1))=0;
all_roi_boundary(all_roi_boundary>=contour_cutoff(1))=1;
all_roi_boundary(all_roi_boundary<contour_cutoff(2))=0;

roi_center=zeros(size(ROI_spatial,2),2);
for roi=1:size(ROI_spatial,2)
    roi_center(roi,1)=sum(sum(roi_location(:,:,roi),1).*[1:side1])/sum(sum(roi_location(:,:,roi)));
    roi_center(roi,2)=sum(sum(roi_location(:,:,roi),2).*[1:side2]')/sum(sum(roi_location(:,:,roi)));
end

roi_seq=string(1:1:size(all_roi,3));

figure;
imshow(mean_image);
hold on;
if ROI_sel==0
    for z=1:size(all_roi,3)
        contour(all_roi_boundary(:,:,z),'color',[0 0.4470 0.7410]);
        contour(all_roi(:,:,z),'color',[0 0.8 0.8]);
        contour_temp=all_roi(:,:,z);
        t=text(mean(find(sum(contour_temp,1)>0)),mean(find(sum(contour_temp,2)>0)),roi_seq(z));
        set(t,'Color','r');
        set(t,'Fontsize',10);
    end
end
set(gcf,'color','w');
hold off
end
```
