## Supplemental Code for "Neural circuit-selective, multiplexed pharmacological targeting of prefrontal cortex-projecting locus coeruleus neurons drives antinociception": ROI_cross_correlation.m

```
function [accept_ROI,repeat_roi,ROI_corrcoef_map]=ROI_cross_correlation(events,ROI_information,threshold)
% Example code: [accept_ROI,repeat_roi,ROI_corrcoef_map]=ROI_cross_correlation(events,ROI_information_sheet,0.3);
% accept_ROI: accepted ROIs
% repeat_roi: found repeated ROIs
% ROI_corrcoef_map: cross correlation matrix across ROIs
% events: binary traces of spike location, default 'events'
% ROI_information: Main sheet for spike inference, default 'ROI_information_sheet'
% threshold: threshold of correlation coefficient for idetifying repeated ROIs, default 0.3

accept_ROI=[];
for q=1:size(events,2)
    if ROI_information.quality(q)==0
        events(:,q)=NaN;
    end
    if ~isnan(events(1,q))
       accept_ROI=cat(1,accept_ROI,q);
    end
    %if ROI_information.manual(q,1)==1 || ROI_information.manual(q,3)==1
    %    events(:,q)=NaN;
    %end
end
events_temp=[];
for r=1:size(events,2)
    if ~isnan(events(1,r))
        events_temp=cat(2,events_temp,events(:,r));
    end
end

a=corrcoef(events_temp);
a(find(a==1))=NaN;
b=reshape(a,[size(a,1)*size(a,2),1])+1;
for i=1:80
    c(i)=length(find(b>(i-1)*0.025))/2-length(find(b>i*0.025))/2;
end

d=[1:80]*0.025-1;

ROI_corrcoef_map=a;

subplot 211; bar(d,c,'k');
title('Spectrum of Cross Correlation','fontsize', 14);
xlabel('Pearson Correlation Coefficients');
ylabel('Pair Counts');
xlim([-0.4 0.8])
set(gcf,'color','w');
ax=gca();
ax.FontSize=10;
subplot 212; bar(d,c,'k');
title('Repeated Cell Judgement','fontsize', 14);
xlabel('Pearson Correlation Coefficients');
ylabel('Pair Counts');
xlim([0 0.8])
ylim([0 4])
set(gcf,'color','w');
ax=gca();
ax.FontSize=10;

e=find(b>threshold+1);
repeat_roi=[];
for j=1:length(e)
    if mod(e(j),size(a,1))~=0
        repeat_roi(j,:)=[floor(e(j)/size(a,1))+1,mod(e(j),size(a,1))];
    else
        repeat_roi(j,:)=[floor(e(j)/size(a,1)),size(a,2)];
    end
end


for k=1:size(ROI_information.quality,1)
    for m=1:length(e)
        if ROI_information.quality(k,1)==0 && repeat_roi(m,1)>=k
            repeat_roi(m,1)=repeat_roi(m,1)+1;
        end
        if ROI_information.quality(k,1)==0 && repeat_roi(m,2)>=k
            repeat_roi(m,2)=repeat_roi(m,2)+1;
        end
    end
end

figure;
h=heatmap(ROI_corrcoef_map,'CellLabelColor','none');
colormap(jet);
caxis([-1,1])
label_temp=[1:size(ROI_information.quality,1)];
for n=1:size(ROI_information.quality,1)
    if ROI_information.quality(n,1)==0
        label_temp(n)=0;
    end
end
label_temp(find(label_temp==0))=[];
label_temp=string(label_temp);
CustomXLabels=label_temp;
CustomYLabels=label_temp;
h.XDisplayLabels=CustomXLabels;
h.YDisplayLabels=CustomYLabels;

end
```
