## Supplemental Code for "Neural circuit-selective, multiplexed pharmacological targeting of prefrontal cortex-projecting locus coeruleus neurons drives antinociception": ROI_QC_sheet.m

```
function [ROI_information_sheet]=ROI_QC_sheet(roi_n)
% roi_n: number of ROIs
% ROI_information_sheet: Main sheet for spike inference
%example code [ROI_information_sheet]=ROI_QC_sheet(20);
ROI_information_sheet.threshold=zeros(roi_n,2); 
ROI_information_sheet.quality=ones(roi_n,1); 
ROI_information_sheet.sm_rpf=zeros(roi_n,4); 
ROI_information_sheet.rph=zeros(roi_n,2); 
ROI_information_sheet.repeated_cell=zeros(roi_n,1); 
ROI_information_sheet.manual=zeros(roi_n,4); 
end
%threshold: MinProminence of prediction trace
%sm: Mean filter window for noisy raw trace
%rpf: MinPeakProminence for raw trace for noise filter;
%rph: MinPeakProminence for raw trace;
%quality: ROI quality
%repeated cell: found repeated ROIs
%manual: manual counting of AP numbers for specific ROIs (if necessary)
```
