## Supplemental Code for "Neural circuit-selective, multiplexed pharmacological targeting of prefrontal cortex-projecting locus coeruleus neurons drives antinociception": showing_traces.m

```
function showing_traces(traces,FPS,row)
% traces: ROI x T traces 
% FPS: frame rate
% row: number of trace per figure
% Example code: showing_traces(traces,20,9);

for i=1:ceil(size(traces,1)/row)
    figure('units','normalized','outerposition',[0 0 1 1]);
    for j=1:row
        subplot(row,1,j);
        if (i-1)*row+j<=size(traces,1)
            plot([1:size(traces,2)]/FPS,traces((i-1)*row+j,:),'k');
            title(append('ROI#',num2str((i-1)*row+j)),FontSize=12);
            xlim([1,size(traces,2)]/FPS);
        end
    end
    
    if i*row<=size(traces,1)
        sgtitle(append('ROI #',num2str((i-1)*row+1),' to #',num2str(i*row)),FontSize=20);
    else
        sgtitle(append('ROI #',num2str((i-1)*row+1),' to #',num2str(size(traces,1))),FontSize=20);
    end
end
end
```
