## Supplemental Code for "Neural circuit-selective, multiplexed pharmacological targeting of prefrontal cortex-projecting locus coeruleus neurons drives antinociception": slice_imaging_analysis_pipeline.m

```
%% Pipeline for spike inference
%% Data preprocess
% Use Fiji to unify constrast of imaging epoches & concatenate imaging epoches
% Use NoRMCorre algorithm to do non-rigid motion correction (rigid correction also works in acute slice too, code availability: https://github.com/flatironinstitute/NoRMCorre/tree/master)
% Trace smooth by floating baseline as below
[smooth_stack]=slice_imaging_smooth(motion_corrected_stack,floating_baseline,fps,1024); 
% Cut off first & last 50 data points for each epoch
% Export image file for ROI extraction as below
tiff_exporter(proceessed_stack,'example.tif',600); 
%% ROI extraction
% Use CNMF-E algorithm to extract temporal & spatial ROI info (default filename: results.A & results.C_raw, code availability: https://github.com/zhoupc/CNMF_E)
% Use 'demo_endoscope' script with common parameters as below
% gSig=4; gSiz=6; min_corr=0.6-0.8; min_pnr=3-5; min_pixel=gSig^2; bd=8; 
% May have MATLAB version issue (MATLAB R2021b used here)
% Use Fiji to calculate mean image from image stack then create images superimposing mean image of image stack & ROI locations as below
roi_contours(mean_image,results.A,128,128,0,[0.25,0.15]); 
%% Spike prediction
% Manually remove bursts artifictial events along traces
% Reformat & look through ROI temporal info for spike prediction model as below
[dF_traces]=prediction_material_convorter(results.C_raw); showing_traces(traces,fps,9);
% Use CASCADE algorithm to process spike prediction (in Python, default results 'spike_prob', code availability: https://github.com/HelmchenLabSoftware/Cascade)
%% Spike inference
% Create main data sheet 
[ROI_information_sheet]=ROI_QC_sheet(number_of_ROI); spike_sheet=[]; crteria_list=[0.8,0.6,0.4,0.2,0.1,0.05,0.01,0.005]; %Setup basic parameters & data sheets at the first time
% Identify peaks in both traces of raw calcium signal and spike likelihood for spike inference as below
[spike_sheet_temp]=spike_inference(dF_traces,spike_prob,crteria_list,[0,0],[0,0],[0,0],roi);
[spike_sheet]=spike_sheet_update(spike_sheet,spike_sheet_temp,roi);
%% Results update
% Update locations of predicted spike as below
[spikes_cascade]=spike_location_update(spike_prob,spike_sheet,ROI_information_sheet);
% Identify repeated ROIs then manual update repeated ROIs in main sheet
[events]=binary_spike_location(dF_traces,spikes_cascade);
[accept_ROI,repeat_roi,ROI_corrcoef_map]=ROI_cross_correlation(events,ROI_information_sheet,threshold_of_corr_coef);
% Update firing rates as below
[accept_ROI,pharmacology_results,spikes_bin]=pharmacology_profiling(spikes_cascade,ROI_information_sheet,3500,1750,fps,bin)
```
