## Supplemental Code for "Neural circuit-selective, multiplexed pharmacological targeting of prefrontal cortex-projecting locus coeruleus neurons drives antinociception": slice_imaging_smooth.m

```
function [smooth_data]=slice_imaging_smooth(data_in,baseline,fps,gap)
%data_in: d1*d2*t imaging data after motion correction 
%baseline: length of running baseline, usually 2 (second)
%fps: Imaging frame rate, usually 20
%gap: batch size for calculating, not affecting results, usually 1024
%Example code: [smooth_data]=slice_imaging_smooth(tiff_stack,2,20,1024);

baseline=baseline*fps;
Y=squeeze(reshape(data_in,[1,size(data_in,1)*size(data_in,2),size(data_in,3)]))';

for i=1:size(Y,2)
    if mod(i,gap)==1
        tic;
    end
    m=Y(:,i);
    head=Y(1:baseline,i);
    for j=1+baseline:size(Y,1)
        m(j,1)=mean(Y(j-baseline+1:j,i));
    end
    Y(1+baseline:size(Y,1),i)=Y(1+baseline:size(Y,1),i)-m(1+baseline:size(Y,1),1);
    Y(1:baseline,i)=head-mean(head);
    Y(:,i)=Y(:,i)+min(m(1+baseline:size(Y,1),1),[],'all');

    if mod(i,gap)==0
        formatSpec = 'Column # %.0f is processed sucessfully\b'; fprintf(formatSpec,i);
        formatSpec = ' out of %.0f pixel column.\n'; fprintf(formatSpec,size(Y,2));
        toc;
    end
end
Y=Y';
Y=Y-min(Y,[],'all');
smooth_data=reshape(Y,[size(data_in,1),size(data_in,2),size(data_in,3)]);

end
```
