## Supplemental Code for "Neural circuit-selective, multiplexed pharmacological targeting of prefrontal cortex-projecting locus coeruleus neurons drives antinociception": spike_inference.m

```
function [spike_sheet_temp]=spike_inference(data_in,spike_prob,crteria_list,sm,rpf,rph,roi)
%example code: [spike_sheet_temp]=spike_inference(dF_traces,spike_prob,crteria_list,[0,0],[0,0],[0,0],roi);
% spike_sheet_temp: temporal spike info for spike number & location update
% data_in: raw traces for prediction, default 'dF_traces'
% spike_prob: predicted traces of spike likelihood, default 'spike_prob'
% crteria_list: list of MinPeakProminence for traces of spike likelihood, default 'crteria_list'
% sm: Mean filter window for noisy raw trace
% rpf: MinPeakProminence for raw trace when doing noise filter
% rph: MinPeakProminence for raw trace
% roi: Interested ROI
%% Data setup
spike_sheet_temp.crteria_list=crteria_list; % Define threshold of for peak selection in predition trace
j=roi; % Define which ROI
a=spike_prob(j,51:size(spike_prob,2)/2); % a: Baseline prediction trace
b=spike_prob(j,1+size(spike_prob,2)/2:end-50); % b: Drug prediction trace
ra=data_in(j,51:size(spike_prob,2)/2); % ra: Baseline raw trace for further filter
rb=data_in(j,1+size(spike_prob,2)/2:end-50);  % rb: Drug raw trace for further filter
rra=ra;rrb=rb; %rra: Baseline raw trace; rra: Drug raw trace
%% Parameter setup
sm_pre=sm(1); sm_post=sm(2); % Mean filter window
rpf_pre=rpf(1); rpf_post=rpf(2); % Peak prominence criteria for raw trace filter
rph_pre=rph(1); rph_post=rph(2); % Peak prominence criteria for peak selection in raw trace
%% Noise filter
[y,x,w,p]=findpeaks(ra,'MinPeakProminence',0);
run=ra;
for i=1:size(x,2)
    if p(i)<rpf_pre || w(i)<=2 
        if x(i)>sm_pre+5 && x(i)<size(ra,2)-sm_pre-5 && sm_pre>0
            run(x(i)-1)=mean(ra(x(i)-sm_pre-1:x(i)+sm_pre-1));
            run(x(i))=mean(ra(x(i)-sm_pre:x(i)+sm_pre));
            run(x(i)+1)=mean(ra(x(i)-sm_pre+1:x(i)+sm_pre+1));
        end
    end
end
ra=run;
[y,x,w,p]=findpeaks(rb,'MinPeakProminence',0);    
run=rb;
for i=1:size(x,2)
    if p(i)<rpf_post || w(i)<=2
        if  x(i)>sm_post+5 && x(i)<size(ra,2)-sm_post-5 && sm_post>0
            run(1,x(i)-1)=mean(rb(x(i)-sm_post-1:x(i)+sm_post-1));
            run(1,x(i))=mean(rb(x(i)-sm_post:x(i)+sm_post));
            run(1,x(i)+1)=mean(rb(x(i)-sm_post+1:x(i)+sm_post+1));
        end
    end
end
rb=run;
%% Peak selection for baseline epoch
[y11,x11]=findpeaks(a,'MinPeakProminence',spike_sheet_temp.crteria_list(1));
runx=[];runy=[];
[ya,xa]=findpeaks(ra,'MinPeakProminence',rph_pre);
for k=1:size(xa,2)
    if length(find((x11-xa(k))>-5))-length(find((x11-xa(k))>10))~=0
        runx=cat(2,runx,xa(k));
        runy=cat(2,runy,ya(k));
    end
end
runxr=zeros(size(runx));runyr=zeros(size(runy));
for o=1:size(runx,2)
    xd=find(abs(x11-runx(o))==min(abs(x11-runx(o))));
    if size(xd,2)>1
        if length(find(runxr==x11(xd(1))))==0
            xd=xd(1);
        else
            xd=xd(2);
        end
    end
    runxr(o)=x11(xd);runyr(o)=y11(xd);
    x11(xd)=1;y11(xd)=1;
end
for p=1:size(runxr,2)-1
    if runxr(p)==runxr(p+1)
        runxr(p)=NaN;runyr(p)=NaN;
    end
end
runxr(isnan(runxr))=[];runyr(isnan(runyr))=[];runxr=sort(runxr);runyr=sort(runyr);
x11=runxr;y11=runyr;
[y12,x12]=findpeaks(a,'MinPeakProminence',spike_sheet_temp.crteria_list(2));
runx=[];runy=[];
[ya,xa]=findpeaks(ra,'MinPeakProminence',rph_pre);
for k=1:size(xa,2)
    if length(find((x12-xa(k))>-5))-length(find((x12-xa(k))>10))~=0
        runx=cat(2,runx,xa(k));
        runy=cat(2,runy,ya(k));
    end
end
runxr=zeros(size(runx));runyr=zeros(size(runy));
for o=1:size(runx,2)
    xd=find(abs(x12-runx(o))==min(abs(x12-runx(o))));
    if size(xd,2)>1
        if length(find(runxr==x12(xd(1))))==0
            xd=xd(1);
        else
            xd=xd(2);
        end
    end
    runxr(o)=x12(xd);runyr(o)=y12(xd);
    x12(xd)=1;y12(xd)=1;
end
for p=1:size(runxr,2)-1
    if runxr(p)==runxr(p+1)
        runxr(p)=NaN;runyr(p)=NaN;
    end
end
runxr(isnan(runxr))=[];runyr(isnan(runyr))=[];runxr=sort(runxr);runyr=sort(runyr);
x12=runxr;y12=runyr;
[y13,x13]=findpeaks(a,'MinPeakProminence',spike_sheet_temp.crteria_list(3));
runx=[];runy=[];
[ya,xa]=findpeaks(ra,'MinPeakProminence',rph_pre);
for k=1:size(xa,2)
    if length(find((x13-xa(k))>-5))-length(find((x13-xa(k))>10))~=0
        runx=cat(2,runx,xa(k));
        runy=cat(2,runy,ya(k));
    end
end
runxr=zeros(size(runx));runyr=zeros(size(runy));
for o=1:size(runx,2)
    xd=find(abs(x13-runx(o))==min(abs(x13-runx(o))));
    if size(xd,2)>1
        if length(find(runxr==x13(xd(1))))==0
            xd=xd(1);
        else
            xd=xd(2);
        end
    end
    runxr(o)=x13(xd);runyr(o)=y13(xd);
    x13(xd)=1;y13(xd)=1;
end
for p=1:size(runxr,2)-1
    if runxr(p)==runxr(p+1)
        runxr(p)=NaN;runyr(p)=NaN;
    end
end
runxr(isnan(runxr))=[];runyr(isnan(runyr))=[];runxr=sort(runxr);runyr=sort(runyr);
x13=runxr;y13=runyr;
[y14,x14]=findpeaks(a,'MinPeakProminence',spike_sheet_temp.crteria_list(4));
runx=[];runy=[];
[ya,xa]=findpeaks(ra,'MinPeakProminence',rph_pre);
for k=1:size(xa,2)
    if length(find((x14-xa(k))>-5))-length(find((x14-xa(k))>10))~=0
        runx=cat(2,runx,xa(k));
        runy=cat(2,runy,ya(k));
    end
end
runxr=zeros(size(runx));runyr=zeros(size(runy));
for o=1:size(runx,2)
    xd=find(abs(x14-runx(o))==min(abs(x14-runx(o))));
    if size(xd,2)>1
        if length(find(runxr==x14(xd(1))))==0
            xd=xd(1);
        else
            xd=xd(2);
        end
    end
    runxr(o)=x14(xd);runyr(o)=y14(xd);
    x14(xd)=1;y14(xd)=1;
end
for p=1:size(runxr,2)-1
    if runxr(p)==runxr(p+1)
        runxr(p)=NaN;runyr(p)=NaN;
    end
end
runxr(isnan(runxr))=[];runyr(isnan(runyr))=[];runxr=sort(runxr);runyr=sort(runyr);
x14=runxr;y14=runyr;
[y15,x15]=findpeaks(a,'MinPeakProminence',spike_sheet_temp.crteria_list(5));
runx=[];runy=[];
[ya,xa]=findpeaks(ra,'MinPeakProminence',rph_pre);
for k=1:size(xa,2)
    if length(find((x15-xa(k))>-5))-length(find((x15-xa(k))>10))~=0
        runx=cat(2,runx,xa(k));
        runy=cat(2,runy,ya(k));
    end
end
runxr=zeros(size(runx));runyr=zeros(size(runy));
for o=1:size(runx,2)
    xd=find(abs(x15-runx(o))==min(abs(x15-runx(o))));
    if size(xd,2)>1
        if length(find(runxr==x15(xd(1))))==0
            xd=xd(1);
        else
            xd=xd(2);
        end
    end
    runxr(o)=x15(xd);runyr(o)=y15(xd);
    x15(xd)=1;y15(xd)=1;
end
for p=1:size(runxr,2)-1
    if runxr(p)==runxr(p+1)
        runxr(p)=NaN;runyr(p)=NaN;
    end
end
runxr(isnan(runxr))=[];runyr(isnan(runyr))=[];runxr=sort(runxr);runyr=sort(runyr);
x15=runxr;y15=runyr;
[y16,x16]=findpeaks(a,'MinPeakProminence',spike_sheet_temp.crteria_list(6));
runx=[];runy=[];
[ya,xa]=findpeaks(ra,'MinPeakProminence',rph_pre);
for k=1:size(xa,2)
    if length(find((x16-xa(k))>-5))-length(find((x16-xa(k))>10))~=0
        runx=cat(2,runx,xa(k));
        runy=cat(2,runy,ya(k));
    end
end
runxr=zeros(size(runx));runyr=zeros(size(runy));
for o=1:size(runx,2)
    xd=find(abs(x16-runx(o))==min(abs(x16-runx(o))));
    if size(xd,2)>1
        if length(find(runxr==x16(xd(1))))==0
            xd=xd(1);
        else
            xd=xd(2);
        end
    end
    runxr(o)=x16(xd);runyr(o)=y16(xd);
    x16(xd)=1;y16(xd)=1;
end
for p=1:size(runxr,2)-1
    if runxr(p)==runxr(p+1)
        runxr(p)=NaN;runyr(p)=NaN;
    end
end
runxr(isnan(runxr))=[];runyr(isnan(runyr))=[];runxr=sort(runxr);runyr=sort(runyr);
x16=runxr;y16=runyr;
[y17,x17]=findpeaks(a,'MinPeakProminence',spike_sheet_temp.crteria_list(7));
runx=[];runy=[];
[ya,xa]=findpeaks(ra,'MinPeakProminence',rph_pre);
for k=1:size(xa,2)
    if length(find((x17-xa(k))>-5))-length(find((x17-xa(k))>10))~=0
        runx=cat(2,runx,xa(k));
        runy=cat(2,runy,ya(k));
    end
end
runxr=zeros(size(runx));runyr=zeros(size(runy));
for o=1:size(runx,2)
    xd=find(abs(x17-runx(o))==min(abs(x17-runx(o))));
    if size(xd,2)>1
        if length(find(runxr==x17(xd(1))))==0
            xd=xd(1);
        else
            xd=xd(2);
        end
    end
    runxr(o)=x17(xd);runyr(o)=y17(xd);
    x17(xd)=1;y17(xd)=1;
end
for p=1:size(runxr,2)-1
    if runxr(p)==runxr(p+1)
        runxr(p)=NaN;runyr(p)=NaN;
    end
end
runxr(isnan(runxr))=[];runyr(isnan(runyr))=[];runxr=sort(runxr);runyr=sort(runyr);
x17=runxr;y17=runyr;
[y18,x18]=findpeaks(a,'MinPeakProminence',spike_sheet_temp.crteria_list(8));
runx=[];runy=[];
[ya,xa]=findpeaks(ra,'MinPeakProminence',rph_pre);
for k=1:size(xa,2)
    if length(find((x18-xa(k))>-5))-length(find((x18-xa(k))>10))~=0
        runx=cat(2,runx,xa(k));
        runy=cat(2,runy,ya(k));
    end
end
runxr=zeros(size(runx));runyr=zeros(size(runy));
for o=1:size(runx,2)
    xd=find(abs(x18-runx(o))==min(abs(x18-runx(o))));
    if size(xd,2)>1
        if length(find(runxr==x18(xd(1))))==0
            xd=xd(1);
        else
            xd=xd(2);
        end
    end
    runxr(o)=x18(xd);runyr(o)=y18(xd);
    x18(xd)=1;y18(xd)=1;
end
for p=1:size(runxr,2)-1
    if runxr(p)==runxr(p+1)
        runxr(p)=NaN;runyr(p)=NaN;
    end
end
runxr(isnan(runxr))=[];runyr(isnan(runyr))=[];runxr=sort(runxr);runyr=sort(runyr);
x18=runxr;y18=runyr;
spike_sheet_temp.spike_n_pre(j,:)=[size(x11,2),size(x12,2),size(x13,2),size(x14,2),size(x15,2),size(x16,2),size(x17,2),size(x18,2)];

%% Peak selection for drug epoch
[y21,x21]=findpeaks(b,'MinPeakProminence',spike_sheet_temp.crteria_list(1));
runx=[];runy=[];
[yb,xb]=findpeaks(rb,'MinPeakProminence',rph_post);
for k=1:size(xb,2)
    if length(find((x21-xb(k))>-5))-length(find((x21-xb(k))>10))~=0
        runx=cat(2,runx,xb(k));
        runy=cat(2,runy,yb(k));
    end
end
runxr=zeros(size(runx));runyr=zeros(size(runy));
for o=1:size(runx,2)
    xd=find(abs(x21-runx(o))==min(abs(x21-runx(o))));
    if size(xd,2)>1
        if length(find(runxr==x21(xd(1))))==0
            xd=xd(1);
        else
            xd=xd(2);
        end
    end
    runxr(o)=x21(xd);runyr(o)=y21(xd);
    x21(xd)=1;y21(xd)=1;
end
for p=1:size(runxr,2)-1
    if runxr(p)==runxr(p+1)
        runxr(p)=NaN;runyr(p)=NaN;
    end
end
runxr(isnan(runxr))=[];runyr(isnan(runyr))=[];runxr=sort(runxr);runyr=sort(runyr);
x21=runxr;y21=runyr;
[y22,x22]=findpeaks(b,'MinPeakProminence',spike_sheet_temp.crteria_list(2));
runx=[];runy=[];
[yb,xb]=findpeaks(rb,'MinPeakProminence',rph_post);
for k=1:size(xb,2)
    if length(find((x22-xb(k))>-5))-length(find((x22-xb(k))>10))~=0
        runx=cat(2,runx,xb(k));
        runy=cat(2,runy,yb(k));
    end
end
runxr=zeros(size(runx));runyr=zeros(size(runy));
for o=1:size(runx,2)
    xd=find(abs(x22-runx(o))==min(abs(x22-runx(o))));
    if size(xd,2)>1
        if length(find(runxr==x22(xd(1))))==0
            xd=xd(1);
        else
            xd=xd(2);
        end
    end
    runxr(o)=x22(xd);runyr(o)=y22(xd);
    x22(xd)=1;y22(xd)=1;
end
for p=1:size(runxr,2)-1
    if runxr(p)==runxr(p+1)
        runxr(p)=NaN;runyr(p)=NaN;
    end
end
runxr(isnan(runxr))=[];runyr(isnan(runyr))=[];runxr=sort(runxr);runyr=sort(runyr);
x22=runxr;y22=runyr;
[y23,x23]=findpeaks(b,'MinPeakProminence',spike_sheet_temp.crteria_list(3));
runx=[];runy=[];
[yb,xb]=findpeaks(rb,'MinPeakProminence',rph_post);
for k=1:size(xb,2)
    if length(find((x23-xb(k))>-5))-length(find((x23-xb(k))>10))~=0
        runx=cat(2,runx,xb(k));
        runy=cat(2,runy,yb(k));
    end
end
runxr=zeros(size(runx));runyr=zeros(size(runy));
for o=1:size(runx,2)
    xd=find(abs(x23-runx(o))==min(abs(x23-runx(o))));
    if size(xd,2)>1
        if length(find(runxr==x23(xd(1))))==0
            xd=xd(1);
        else
            xd=xd(2);
        end
    end
    runxr(o)=x23(xd);runyr(o)=y23(xd);
    x23(xd)=1;y23(xd)=1;
end
for p=1:size(runxr,2)-1
    if runxr(p)==runxr(p+1)
        runxr(p)=NaN;runyr(p)=NaN;
    end
end
runxr(isnan(runxr))=[];runyr(isnan(runyr))=[];runxr=sort(runxr);runyr=sort(runyr);
x23=runxr;y23=runyr;
[y24,x24]=findpeaks(b,'MinPeakProminence',spike_sheet_temp.crteria_list(4));
runx=[];runy=[];
[yb,xb]=findpeaks(rb,'MinPeakProminence',rph_post);
for k=1:size(xb,2)
    if length(find((x24-xb(k))>-5))-length(find((x24-xb(k))>10))~=0
        runx=cat(2,runx,xb(k));
        runy=cat(2,runy,yb(k));
    end
end
runxr=zeros(size(runx));runyr=zeros(size(runy));
for o=1:size(runx,2)
    xd=find(abs(x24-runx(o))==min(abs(x24-runx(o))));
    if size(xd,2)>1
        if length(find(runxr==x24(xd(1))))==0
            xd=xd(1);
        else
            xd=xd(2);
        end
    end
    runxr(o)=x24(xd);runyr(o)=y24(xd);
    x24(xd)=1;y24(xd)=1;
end
for p=1:size(runxr,2)-1
    if runxr(p)==runxr(p+1)
        runxr(p)=NaN;runyr(p)=NaN;
    end
end
runxr(isnan(runxr))=[];runyr(isnan(runyr))=[];runxr=sort(runxr);runyr=sort(runyr);
x24=runxr;y24=runyr;
[y25,x25]=findpeaks(b,'MinPeakProminence',spike_sheet_temp.crteria_list(5));
runx=[];runy=[];
[yb,xb]=findpeaks(rb,'MinPeakProminence',rph_post);
for k=1:size(xb,2)
    if length(find((x25-xb(k))>-5))-length(find((x25-xb(k))>10))~=0
        runx=cat(2,runx,xb(k));
        runy=cat(2,runy,yb(k));
    end
end
runxr=zeros(size(runx));runyr=zeros(size(runy));
for o=1:size(runx,2)
    xd=find(abs(x25-runx(o))==min(abs(x25-runx(o))));
    if size(xd,2)>1
        if length(find(runxr==x25(xd(1))))==0
            xd=xd(1);
        else
            xd=xd(2);
        end
    end
    runxr(o)=x25(xd);runyr(o)=y25(xd);
    x25(xd)=1;y25(xd)=1;
end
for p=1:size(runxr,2)-1
    if runxr(p)==runxr(p+1)
        runxr(p)=NaN;runyr(p)=NaN;
    end
end
runxr(isnan(runxr))=[];runyr(isnan(runyr))=[];runxr=sort(runxr);runyr=sort(runyr);
x25=runxr;y25=runyr;
[y26,x26]=findpeaks(b,'MinPeakProminence',spike_sheet_temp.crteria_list(6));
runx=[];runy=[];
[yb,xb]=findpeaks(rb,'MinPeakProminence',rph_post);
for k=1:size(xb,2)
    if length(find((x26-xb(k))>-5))-length(find((x26-xb(k))>10))~=0
        runx=cat(2,runx,xb(k));
        runy=cat(2,runy,yb(k));
    end
end
runxr=zeros(size(runx));runyr=zeros(size(runy));
for o=1:size(runx,2)
    xd=find(abs(x26-runx(o))==min(abs(x26-runx(o))));
    if size(xd,2)>1
        if length(find(runxr==x26(xd(1))))==0
            xd=xd(1);
        else
            xd=xd(2);
        end
    end
    runxr(o)=x26(xd);runyr(o)=y26(xd);
    x26(xd)=1;y26(xd)=1;
end
for p=1:size(runxr,2)-1
    if runxr(p)==runxr(p+1)
        runxr(p)=NaN;runyr(p)=NaN;
    end
end
runxr(isnan(runxr))=[];runyr(isnan(runyr))=[];runxr=sort(runxr);runyr=sort(runyr);
x26=runxr;y26=runyr;
[y27,x27]=findpeaks(b,'MinPeakProminence',spike_sheet_temp.crteria_list(7));
runx=[];runy=[];
[yb,xb]=findpeaks(rb,'MinPeakProminence',rph_post);
for k=1:size(xb,2)
    if length(find((x27-xb(k))>-5))-length(find((x27-xb(k))>10))~=0
        runx=cat(2,runx,xb(k));
        runy=cat(2,runy,yb(k));
    end
end
runxr=zeros(size(runx));runyr=zeros(size(runy));
for o=1:size(runx,2)
    xd=find(abs(x27-runx(o))==min(abs(x27-runx(o))));
    if size(xd,2)>1
        if length(find(runxr==x27(xd(1))))==0
            xd=xd(1);
        else
            xd=xd(2);
        end
    end
    runxr(o)=x27(xd);runyr(o)=y27(xd);
    x27(xd)=1;y27(xd)=1;
end
for p=1:size(runxr,2)-1
    if runxr(p)==runxr(p+1)
        runxr(p)=NaN;runyr(p)=NaN;
    end
end
runxr(isnan(runxr))=[];runyr(isnan(runyr))=[];runxr=sort(runxr);runyr=sort(runyr);
x27=runxr;y27=runyr;
[y28,x28]=findpeaks(b,'MinPeakProminence',spike_sheet_temp.crteria_list(8));
runx=[];runy=[];
[yb,xb]=findpeaks(rb,'MinPeakProminence',rph_post);
for k=1:size(xb,2)
    if length(find((x28-xb(k))>-5))-length(find((x28-xb(k))>10))~=0
        runx=cat(2,runx,xb(k));
        runy=cat(2,runy,yb(k));
    end
end
runxr=zeros(size(runx));runyr=zeros(size(runy));
for o=1:size(runx,2)
    xd=find(abs(x28-runx(o))==min(abs(x28-runx(o))));
    if size(xd,2)>1
        if length(find(runxr==x28(xd(1))))==0
            xd=xd(1);
        else
            xd=xd(2);
        end
    end
    runxr(o)=x28(xd);runyr(o)=y28(xd);
    x28(xd)=1;y28(xd)=1;
end
for p=1:size(runxr,2)-1
    if runxr(p)==runxr(p+1)
        runxr(p)=NaN;runyr(p)=NaN;
    end
end
runxr(isnan(runxr))=[];runyr(isnan(runyr))=[];runxr=sort(runxr);runyr=sort(runyr);
x28=runxr;y28=runyr;
%% Create files for AP location update
spike_sheet_temp.spike_n_post(j,:)=[size(x21,2),size(x22,2),size(x23,2),size(x24,2),size(x25,2),size(x26,2),size(x27,2),size(x28,2)];
x21=x21+size(spike_prob,2)/2-50;
x22=x22+size(spike_prob,2)/2-50;
x23=x23+size(spike_prob,2)/2-50;
x24=x24+size(spike_prob,2)/2-50;
x25=x25+size(spike_prob,2)/2-50;
x26=x26+size(spike_prob,2)/2-50;
x27=x27+size(spike_prob,2)/2-50;
x28=x28+size(spike_prob,2)/2-50;
spike_sheet_temp.deconvolution{j,1}=[x11 x21];
spike_sheet_temp.deconvolution{j,2}=[x12 x22];
spike_sheet_temp.deconvolution{j,3}=[x13 x23];
spike_sheet_temp.deconvolution{j,4}=[x14 x24];
spike_sheet_temp.deconvolution{j,5}=[x15 x25];
spike_sheet_temp.deconvolution{j,6}=[x16 x26];
spike_sheet_temp.deconvolution{j,7}=[x17 x27];
spike_sheet_temp.deconvolution{j,8}=[x18 x28];
%% Prepare raw/filter traces for ploting
c=[ra rb];
c(find(c<0))=0;
c=0-c;
c=c+1.5;
d=[rra rrb];
d(find(d<0))=0;
d=0-d;
d=d+1.5;
%% Plot-part1
figure('units','normalized','outerposition',[0 0 1 1]);
subplot 411;
plot(d,'m');
hold on
plot(c,'b');
plot([a b],'k');
xbar=zeros(1,size(c,2));
run=[x11 x21];
for n=1:size(run,2)
    xbar(run(n))=1.5;
end
bar(xbar,'r',LineStyle='none');
set(gcf,'color','w');
ax = gca;
ax.FontSize=8;
title(append('Spike Prediction, Local Prominence > ',num2str(crteria_list(1))),'fontsize', 12);
hold off

subplot 412;
plot(d,'m');
hold on
plot(c,'b');
plot([a b],'k');
xbar=zeros(1,size(c,2));
run=[x12 x22];
for n=1:size(run,2)
    xbar(run(n))=1.5;
end
bar(xbar,'r',LineStyle='none');
set(gcf,'color','w');
ax = gca;
ax.FontSize=8;
title(append('Spike Prediction, Local Prominence > ',num2str(crteria_list(2))),'fontsize', 12);
hold off

subplot 413;
plot(d,'m');
hold on
plot(c,'b');
plot([a b],'k');
xbar=zeros(1,size(c,2));
run=[x13 x23];
for n=1:size(run,2)
    xbar(run(n))=1.5;
end
bar(xbar,'r',LineStyle='none');
set(gcf,'color','w');
ax = gca;
ax.FontSize=8;
title(append('Spike Prediction, Local Prominence > ',num2str(crteria_list(3))),'fontsize', 12);
hold off

subplot 414;
plot(d,'m');
hold on
plot(c,'b');
plot([a b],'k');
xbar=zeros(1,size(c,2));
run=[x14 x24];
for n=1:size(run,2)
    xbar(run(n))=1.5;
end
bar(xbar,'r',LineStyle='none');
set(gcf,'color','w');
ax = gca;
ax.FontSize=8;
title(append('Spike Prediction, Local Prominence > ',num2str(crteria_list(4))),'fontsize', 12);
hold off

sgtitle(append('ROI #',num2str(roi)),'fontsize', 16)
%% Plot-part2
figure('units','normalized','outerposition',[0 0 1 1]);
subplot 411;
plot(d,'m');
hold on
plot(c,'b');
plot([a b],'k');
xbar=zeros(1,size(c,2));
run=[x15 x25];
for n=1:size(run,2)
    xbar(run(n))=1.5;
end
bar(xbar,'r',LineStyle='none');
set(gcf,'color','w');
ax = gca;
ax.FontSize=8;
title(append('Spike Prediction, Local Prominence > ',num2str(crteria_list(5))),'fontsize', 12);
hold off

subplot 412;
plot(d,'m');
hold on
plot(c,'b');
plot([a b],'k');
xbar=zeros(1,size(c,2));
run=[x16 x26];
for n=1:size(run,2)
    xbar(run(n))=1.5;
end
bar(xbar,'r',LineStyle='none');
set(gcf,'color','w');
ax = gca;
ax.FontSize=8;
title(append('Spike Prediction, Local Prominence > ',num2str(crteria_list(6))),'fontsize', 12);
hold off

subplot 413;
plot(d,'m');
hold on
plot(c,'b');
plot([a b],'k');
xbar=zeros(1,size(c,2));
run=[x17 x27];
for n=1:size(run,2)
    xbar(run(n))=1.5;
end
bar(xbar,'r',LineStyle='none');
set(gcf,'color','w');
ax = gca;
ax.FontSize=8;
title(append('Spike Prediction, Local Prominence > ',num2str(crteria_list(7))),'fontsize', 12);
hold off

subplot 414;
plot(d,'m');
hold on
plot(c,'b');
plot([a b],'k');
xbar=zeros(1,size(c,2));
run=[x18 x28];
for n=1:size(run,2)
    xbar(run(n))=1.5;
end
bar(xbar,'r',LineStyle='none');
set(gcf,'color','w');
ax = gca;
ax.FontSize=8;
title(append('Spike Prediction, Local Prominence > ',num2str(crteria_list(8))),'fontsize', 12);
hold off

sgtitle(append('ROI #',num2str(roi)),'fontsize', 16)
end
```
