## Supplemental Code for "Neural circuit-selective, multiplexed pharmacological targeting of prefrontal cortex-projecting locus coeruleus neurons drives antinociception": spike_location_update.m

```
function [spikes_cascade]=spike_location_update(spike_prob,spike_sheet,ROI_information)
% Example code: [spikes_cascade]=spike_location_update(spike_prob,spike_sheet,ROI_information_sheet);
% spikes_cascade: spike locations as spike inference
% spike_prob: Spike likelihood traces
% spike_sheet: Temporal info for spike deconvolution using all peak criteria
% ROI_information: Main sheet of spike inference, default 'ROI_information_sheet'
spikes_cascade={};
for i=1:size(spike_prob,1)
    threshold=ROI_information.threshold(i,1:2);
    if ROI_information.threshold(i,1)==0
        spikes_temp01=[];
    else
        spikes_temp01=spike_sheet.deconvolution{i,threshold(1)}';
        spikes_temp01(find(spikes_temp01>size(spike_prob,2)/2-50))=[];
    end
    if ROI_information.threshold(i,2)==0
        spikes_temp02=[];
    else
        spikes_temp02=spike_sheet.deconvolution{i,threshold(2)}';
        spikes_temp02(find(spikes_temp02<=size(spike_prob,2)/2-50))=[];
    end
    spikes_cascade{i}=cat(1,spikes_temp01,spikes_temp02);
end
end
```
