## Supplemental Code for "Neural circuit-selective, multiplexed pharmacological targeting of prefrontal cortex-projecting locus coeruleus neurons drives antinociception": spike_sheet_update.m

```
function [spike_sheet]=spike_sheet_update(spike_sheet,spike_sheet_temp,roi)
% spike_sheet: File that is updating
% spike_sheet_temp: Temporal file of spike inference, default 'spike_sheet_temp'
% roi: Interested ROI
% Example code: [spike_sheet]=spike_sheet_update(spike_sheet,spike_sheet_temp,1);

j=roi;
spike_sheet.crteria_list=spike_sheet_temp.crteria_list;
for k=1:size(spike_sheet_temp.crteria_list,2)
spike_sheet.deconvolution{j,k}=spike_sheet_temp.deconvolution{j,k};
end
spike_sheet.spike_n_pre(j,:)=spike_sheet_temp.spike_n_pre(j,:);
spike_sheet.spike_n_post(j,:)=spike_sheet_temp.spike_n_post(j,:);
end
```
