## Supplemental Code for "Neural circuit-selective, multiplexed pharmacological targeting of prefrontal cortex-projecting locus coeruleus neurons drives antinociception": tiff_exporter.m

```
function tiff_exporter(data,filename,gap)
%data: d1*d2*t stack matrix
%filename: name for output file
%gap: gap of progress indication, not affecting results 
%code example = tiff_exporter(proceessed_stack,'example.tif',600); 
M=data/max(data,[],'all');
t=Tiff(filename,'w');
tagstruct.ImageLength=size(M,1);
tagstruct.ImageWidth=size(M,2);
tagstruct.Photometric=Tiff.Photometric.MinIsBlack;
tagstruct.PlanarConfiguration=Tiff.PlanarConfiguration.Chunky;
tagstruct.BitsPerSample = 16;
tagstruct.SamplesPerPixel = 1;
tagstruct.Software = 'MATLAB';

for i=1:size(M,3)
    if mod(i,gap)==1
        tic;
    end
    setTag(t,tagstruct);
    write(t,squeeze(im2uint16(M(:,:,i))));
    writeDirectory(t);
    if mod(i,gap)==0
        formatSpec = 'Stack # %.0f is processed sucessfully\b'; fprintf(formatSpec,i);
        formatSpec = ' out of %.0f stack.\n'; fprintf(formatSpec,size(M,3));
        toc;
    end
end
close(t)
end
```
